# Structural Basis and Designing of Peptide Vaccine using PE-PGRS Family Protein of *Mycobacterium ulcerans* – An Integrated Vaccinomics Approach

**DOI:** 10.1101/795146

**Authors:** Zulkar Nain, Mohammad Minnatul Karim, Monokesh Kumer Sen, Utpal Kumar Adhikari

**Affiliations:** Department of Biotechnology and Genetic Engineering, Faculty of Biological Sciences, Islamic University, Kushtia-7003, Bangladesh; School of Medicine, Western Sydney University, Locked Bag 1797, Penrith, NSW-2751, Australia

**Keywords:** Buruli ulcer, *Mycobacterium ulcerans*, PE-PGRS protein, Multi-epitope vaccine, Skin disease, Immunoinformatics, Bioinformatics

## Abstract

Buruli ulcer is an emerging-necrotizing skin infection, responsible for permanent deformity if untreated, caused by the pathogen *Mycobacterium ulcerans* (*M. ulcerans*). Despite this debilitating condition, no specific disease-modifying therapeutics or vaccination is available. Therefore, we aimed to design an effective multi-epitope vaccine against *M. ulcerans* through an integrated vaccinomics approach. Briefly, the highest antigenic PE-PGRS protein was selected from which the promiscuous T- and B-cell epitopes were predicted. After rigorous assessment, 15 promising CTL, HTL and LBL epitopes were selected. The identified T-cell epitopes showed marked interactions towards the HLA binding alleles and provided 99.8% world population coverage. Consequently, a vaccine chimera was designed by connecting these epitopes with suitable linkers and adjuvant (LprG). The vaccine construct was antigenic and immunogenic as well as non-allergenic; hence, subjected to homology modelling. The molecular docking and dynamic simulation revealed strong and stable binding affinity between the vaccine and TLR2 receptor. The binding energy (ΔG) and dissociation constant (K_d_) were −15.3 kcal/mol and 5.9×10^−12^ M, respectively. Further, disulfide engineering was applied to improve vaccine’ stability and higher expression in *Escherichia coli* K12 system was ensured by codon optimization and cloning *in silico*. The computer-simulated immune responses were characterized by higher levels of IgM and IgG antibodies, helper T-cells with increased IFN-γ production, and macrophage activity crucial for immunity against *M. ulcerans*. Therefore, our data suggest that, if the designed vaccine is validated experimentally, it will prevent Buruli ulcer by generating robust immune response against *M. ulcerans*.

## 1. Introduction

Buruli ulcer is a neglected tropical disease of the skin characterized by chronic skin lesions and tissue necrosis leading to permanent cosmetic deformity and functional disability affecting thousands of people worldwide (Pluschke and Röltgen 2015; Simpson et al. 2019; Barksby 2019). Despite being endemic to West and Central Africa (Johnson 2019), Buruli ulcer has been reported from more than 34 countries worldwide including countries from South America and Western Pacific regions (Pluschke and Röltgen 2015; Singh et al. 2019; Simpson et al. 2019). About 80% of cases were from West African countries *i.e.*, Côte d’Ivoire, Ghana, Benin and Cameroon (Mitra AK 2017). However, there is no consensus on the current distribution of this mycobacterial disease, except for 12 countries which constitute ~34,890 reported cases worldwide from 2007 to 2016 (Simpson et al. 2019). Buruli ulcer has become the third most common mycobacterial disease after tuberculosis and leprosy (Phillips et al. 2015; Huygen et al. 2009). Although it may affect everyone irrespective of ages but the risk is the highest in the children (~15 years old) (Huygen et al. 2009) and aged people (>50 years old) (N’krumah et al. 2016).

*Mycobacterium ulcerans* (*M. ulcerans*) is the causative agent of Buruli ulcer (N’krumah et al. 2016; Singh et al. 2019; Simpson et al. 2019). The pathogen *M. ulcerans* has evolved from the fish pathogen *M. marinum* which rarely causes skin lesions in humans (Yip et al. 2007; Petrini 2006). The emergence of *M. ulcerans* has been associated by the acquisition of pMUM plasmid that encodes the gene for the production of mycolactone (Johnson 2019; George et al. 1999), a polyketide-derived diffusible exotoxin, that has cytotoxic and immunosuppressive properties responsible for chronic skin lesions (Adusumilli et al. 2005; Walsh et al. 2005; Hong et al. 2008) and destroying the immune cells before reaching the infection site (Pluschke and Röltgen 2015). This exotoxin (mycolactone) is the main virulence factor of *M. ulcerans* (George et al. 1999; Barksby 2019). However, the occurrence of Buruli ulcer is still perplexing due to the incomplete knowledge about the reservoirs and transmission pathways of *M. ulcerans* (Barksby 2019). A few lines of evidence has described that disease transmission is associated with contaminated water (Huygen et al. 2009; Singh et al. 2019; Sears and Hay 2015). Recent studies suggest that *M. ulcerans* uses different transmission vehicles in different geographic areas (Merritt et al. 2010). For example, aquatic insects in Benin contain *M. ulcerans* DNA which indicated their possible involvement in the transmission (Marsollier et al. 2002). Moreover, mosquitoes have been suspected of playing a detrimental role in bacterial transmission in Australia (Quek et al. 2007; Singh et al. 2019).

Buruli ulcer has become a major public health concern due to the lack of specific treatment and preventive measures (Simpson et al. 2019; Johnson 2019; Singh et al. 2019). Despite the effectiveness of antibiotics against Buruli ulcer, failures of treatment are common and the pathogen is prone to develop resistance (Phillips et al. 2015). In addition, there is no effective vaccination available for preventing Buruli ulcer (Phillips et al. 2015). There are, however, considerable evidences that supported the feasibility of vaccine preparation against *M. ulcerans* (Huygen et al. 2009). To date, several attempts have been made to develop a potent vaccine against *M. ulcerans* but were resulted in a limited success (Tanghe et al. 2008; Hart, Hale, and Lee 2016; Coutanceau et al. 2006; Ravenel 1928). The only vaccine that is widely used to control Buruli ulcer is a live attenuated *Mycobacterium bovis* vaccine known as Bacillus Calmette-Guérin (BCG) vaccine. However, effectiveness of BCG vaccine against Buruli ulcer is found contradictory to some extent (Phillips et al. 2015; Ravenel 1928). In a recent study, for instance, no significant evidence was found in favor of BCG vaccine’s protective effect against mild to severe form of Buruli ulcer (Phillips et al. 2015). Besides, there are also possible side-effects with the use of killed or live attenuated bacterial vaccines (Saadi, Karkhah, and Nouri 2017). For example, mycolactone-deficient attenuated *M. ulcerans* strain 5114 provides a short-term protection against infected mice (Fraga et al. 2012). Nevertheless, this attenuated strain may still retain factors which can hinder effective memory cell development (Hart, Hale, and Lee 2016). Furthermore, DNA vaccine encoding immunodominant Ag85A protein of *M. ulcerans* showed protection similar to BCG vaccination (Tanghe et al. 2008; Hart, Hale, and Lee 2016). These evidences are suggesting that the subunit vaccine could be an alternative in terms of safety, efficacy and specificity (Saadi, Karkhah, and Nouri 2017; Yin et al. 2016).

The subunit vaccine contains the fragment(s) of antigenic proteins that can mimic the presence of the natural pathogen and generate immune response against the target pathogen (Saadi, Karkhah, and Nouri 2017). In 1985, the first epitope-based subunit vaccine was developed against cholera in *Escherichia coli* (Jacob et al. 1985), while many are still under development such as the vaccine for malaria, swine flue, influenza, and anthrax (Li et al. 2014). Besides, some multi-epitope vaccines have even entered phase I clinical trial. For instance, survivin-derived multi-epitope cancer vaccine (EMD640744) used in patients with advanced tumors (L. Zhang 2018; Lennerz et al. 2014). Interestingly, multiple-component peptide vaccine are able to generate more protective immune response than the single component vaccine (Saadi, Karkhah, and Nouri 2017). Recently, *in silico* designing of multi-epitope vaccine against viruses, bacteria and parasites have become a commonplace due to the ease of access to bioinformatics tools and servers (Adhikari and Rahman 2017; A. Ali et al. 2019; M. Ali et al. 2017; Ikram et al. 2018; Mirza et al. 2016; Yasmin and Nabi 2016). Conceptually, multi-epitope vaccines have some advantages over classical (*i.e.*, live and attenuated) and single-epitope vaccines (Lin et al. 2016; Lu et al. 2017; Saadi, Karkhah, and Nouri 2017). For example, it can be manipulated in a variety of ways such as combining T- and B-cell epitopes from same or different antigen(s) derived from same source. Besides, unwanted component that produces toxic or allergenic reactions can be eliminated. Furthermore, adjuvant can be added to enhance the immunogenic potency (Saadi, Karkhah, and Nouri 2017). Therefore, it can be hypothesised that a well-designed multi-epitope vaccine with such advantages could be a potent prophylactic against *M. ulcerans*.

In this study, *M. ulcerans* proteome was explored to determine the highest antigenic protein(s) followed by the prediction of different T- and B-cell epitopes with their corresponding major histocompatibility complex (MHC) alleles. These epitopes were evaluated for their immunological profile. Finally, a multi-epitope vaccine was designed using the most potent epitopes with appropriate adjuvant and linkers. The primary sequence of the vaccine construct was used for the immunogenic and physiochemical profiling followed by homology modelling. The crude three-dimensional (3D) structure was then refined and subjected to disulfide engineering for stability enhancement. The binding interaction and stability of the vaccine-receptor complex were analyzed by molecular docking and dynamic simulation, respectively. Moreover, immune responses induced by the vaccine antigen were simulated to investigate the real-life potency. Finally, the vaccine codon was optimized for *E. coli* system and in silico cloning was performed. The computational workflow of the study is graphically illustrated in Fig. 1.

**Fig. 1:**
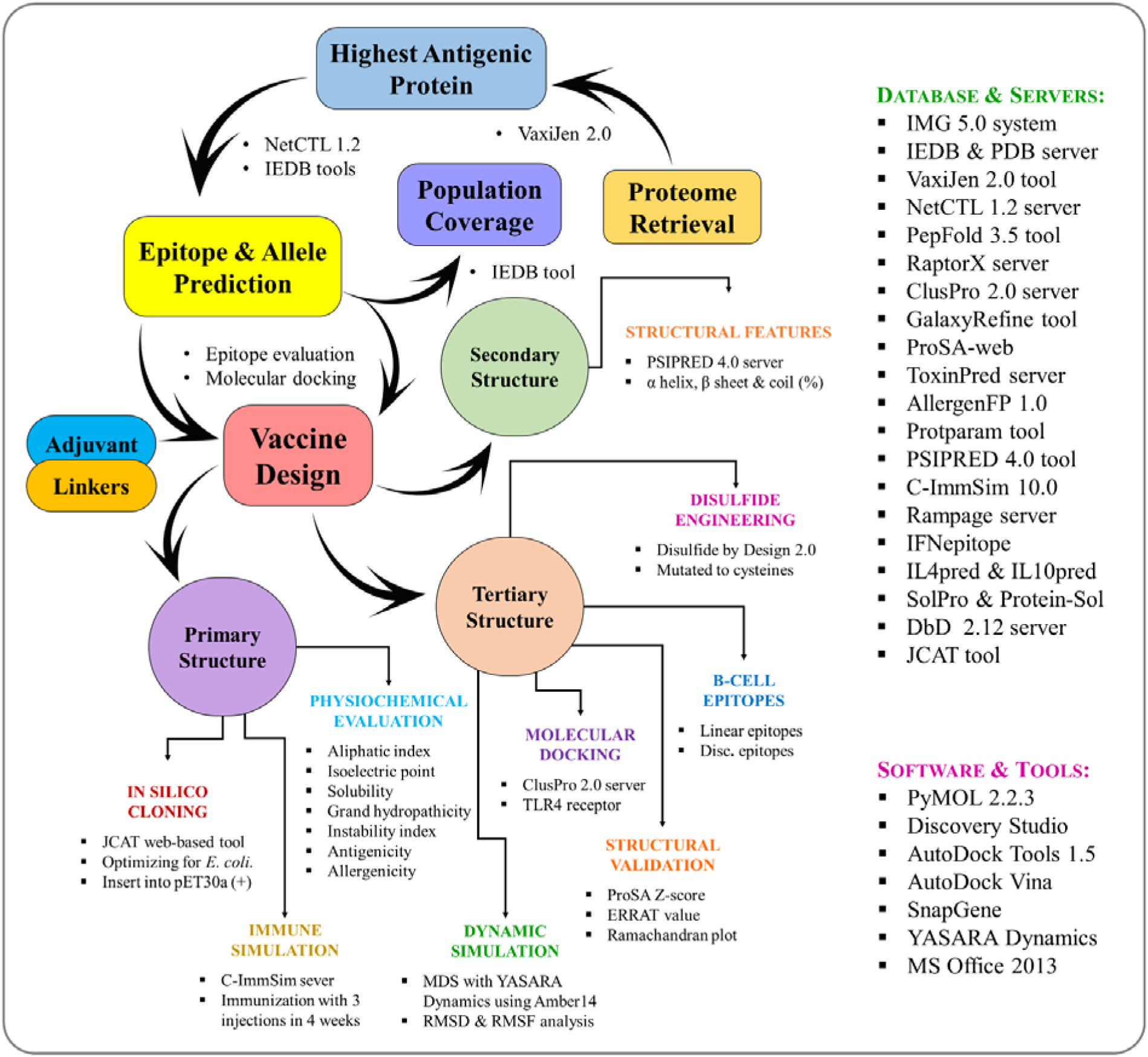
The overall experimental workflow used to develop a multi-epitope vaccine against *M. ulcerans*.

## 2. Materials and Methods

### 2.1. Immunoinformatic study of the antigen

#### 2.1.1. Proteome retrieval and prediction of antigenicity and transmembrane helix

The whole proteome of *M. ulcerans* Agy99 was collected from Integrated Microbial Genomes & Microbiomes (IMG/M) v5.0 system (Chen et al. 2019). Fed by the national center for biotechnology information (NCBI) database, this system integrates the draft and complete genomes of Archaea, Bacteria and Eukaryota which can be retrieved with or without protein homolog in any target species (Markowitz et al. 2012). Vaccine candidates should not have homology with human proteins to avoid autoimmune response (Monterrubio-López, González-Y-Merchand, and Ribas-Aparicio 2015). Therefore, human protein homologs were avoided during the proteome retrieval. Since for an effective peptide vaccine the target protein should be able to induce an immune response, the antigenicity of all protein sequences included in the proteome was predicted using the VaxiJen v2.0 server (Doytchinova and Flower 2007) with 0.5 threshold. This server uses auto cross-covariance transformation method to maintain 70-89% accurate prediction (Doytchinova and Flower 2007). Proteins with transmembrane helices are often created difficulty in purification (Monterrubio-López, González-Y-Merchand, and Ribas-Aparicio 2015); hence, they are not desired in target protein selection. The top-ten antigenic proteins were further subjected to transmembrane (TM) helix prediction with TMHMM v2.0 server (Krogh et al. 2001).

#### 2.1.2. Prediction and assessment of cytotoxic T-lymphocyte (CTL) epitopes

Cytotoxic T-cells play key role in specific antigen recognition that made CTL epitopes essential for coherent vaccine design (Chaudhri et al. 2009). Therefore, the CTL epitopes present in the selected protein were determined using the NetCTL v1.2 server with 0.5 threshold (Larsen et al. 2007). This server predicts CTL epitopes (9-mer) based on the MHC-I binding peptides, C-terminal cleavage, and TAP transport efficiency through artificial neural networks and weight matrix (Larsen et al. 2007). The IEDB MHC-I binding tool was used to anticipate the MHC-I binding alleles for each CTL epitope based on the CONSENSUS method (Moutaftsi et al. 2006). In this study, we considered percentile rank score ≤2; since the lower the rank, the higher the affinity (Moutaftsi et al. 2006). Vaccine components should be antigenic and immunogenic; simultaneously, it should be devoid of allergic and toxic reaction. So, each CTL epitope was evaluated for their individual antigenic, immunogenic, allergenic and toxic profiles. Therefore, the immunogenicity and antigenicity were evaluated with IEDB MHC-I immunogenicity tool (Calis et al. 2013) and VaxiJen v2.0 server (Doytchinova and Flower 2007), respectively. Furthermore, the AllergenFP v1.0 server (Dimitrov et al. 2014)was used for allergenicity and ToxinPred (Gupta et al. 2013) for toxicity prediction. Herein, the AllergenFP v1.0 applies novel alignment-free descriptor-based fingerprint approach for ensuring 88.9% prediction accuracy (Dimitrov et al. 2014) while ToxinPred works based on different peptide properties using machine learning techniques accompanied with quantitative matrix (Gupta et al. 2013).

#### 2.1.3. Prediction and evaluation of helper T-lymphocyte (HTL) epitopes

Being the most important cells in adaptive immunity, helper T-cells influence B-cells to secrete antibodies, macrophages to phagocyte pathogens, and cytotoxic T-cells to eliminate targeted parasitized cells (Zhu and Paul 2008), that are important in cell-mediated immunity against *M. ulcerans* (Fraga et al. 2011). Therefore, vaccine construct should contain HTL epitopes from rational perspective. Consequently, the selected protein was submitted to IEDB MHC-II binding tool (Wang et al. 2010) for the prediction of 15-mer HTL epitopes along with their respective binding alleles using the CONSENSUS method (Wang et al. 2008) The percentile rank threshold ≤2 was considered. Helper T-cells produce different cytokines such as interferon-gamma (IFN-γ), interleukin-4 (IL-4) and IL-10 resulting in the activation of cytotoxic T-cells and other immune cells such as macrophages (Luckheeram et al. 2012). Therefore, cytokine-inducing HTL epitopes are crucial for vaccine development. Consequently, IFNepitope server was applied to predict IFN-γ producing HTL epitopes with Motif and SVM based hybrid method and IFN-γ versus Non-IFN-γ model (Dhanda, Vir, and Raghava 2013). On the other hand, IL-4 and IL-10 inducing properties were predicted using IL4pred (Dhanda et al. 2013) and IL10pred (Nagpal et al. 2017) servers, respectively. The IL4pred and IL10pred operations were carried out based on SVM method with threshold value 0.2 and −0.3, respectively.

#### 2.1.4. Prediction and assessment of linear B-lymphocyte (LBL) epitopes

A surface receptor of B-cell recognizes B-cell epitopes, resulting the generation of antigen-specific immunoglobulins (Cooper 2015). Therefore, designing a vaccine consisting of B-cell epitopes can play an essential role in adaptive immunity. There are two types of B-cell epitopes, namely, linear and conformational epitopes (J. Zhang et al. 2014). In vaccine construction, only linear B-cell epitopes can be incorporated into the multi-peptide sequence (Oscherwitz 2016; Dormitzer, Ulmer, and Rappuoli 2008), hence, predicted using iBCE-EL server (Manavalan et al. 2018). This server predict 12-mer LBL epitopes by default using a novel ensemble learning framework consisting of two independent predictors, namely, extremely randomized tree and gradient boosting classifiers (Manavalan et al. 2018) The predicted B-cell epitopes were further assessed through VaxiJen v2.0 (Doytchinova and Flower 2007), AllergenFP v1.0 (Dimitrov et al. 2014) and ToxinPred (Gupta et al. 2013) server for their antigenic, allergenic and toxic profiles, respectively.

#### 2.1.5. Peptide modelling and molecular docking studies

Molecular docking is a computational method used to measure the binding affinity between protein and ligand molecule. The selected epitopes were docked into their respective human leukocytes antigen (HLA) binding alleles to further prove their efficiency to be a part of multi-epitope vaccine. Initially, CTL and HTL epitopes were modeled using PEP-FOLD v3.0 server (Lamiable et al. 2016) with sOPEP sorting scheme in 200 simulations. This server was designed to predict the conformations of small peptides (5-50 amino acids) based on Forward Backtrack/Taboo Sampling algorithm (Lamiable et al. 2016). The crystal structure of the alleles with co-crystallized ligands was then downloaded from RCSB Protein Data Bank (https://www.rcsb.org/). The structures were processed with Discovery Studio followed by the generation of a grid box around their native active sites with AutoDock tools (Morris et al. 2009). For CTL epitopes, allele HLA-B*0702, HLA-A*1101, HLA A*0101 and HLA-B*3901 were considered while DRB1*1101, DQA1*0301, and DRB1*0101 were used for HTL epitopes. Finally, the docking simulation was performed using AutoDock Vina (Trott and Olson 2009). As a positive control, the co-crystallized ligands were considered. The PyMOL Molecular Graphics System v2.2.3 was used to make docked complex and visualized in BIOVIA Discovery Studio 2017 (https://3dsbiovia.com).

#### 2.1.6. Estimation of population coverage

The distribution and expression of HLA alleles vary according to the ethnicities and regions throughout the world (Adhikari, Tayebi, and Rahman 2018), thus, influence the successful development of an epitope-based vaccine (Bui et al. 2006). The population coverage by the designed vaccine was calculated with the IEDB Population Coverage tool (Bui et al. 2006). For this purpose, selected CTL and HTL epitopes and corresponding HLA binding alleles were considered. This tool estimates population coverage of each epitope for different regions of the world based on the distribution of human MHC binding alleles. In this study, however, areas of particular importance regarding our pathogen will be emphasized such as *M. ulcerans* endemic regions and areas of Buruli ulcer occurrence.

### 2.2. Designing and evaluation of multi-epitope vaccine

#### 2.2.1. Mapping of vaccine construct

The multi-epitope vaccine was designed by joining adjuvant, T-cell and B-cell epitopes with appropriate linkers as described previously (Nezafat et al. 2016; Khatoon, Pandey, and Prajapati 2017). An adjuvant is an immunogenic component which enhance the vaccine’s immunogenicity, hence, should be chosen carefully (Coffman, Sher, and Seder 2010). For example, it should have affinity towards immune receptor such toll-like receptor (TLR) receptor (Kaisho and Akira 2002). The TLR2 on epidermal keratinocytes are actively involved in the internalization of *M. ulcerans* and concomitant innate immune response. Therefore, TLR2 is essential for provoking an immune response against *M. ulcerans* (Lee et al. 2009). In addition, PE-PGRS protein interacts with TLR2 on macrophages and antigen presenting cells such as dendritic cells (Brennan 2017). Moreover, TLR agonists (*i.e.*, lipoproteins, lipopolysaccharides, monophosphoryl lipid A, and mannan-protein) are well-known to be used as vaccine adjuvants (Gnjatic, Sawhney, and Bhardwaj 2010; Kumar, Sunagar, and Gosselin 2019). The mycobacterial lipoprotein LprG is a prominent TLR2 agonist (Gehring et al. 2004). Therefore, *M. ulcerans* derived LprG (GenBank: ABL04283) was taken as the adjuvant in this study (Stinear et al. 2007). The EAAAK linker facilitates the effective separation of bifunctional fusion protein domains (Arai et al. 2001), therefore, utilized to integrate the adjuvant and first HTL epitope. For the efficient recognition of epitopes within the vaccine, GPGPG and AAY linkers were used to merge HTL and CTL epitopes, respectively. The LBL epitopes were added together with bi-lysine (KK) linker to preserve their independent immunogenic activities as shown before (Gu et al. 2017).

#### 2.2.2. Primary and secondary structural analysis

The physicochemical properties of the designed vaccine construct, including molecular weight (MW), theoretical isoelectric point (theoretical PI), instability index (II), aliphatic index (AI), and grand average of hydropathicity (GRAVY), *in vitro* and *in vivo* half-life were assessed using ProtParam server (Wilkins et al. 1999). Moreover, antigenic and immunogenic profiles were predicted with VaxiJen v2.0 server and IEDB Immunogenicity tool, respectively. The vaccine protein should be free from allergic reaction. Therefore, AllergenFP v1.0 server was used to predict the allergenic nature of the construct. Furthermore, the solubility of the construct upon overexpression in *E. coli* host was predicted by SOLpro tool (Magnan, Randall, and Baldi 2009). This server use 10-fold cross-validation method and provides more than 74% accurate prediction. We also use Protein-Sol web-tool to further clarify the solubility of the chimeric protein (Hebditch et al. 2017). Finally, the secondary structural features were revealed using the primary with PSIPRED 4.0 server that uses two feed-forward neural networks for precise prediction based on position-specific scoring matrix (Buchan et al. 2013).

#### 2.2.3. Tertiary structure modelling

The 3D structure is the lowest energy state of a protein which provides the maximum stability through proper twisting and bending. RaptorX server was used to predict the tertiary structure of the vaccine construct (Källberg et al. 2012). It predicts 3D protein model based on multiple-template threading (MTT) and provides some confidence scores as a quality indicator. For example, *p*-value for relative global quality, global distance test (GDT), and unnormalized GDT (uGDT) for absolute global quality, and modelling error at each residue (Källberg et al. 2012). Finally, the tertiary structure was visualized with BIOVIA Discovery Studio 2017.

#### 2.2.4. Refinement and validation of tertiary structure

The crude vaccine model should be refined to carry out further evaluation. For this purpose, we used GalaxyRefine web-server (Ko et al. 2012) which uses CASP10 tested refinement method (Nugent, Cozzetto, and Jones 2014). It rehashed structure perturbation followed by overall structural relaxation through dynamics simulation (Pandey, Bhatt, and Prajapati 2018). Further, refined structure needs to be validated based on experimentally validated 3D protein structure. Therefore, the refined vaccine protein was applied in ProSA-web (Wiederstein and Sippl 2007) which provides an overall quality score for a given structure. The quality score outside the usual range of native proteins indicates possible errors in the predicted protein structure. Besides, ERRAT server (Colovos and Yeates 1993) was used to evaluate the statistics of non-bonded interactions. Ramachandran plot was created with the RAMPAGE server (Ramachandran, Ramakrishnan, and Sasisekharan 1963; Lovell et al. 2003) where PROCHECK principle is applied to validate protein structure based on energetically allowed and disallowed dihedral angles psi (ψ) and phi (ϕ) of amino acid residues (Laskowski et al. 1993). The crude vaccine model was also subjected to validation as a control for refined model.

#### 2.2.5. Screening for conformational B lymphocyte epitopes

Since antibody-mediated humoral immunity initiates when B-cells meet their epitopic counterparts, vaccine protein should contain B-cell epitopes. IEDB ElliPro tool (Ponomarenko et al. 2008) was applied for the prediction of conformational B-cell epitopes in the final vaccine protein using default setting (*i.e.*, 0.5 min-score and 6Å max-distance). The ElliPro predicts epitopes through an approximation of protein shape, residual protrusion index (PI) and neighbor residue clustering (Ponomarenko et al. 2008).

#### 2.2.6. Disulfide engineering of the vaccine protein

Two adjacent protein chains can be bridged with covalent bond through disulfide engineering which increases the structural stability of a protein (Khatoon, Pandey, and Prajapati 2017; Pandey, Bhatt, and Prajapati 2018). Therefore, the refined vaccine model was submitted to Disulphide by Design 2.12 server (Craig and Dombkowski 2013). This web-platform identifies residue pairs in a given protein that can be suitable for covalent modifications based on their energy score and χ3 value. For disulfide bridges to be formed, the energy score should be lower than 2.2 kcal/mol and χ3 angle should be within −87 to +97 degree as previously optimized (Craig and Dombkowski 2013). Finally, the potential residue pairs were selected and mutated to cysteine residues to allow bridging between them using ‘Create/View Mutant’ function of the Disulfide by Design 2.12 server.

#### 2.2.7. Docking between vaccine protein and TLR2 receptor

Toll-like receptor 2 (TLR2) plays an important role in triggering immune responses to *M. ulcerans* (Lee et al. 2009). In addition, the adjuvant LprG added in the vaccine protein was a TLR2 agonist. Therefore, TLR2 (PDB ID: 3a7c) was considered as the docking receptor, hence, retrieved from the RCSB PDB database (Berman et al. 2000) and the vaccine protein was used as a ligand. Finally, the vaccine protein and TLR2 immune receptor were submitted to the ClusPro v2.0 server for molecular docking (Kozakov et al. 2017). The ClusPro server calculates the binding affinity (ΔE) according to the following equation:

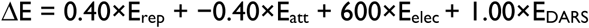

Where, E_rep_ and E_attr_ denote the repulsive and attractive contributions to the van der Waals interaction energy, respectively, E_elec_ defines the electrostatic energy term, and E_DARS_ is a pairwise structure-based potential constructed by the Decoys as the Reference State (DARS) approach (Chuang et al. 2008).

The docking approach of ClusPro server consists of three successive steps such as rigid body docking, clustering of lowest energy structure, and structural refinement (Kozakov et al. 2017). Based on the lowest energy score and binding efficacy, the best docked complex was selected for dynamics simulation.

#### 2.2.8. Gibbs free energy (ΔG) and dissociation constant (K_d_) calculation

The binding affinity of the protein complex is important in therapeutics development (Kastritis and Bonvin 2013) because it determines potentiality and strength of a complex to be formed (Xue et al. 2016). Therefore, the Gibbs free energy (ΔG) of the vaccine-TLR2 complex was predicted with the PRODIGY server (Xue et al. 2016). This web-application works based on pair-wise intermolecular contacts (within 5.5Å distance threshold) of the binding interface in a protein-protein complex and comparing it with an internal dataset comprising 122 complexes with reliable experimental binding affinities (Vangone and Bonvin 2015). Based on the predicted δG, the dissociation constant (K_d_) of the vaccine-receptor complex was also calculated by δG = RT×lnK_d_ relationship where, R and T are the ideal gas constant and temperature in Kelvin scale, respectively. The PRODIGY server calculates δG by the following equation (Vangone and Bonvin 2015).

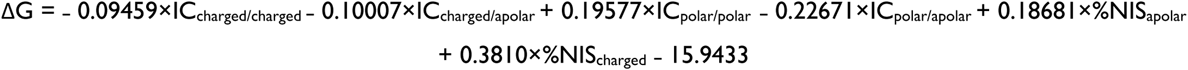

Where, IC_X/Y_ indicates the number of interfacial contacts (IC) found at the interface between chain A and B in terms of the polar, apolar or charged nature of the interacting residues, and %NIS denotes percentage of the apolar and charged non-interacting surfaces (NIS).

In this study, TLR2 and vaccine molecules were defined as chain A and B, respectively and temperature was set to 25°C (298K). The TLR2 protein with co-crystallized SSL3 (*Staphylococcal* superantigen-like protein 3, PDB ID: 5d3i) was used as a positive control.

#### 2.2.9. Molecular dynamics simulation

The study of molecular dynamics is essential for determining the binding stability of a receptor-ligand complex (Pandey, Bhatt, and Prajapati 2018). The molecular dynamics (MD) simulation between TLR2 (as receptor) and vaccine (as ligand) was performed using YASARA Dynamics software v17.8 (Krieger et al. 2004) under AMBER14 force-field (Dickson et al. 2014). Initially, the complex was cleaned and hydrogen was added to all atoms. The whole procedure was accomplished in a TIP3P-solvated (density: 0.997 g/L–1) simulation cell with default macro setting (md_run.mcr) for 3 nanoseconds until the complex achieved its stable state. A cut-off radius of 8.0 Å was considered at predefined physiological state (298 K, pH 7.4, and 0.9% NaCl). The simulation snapshots were captured at every 100 picoseconds. Finally, the simulation-derived trajectories were analyzed using built-in macros (*i.e.*, md_analyze.mcr and md_analyzeres.mcr) to calculate root mean square deviation (RMSD) and root mean square fluctuation (RMSF), respectively. Besides, the number of hydrogen bonds present in the initial and final YASARA scene was evaluated with BIOVIA Discovery Studio 2017.

#### 2.2.10. Immune simulation studies

The *in silico* immune response for the vaccine antigen was measured with a dynamic immune simulator embedded in C-ImmSim server (Rapin et al. 2010). This server takes vaccine construct as an input to predict the epitopes and generate real-life-like immune interactions based on position-specific scoring matrix and machine learning techniques, respectively (Rapin et al. 2010). The minimum recommended interval between dose 1 and dose 2 for most of the commercial vaccines is 4 weeks (Castiglione et al. 2012). Therefore, the dose-dependent immune response by the vaccine construct was determined using three injections (doses) with four weeks interval period translated into 1, 84 and 168 time-steps (1 time-step is equivalent to 8 hours in real-life). Furthermore, probable immune response in the pathogen invaded area was calculated using 12 subsequent injections as repeated exposure. In both cases, each dose comprised of 1000 vaccine particles and the simulation was continued for 1050 time-steps (~350 days). The Simpson index (D) was interpreted as a measure of immune response diversity (Rapin et al. 2010).

#### 2.2.11. Codon adaptation and in silico cloning

Codon usage in organism differs from species to species, hence, unadapted codon may result in the minor expression rate in the host (Pandey, Bhatt, and Prajapati 2018). Therefore, it should be optimized according to the host translational machinery to improve gene expression (Khatoon, Pandey, and Prajapati 2017). In this study, we used Java Codon Adaptation Tool (JCAT) server (Grote et al. 2005) to adapt our vaccine codon for a widely used prokaryotic model organism, *E. coli* strain K12. To avoid the prokaryote ribosome binding site, rho-independent transcription termination, and restriction enzymes cleavage sites, three additional options at the bottom were selected. The codon adaptation index (CAI) value (Sharp and Li 1987) and guanine-cytosine (GC) content of the adapted sequence were calculated. The CAI score is the measure of codon usage biasness and an optimal CAI score should be 1.0 but score greater than 0.8 can be considered as good (Morla, Makhija, and Kumar 2016). An optimal score of GC content range in between 30–70% (Khatoon, Pandey, and Prajapati 2017). Then, XhoI and NcoI restriction sites were introduced to the N and C-terminal of the optimized nucleotide sequence, respectively. Finally, the adapted deoxyribonucleic acid (DNA) sequence of the designed vaccine was cloned into the multiple cloning site (MCS) of *E. coli* plasmid pET30a(+) with SnapGene 4.3 tool (https://snapgene.com/) to ensure the expression of desired vaccine protein.

#### 2.2.12. Data availability

Data not embedded within the manuscript are provided as a supplementary file. The protein sequences used in this study can be retrieved from IMG/M, RCSB PDB and NCBI database using their corresponding accession codes.

## 3. Results

### 3.1. Pre-vaccine design analysis

#### 3.1.1. Selection of PE-PGRS family protein

The proteome of the *M. ulcerans* Ag99, consisting of 4047 proteins, excluding human homologs, was retrieved from IMG/M system and subjected to antigenicity prediction. Based on the antigenic scores, the top-ten proteins were selected for which the scores were ranged from 1.825 to 2.394. All the 10 proteins belonged to the conserved proline-glutamate polymorphic GC-rich sequence (PE-PGRS) family protein of the *M. ulcerans*. Moreover, no transmembrane helices was found for any of those proteins as shown in Table 1. Therefore, we selected the PE-PGRS protein (GenBank: WP_011740336) with the highest antigenic score (2.3941) that contains 642 amino acid residues.

**Table 1:** The top 10 antigenic proteins resulted in the initial proteome screening.

| SN | IMG/M Identifier | Protein name | Antigenicity score | Number of TM helices | Length (AA) |
| --- | --- | --- | --- | --- | --- |
| 1 | 642569202 | PE-PGRS family protein | 2.394 | 0 | 642 |
| 2 | 642570886 | PE-PGRS family protein | 2.308 | 0 | 929 |
| 3 | 642569998 | PE-PGRS family protein | 2.231 | 0 | 667 |
| 4 | 642570321 | PE-PGRS family protein | 2.133 | 0 | 558 |
| 5 | 642570805 | PE-PGRS family protein | 2.060 | 0 | 728 |
| 6 | 642568686 | PE-PGRS family protein | 1.995 | 0 | 461 |
| 7 | 642570293 | PE-PGRS family protein | 1.940 | 0 | 458 |
| 8 | 642568458 | PE-PGRS family protein | 1.930 | 0 | 403 |
| 9 | 642568210 | PE-PGRS family protein | 1.925 | 0 | 789 |
| 10 | 642569115 | PE-PGRS family protein | 1.825 | 0 | 636 |

#### 3.1.2. Evaluation and selection of T and B-cell epitopes

Total 58 unique CTL epitopes (9-mer) were predicted from the *M. ulcerans* derived PE-PGRS protein in which 24 epitopes have MHC-I binding alleles. The number of alleles for CTL epitopes was ranged from 1 to 17 (Table S1). After evaluation, 14 epitopes were found as antigenic, immunogenic, non-toxic and had at least 5 MHC-I binding alleles as shown in Table 2. Further, allergenicity assessment showed 5 CTL epitopes were non-allergic, hence, selected for vaccine construction (Table 2). Similarly, a total of 119 unique HTL epitopes (15-mer) were predicted along with their respective MHC-II binding molecules. Firstly, the HTL epitopes were evaluated based on the number of alleles which resulted in 53 epitopes having a minimum of 5 binding alleles (Table S2). Based on the cytokine inducing ability, 13 HTL epitopes were found as IFN-γ and IL-4 positive in which two epitopes were also IL-10 inducers as provided in Table 3. Finally, 7 epitopes (including 3 overlapping epitopes) were selected based on their antigenic score at 0.5 threshold (Table 3). B-cell epitopes are antigenic regions of a protein that can trigger antibody formation. We found a total of 199 linear B-cell epitopes (12-mer) with probability score 0.502 to 0.843 (Table S3). After evaluation, however, only 6 B-cell epitopes with probability score higher than 0.8 were found to be highly antigenic, non-allergenic and non-toxic as presented in Table 4. Finally, 3 epitopes were selected as unique, therefore, considered for vaccine construction (Table 4).

**Table 2:**
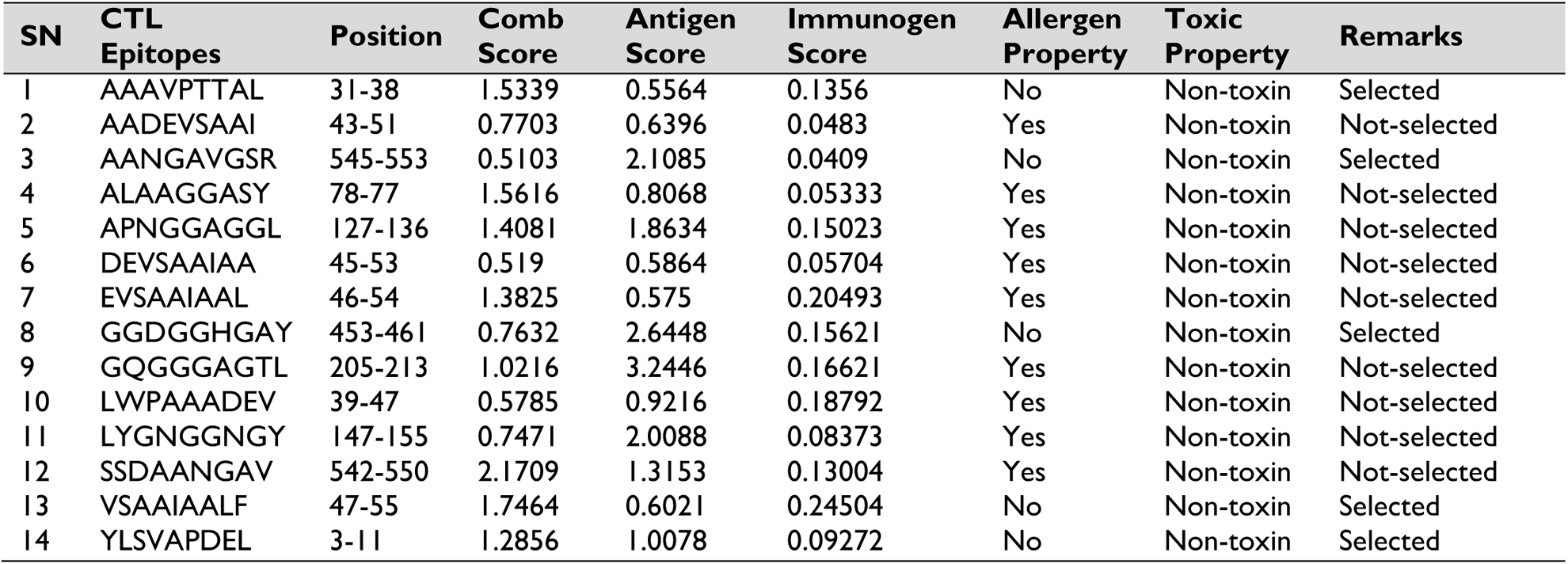
The evaluation and selection of cytotoxic T-lymphocyte (CTL) epitopes.

**Table 3:**
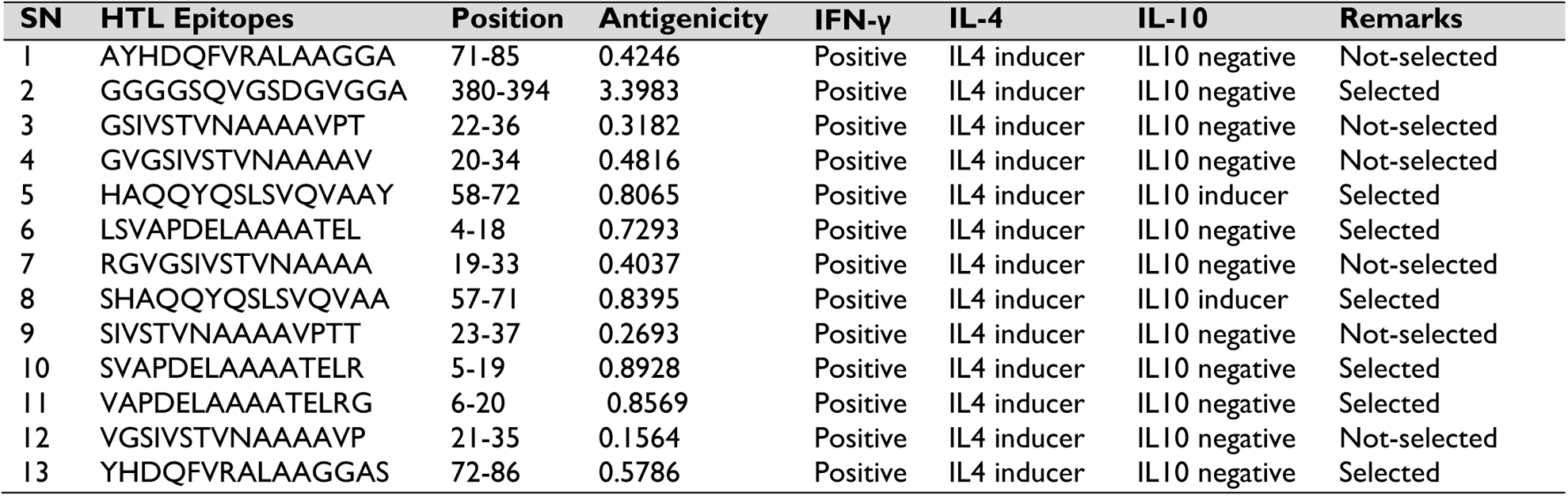
The evaluation and selection of helper T-lymphocyte (HTL) epitopes.

**Table 4:**
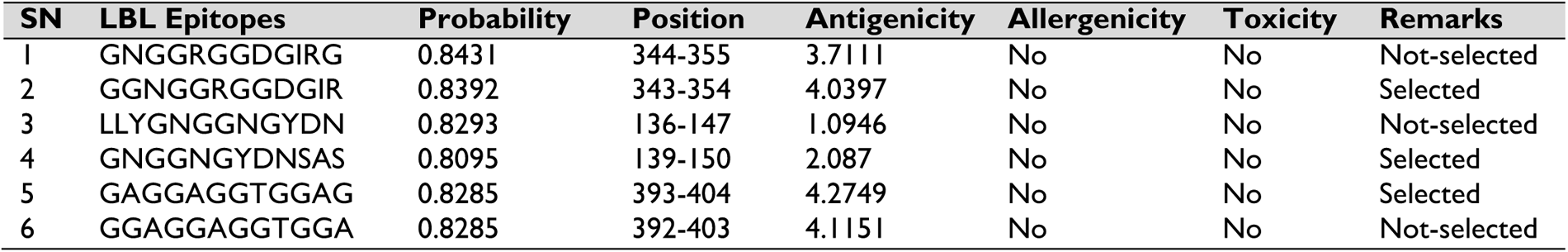
The evaluation and selection of linear B-lymphocyte (LBL) epitopes.

#### 3.1.3. Molecular docking between epitopes and HLA alleles

The epitopes selected for vaccine construction are meant to have a good affinity towards their HLA binding alleles. In this study, the binding affinity of the CTL epitopes and control was ranged within −6.3 to −9.2 and −6.7 to −9.6, respectively as given in Table 5. For HTL epitopes, the binding affinity was determined in between −6.2 to −7.8 for epitopes while control provided −5.5 to −7.8 kcal/mole (Table 5). Therefore, 2 CTL and 4 HTL epitopes showed better binding affinity towards their respective HLA alleles as compare to control ligands. The rest of the epitopes showed binding affinity relatively close to the control ligands. The amino acid residues involved in the hydrogen bond interactions were also analyzed and included in Table 5. Here, Fig. 2 shows the best CTL (YLSVAPDEL) and HTL (YHDQFVRALAAGGAS) epitopes in terms of docking score and rest are shown in Fig. S1.

**Table 5:**
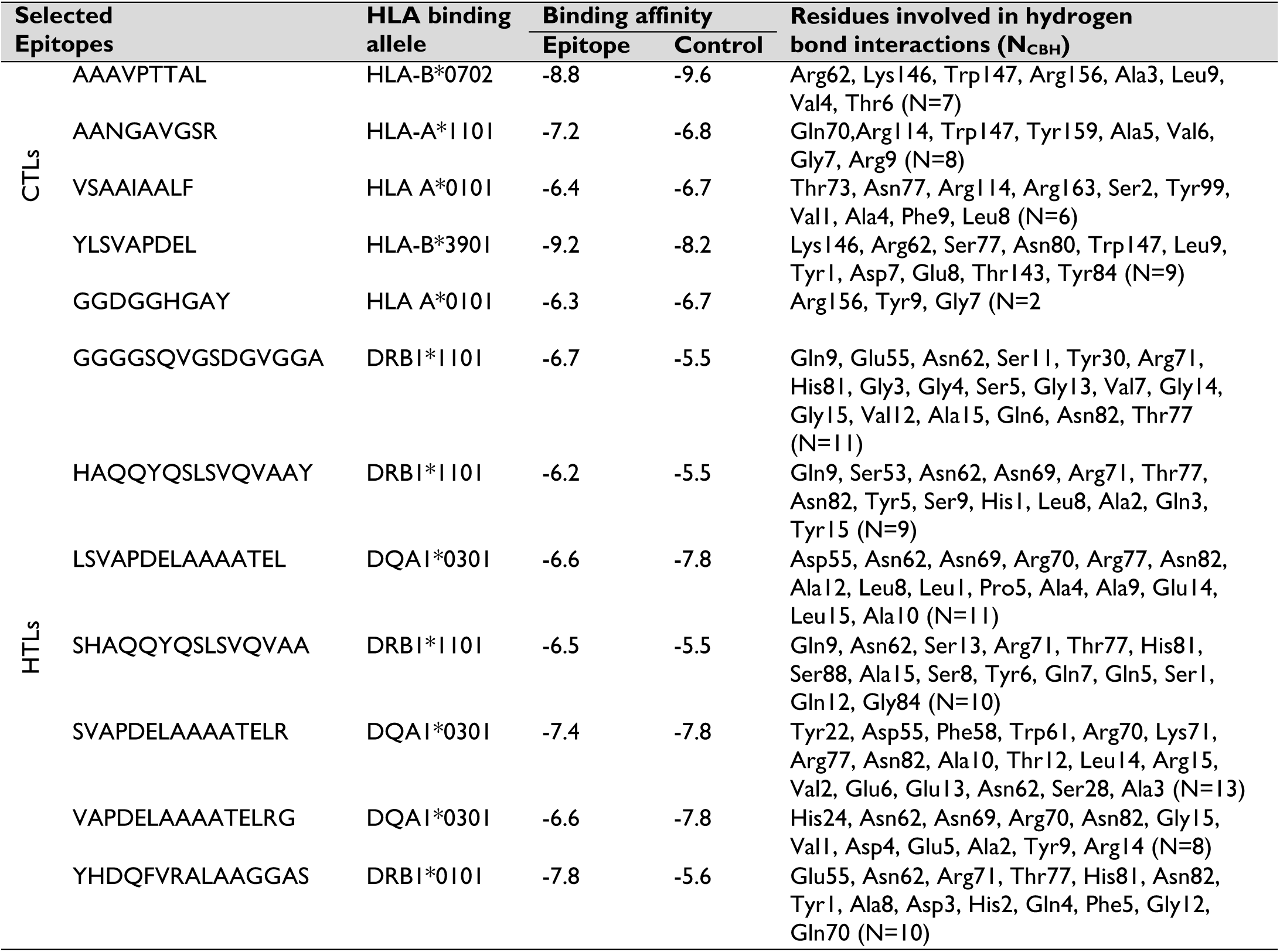
The efficacy of the selected CTL & HTL epitopes to be used in vaccine construction.

**Fig. 2:**
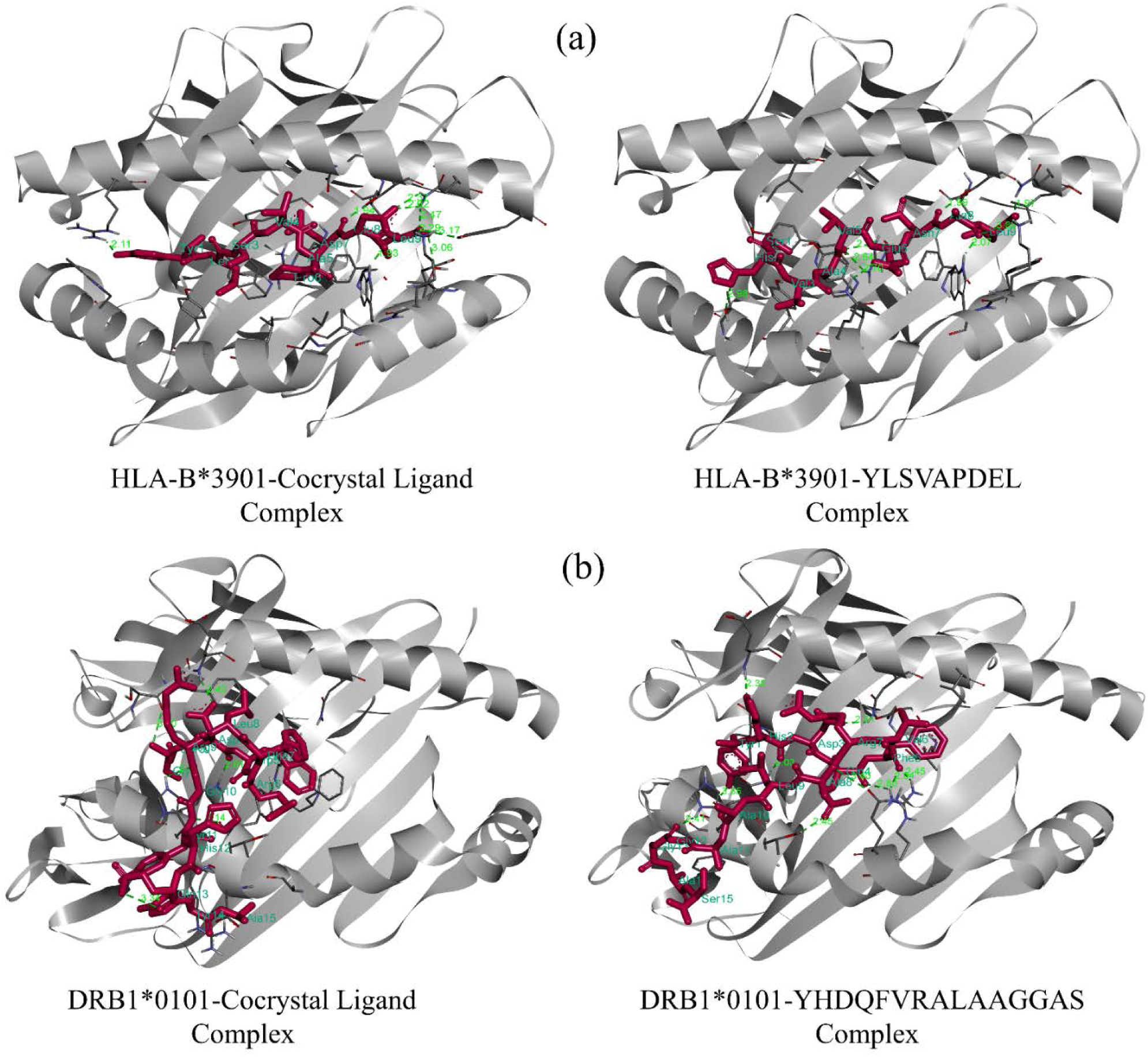
Molecular docking between T-cell epitopes and their respective HLA binding alleles. (a) docking of HLA-B*3901 allele with the co-crystal ligand (left) and the best CTL epitope (YLSVAPDEL) (right) where residue Lys146, Arg62, Ser77, Asn80, Trp147, Leu9, Tyr1, Asp7, Glu8, Thr143, and Tyr84 were involved forming total nine conventional hydrogen bonds, (b) docking of DRB1*0101 with the co-crystal ligand (left) and the best HTL epitope (YHDQFVRALAAGGAS) (right) where residue Glu55, Asn62, Arg71, Thr77, His81, Asn82, Tyr1, Ala8, Asp3, His2, Gln4, Phe5, Gly12, and Gln70 took part to provide ten conventional hydrogen bonds.

The best CTL epitope ‘YLSVAPDEL’ formed 9 hydrogen bonds with 11 active site residues (Table 5) while respective control provided 8 hydrogen bonds which involved amino acid residues Ser77, Asn80, Arg97, Lys146, Trp147, Gln155, His2, Asn7, Leu9, Ala4, Ala8, Glu6, and Thr163 of active site. On the other hand, best HTL epitope ‘YHDQFVRALAAGGAS’ was involved with 14 residues (Table 5) through 10 hydrogen bond interactions while control ligand interacted with residues Glu55, His81, Asn82, Ser3, Tyr11, Gln13, Asp4, Arg9, Ser53, Thr77 to form 6 hydrogen bonds. In conclusion, all tested epitopes showed good binding affinity towards the respective HLA alleles. Therefore, the selected epitopes were considered as the most potential epitopes to be used in multi-epitope based vaccine design.

#### 3.1.4. World population coverage

The population coverage was estimated with finally selected 5 CTL and 7 HTL epitopes and corresponding HLA alleles. Our selected CTL and HTL epitopes showed 99.55% and 56.36% of global population coverage, respectively (Table S4). Since vaccine will contain both types of epitopes, we emphasized on their combined coverage which was 99.8% of world population as shown in Fig. 3. The highest coverage (100%) was found within the population from two countries such as Bulgaria (other ethnic group) and Peru (Fig. 3). In Uganda, where *M. ulcerans* was first identified, the population coverage was 96.9% (Fig. 3b). Also, the population coverage was higher in regions where Buruli ulcer occurrence have been reported such as West Africa (98.59%), Central Africa (97.13%), South America (99.63%), and Western Pacific regions *i.e.*, Australia (96.67%), Japan (99.74%), New Zealand (47.99%), and Singapore (96.55%) (Fig. 3 and Table S4). The population coverage for South Asia, India and Europe were 99.75%, 99.63% and 99.84%, respectively. However, the least population coverage (0%) was predicted for the European country Slovakia (Table S4), which is plausible due to the lower allele frequency based on the data available in Allele Frequencies (http://allelefrequencies.net) database.

**Fig. 3:**
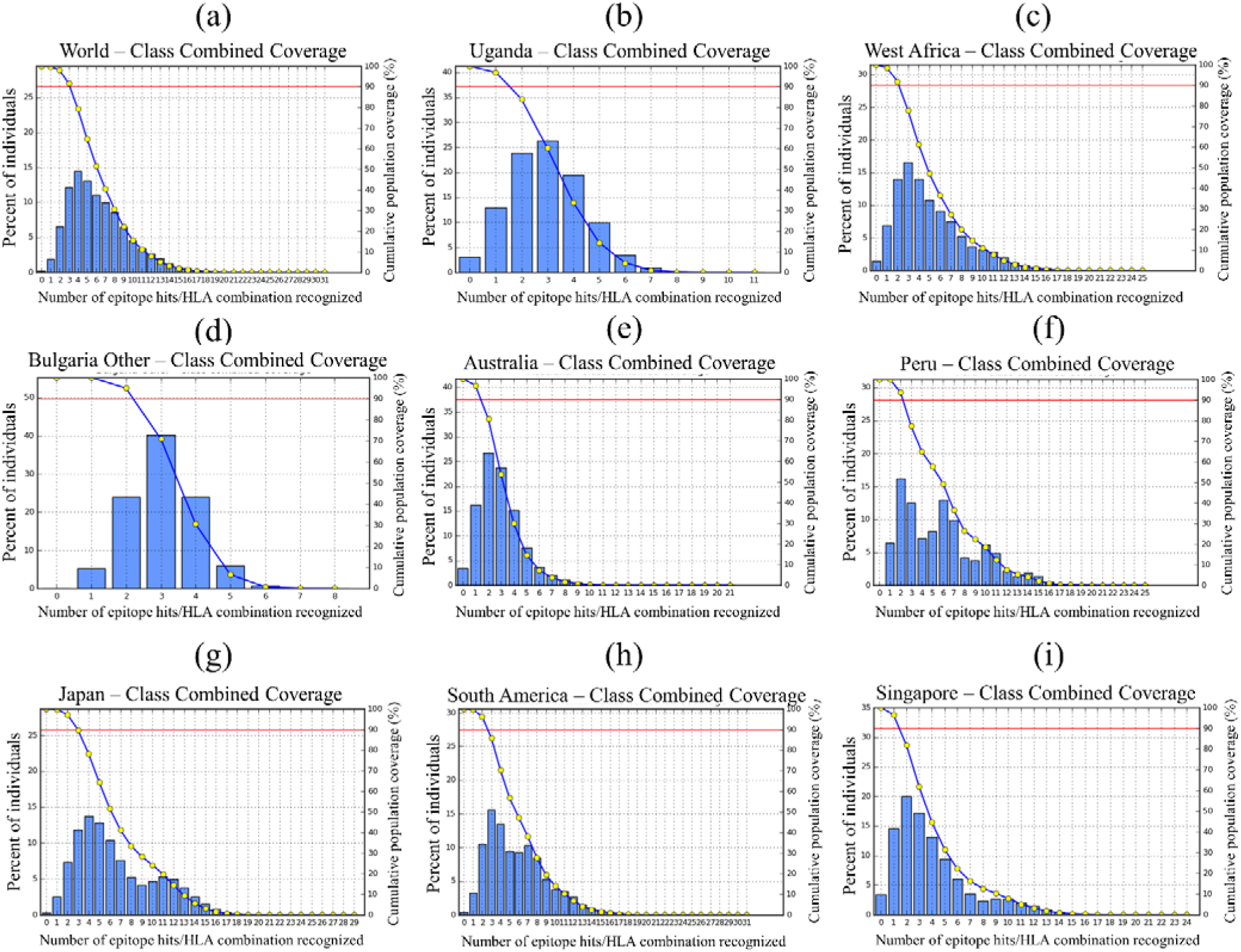
Worldwide population coverage by T-cell epitopes based on their respective HLA binding alleles. Regions of particular interest were considered here: (a) World, 99.80%; (b) Uganda, 96.90%; (c) West Africa, 98.59% Bulgaria other, 100%; (e) Australia, 96.67%; (f) Peru, 100%; (g) Japan, 99.74%; (h) South America, 99.63%; (i) Singapore, 96.55% coverage.

### 3.2. Vaccine design and post-vaccine studies

#### 3.2.1. Construction of multi-epitope vaccine

A total of 15 epitopes were arranged in an aforementioned way to construct the chimeric vaccine protein. The selected 7 HTL, 5 CTL and 3 LBL epitopes were merged with GPGPG, AAY, and KK linkers, respectively. The vaccine construct contains epitopes in order of HTL, CTL and LBL in which epitopes in each group were mapped according to their higher to lower antigenic score. Furthermore, TLR2 agonist LprG protein (236 residues) was added as an adjuvant to the first HTL epitope using EAAAK linkers. The arrangement of different epitopes along with their joining linkers is shown in Fig 4. The final vaccine construct comprises 520 amino acid residues.

**Fig. 4:**
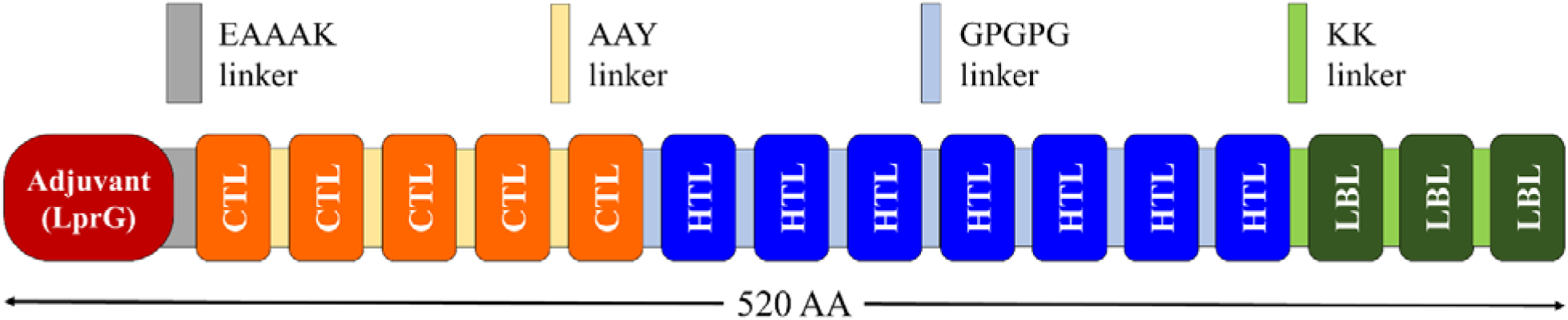
The organization of T and B-cell epitopes in multi-epitope vaccine construct. Finally constructed protein chimera was 520 amino acid residues long in which first 241 residues have been represented as adjuvant (*M. ulcerans* lprG protein) attached to EAAAK linker (AA_1_-AA_241_) followed by seven HTL epitopes with GPGPG linkers (AA_242_–AA_376_), five CTL epitopes with AAY linkers (AA_377_–AA_436_), and three LBL epitopes with KK linkers (AA_437_-AA_520_).

#### 3.2.2. Physicochemical and immunogenic profiles

The evaluated physicochemical and immunogenic properties of vaccine construct are shown in Fig. 5. The construct was found to be slightly basic (theoretical PI > 7), stable (Instability Index < 40), and hydrophilic (negative GRAVY value) in nature. The aliphatic index (AI) contributes to the protein’s thermal stability and protein with higher AI score are more thermostable (Panda and Chandra 2012). Therefore, the construct could be thermostable (AI value of 74.12) and it may also contain high amount of hydrophobic amino acids. Moreover, both SolPro and Protein-Sol servers predicted the construct as highly soluble (≈1.0) during overexpression in *E. coli* system (Fig. 5 and 7b). Besides, the half-life was estimated to be about 30 hours in mammalian reticulocytes (*in vitro*), more than 20 hours in yeast (*in vivo*) and over 10 hours in *E. coli* (*in vivo*). Furthermore, the immunogenic assessment also revealed that the vaccine construct is highly antigenic (0.5 threshold), immunogenic (positive value) and non-allergenic (Fig. 5). These results indicated that the designed chimeric construct is suitable to be a potential vaccine candidate.

**Fig. 5:**
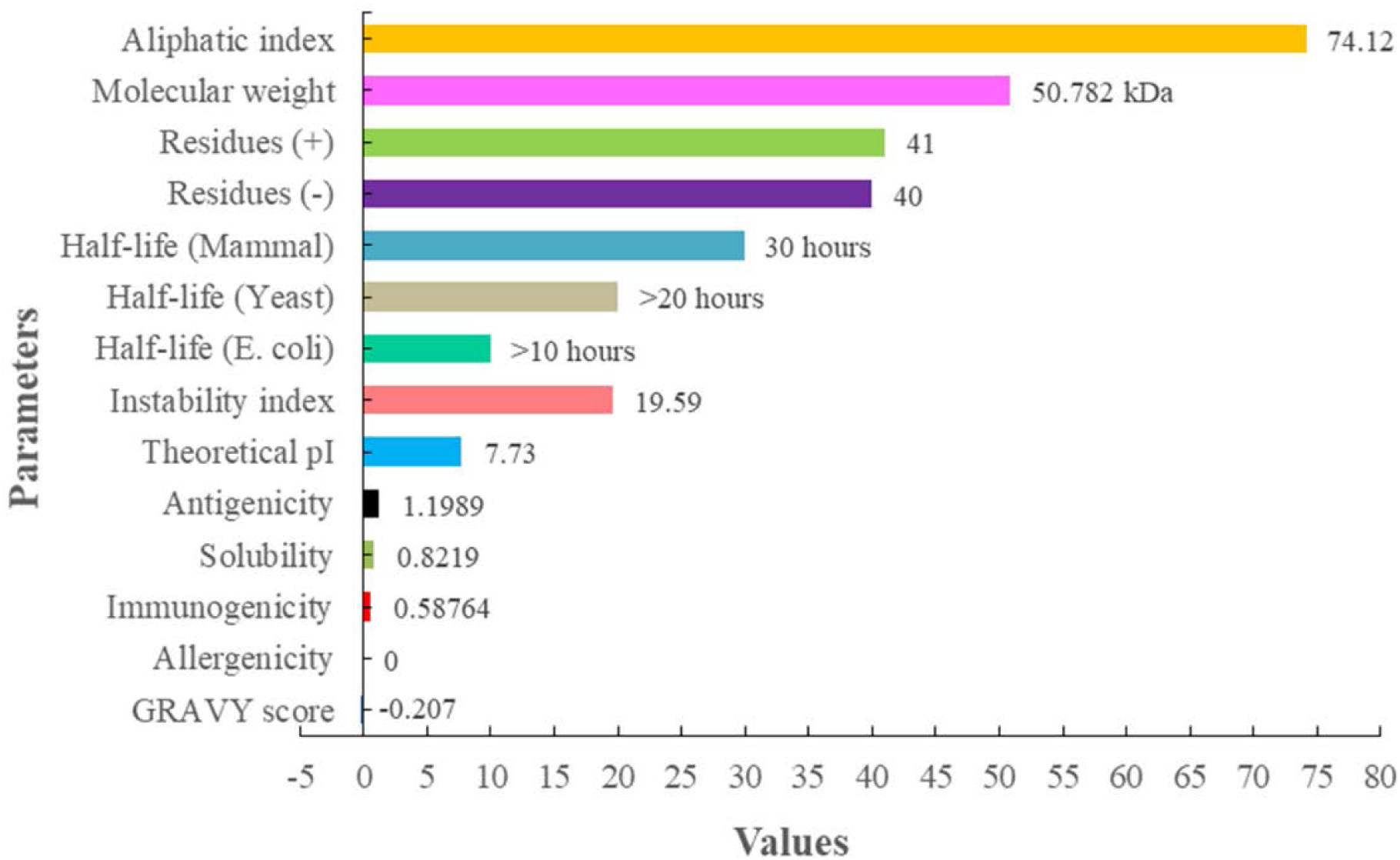
Physicochemical and immunogenic profile of the designed vaccine protein. The vaccine construct is 50.78 kDa in size, slightly basic (theoretical PI above 7.0), hydrophilic (negative GRAVY value), stable (II below 40), thermostable (higher AI value), antigenic (0.5 threshold), immunogenic (positive score), non-allergenic (null), and highly soluble (≈1.0) upon overexpression within *E. coli* system.

**Fig. 6:**
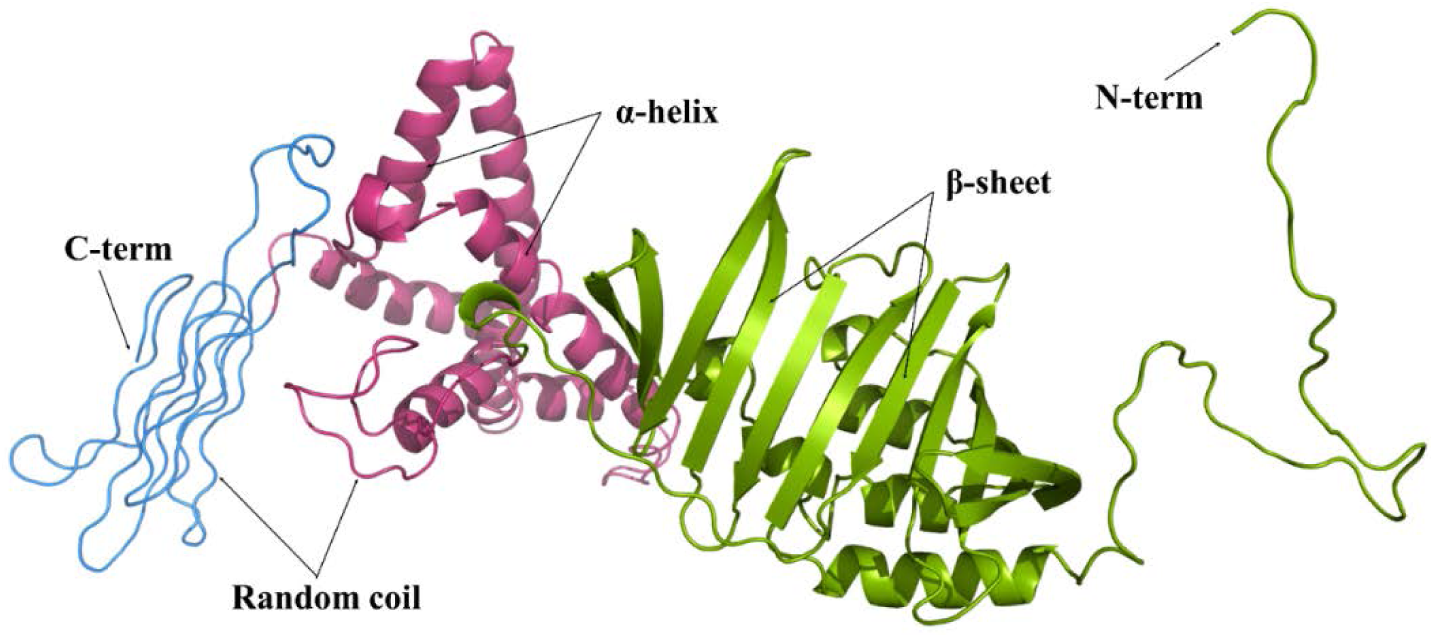
The three-dimensional structure of the vaccine construct. The figure shows predicted domain-1 (AA_1-241_), domain-2 (AA_242-420_), and domain-3 (AA_421-520_) of the vaccine protein in green, pink and blue color, respectively.

**Fig. 7:**
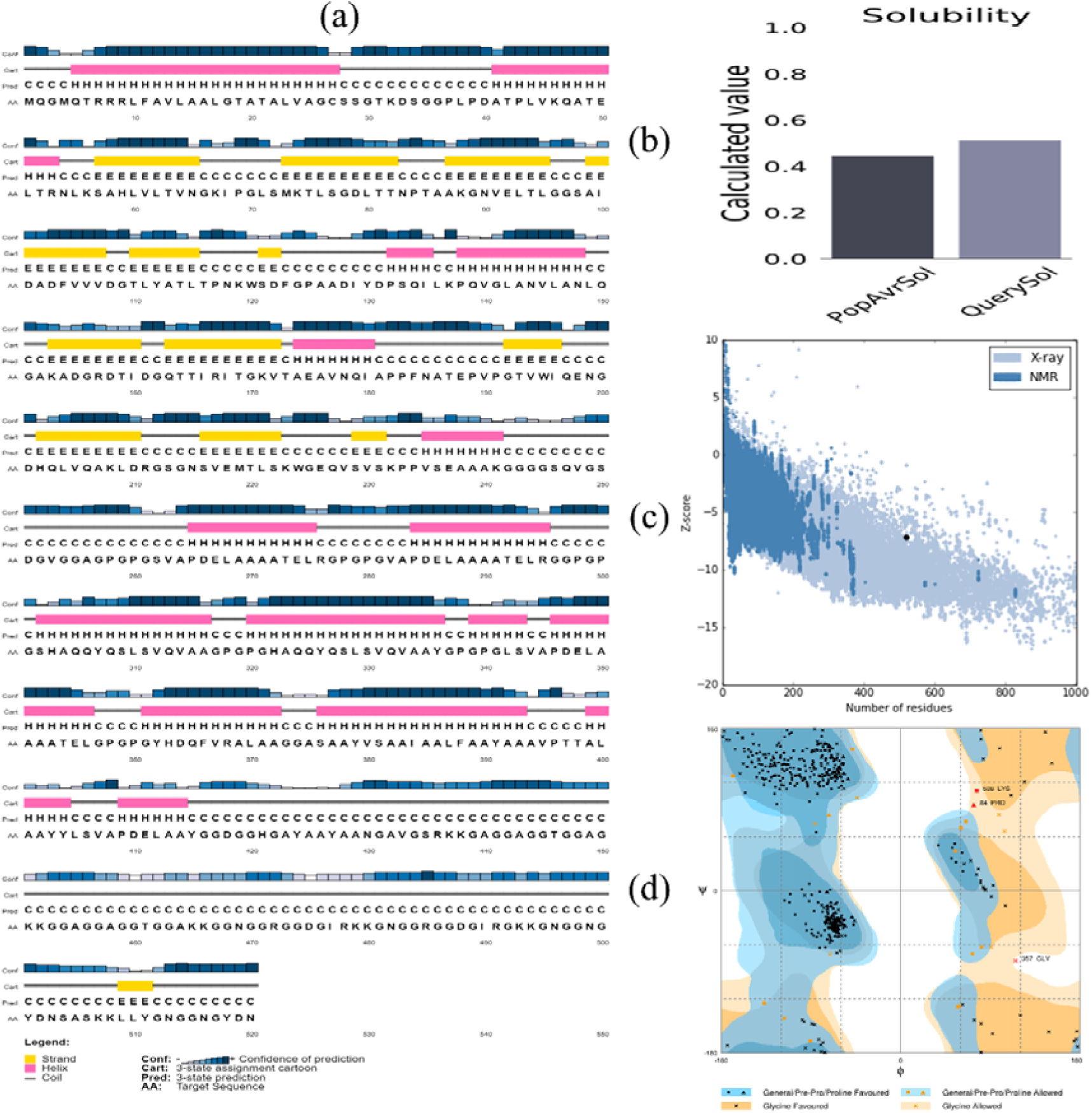
The secondary structural features and assessment of the tertiary structure. (a) the vaccine construct contains α-helix (34.23%, 178), β-strand (17.31%, 90) and random coil (48.46%, 252); (b) the solubility of the vaccine protein is higher than average protein as suggested by Protein-Sol server, (c) the Z-score (−7.14) of the vaccine model indicates quality closer to X-ray crystal protein structure and refined positively; and (d) the Ramachandran plot of refined structure shows 95.8%, 3.7% and 0.6% residues in favored, allowed and disallowed region, respectively.

#### 3.2.3. Secondary and tertiary structures

The secondary structural analysis demonstrated the presence of 34.23% α-helix (178), 17.31% β-sheet (90) and 48.46% random coil (252) structure in the formulated vaccine construct (Fig. 7a). Furthermore, homology modelling of vaccine protein was performed using the RaptorX server. A total of 520 amino acid residues was modelled as three-domains (Fig. 6). The RaptorX server uses multi-template based approach to build a protein model. For example, the template PDB ID: 3mh9_A and 3mh8_A were chosen to model domain 1 (1-241 residues, *p*-value: 1.43e-09). Similarly, for domain 2 (242-420 residues, *p*-value: 1.67e-04) PDB ID: 2a65_A, 4us3_A, and 5i6x_A; and domain 3 (421-520 residues, *p*-value: 1.43e-07) PDB ID: 6f45_D were considered (Table S7). The *p*-value is a quality indicator in homology modelling where *p*-value lower than 10^-3 defines higher model quality. Since we found very small and significant *p*-value, the chimeric protein was considered to be of good quality.

#### 3.2.4. Tertiary structure refinement and validation

Structure refinement with GalaxyRefine leads to an increased number of residues in the favored region. Of all the refined models, the model 2 proved to be the best based on various parameters including GDT-HA (0.9558), RMSD (0.413), MolProbity (1.974), Clash score (14.0), Poor rotamers (0.6) and Rama favored (95.4) as shown in Table S8. This model was taken as the final vaccine model and subjected to structural validation along with the crude model. The quality score (*e.g.*, Z-score) of the refined model was found to be −7.14 as compared to −6.47 of crude model which indicated that the quality was improved as presented in Table 6 and Fig. 7c. In Ramachandran plot, 95.8% (496) residues were found in the favored region with 3.7% (19) and 0.6% (3) residues in allowed region and outlier region, respectively (Fig. 7d). On the other hand, crude model showed 94.2% (488) residues in the favored region (Table 6). The overall quality factor (ERRAT score) of the crude model was 76.53 which was increased to 86.84 in the refined vaccine model. Furthermore, the refined model showed 3 errors in PROCHECK operation while crude structure had 4 errors. These results suggest that the quality of the refined model is better than the crude model.

**Table 6:**
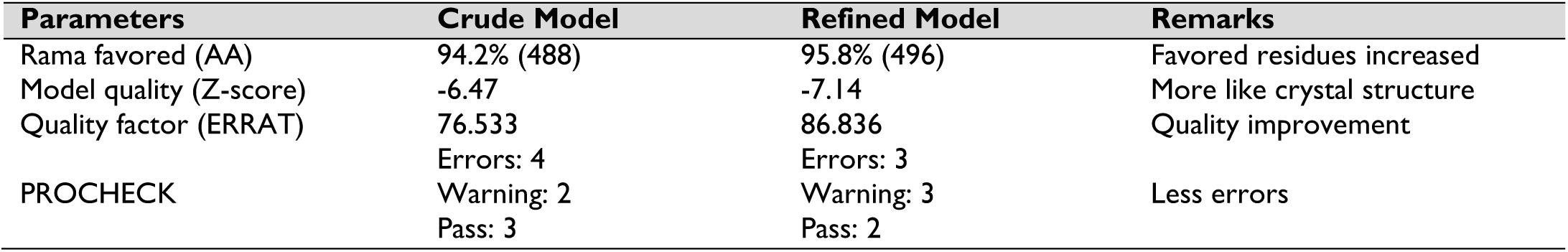
The evaluation of the crude and refined vaccine model for validation.

| Parameters | Crude Model | Refined Model | Remarks |
| --- | --- | --- | --- |
| Rama favored (AA) | 94.2% (488) | 95.8% (496) | Favored residues increased |
| Model quality (Z-score) | -6.47 | -7.14 | More like crystal structure |
| Quality factor (ERRAT) | 76.533 | 86.836 | Quality improvement |
| PROCHECK | Errors: 4 | Errors: 3 | Less errors |
|  | Warning: 2 | Warning: 3 |  |
|  | Pass: 3 | Pass: 2 |  |

#### 3.2.5. Discontinuous B-cell epitopes

A total of 12 conformational B-cell epitopes were identified in the vaccine protein as presented in Fig. 8. The size of the epitopes was fall in between 3 to 98 residues while the numbers of total amino acids were 263 residues as provided in Table S9. Further, the scores of the conformational B-cell epitopes ranged from 0.51 to 0.99.

**Fig. 8:**
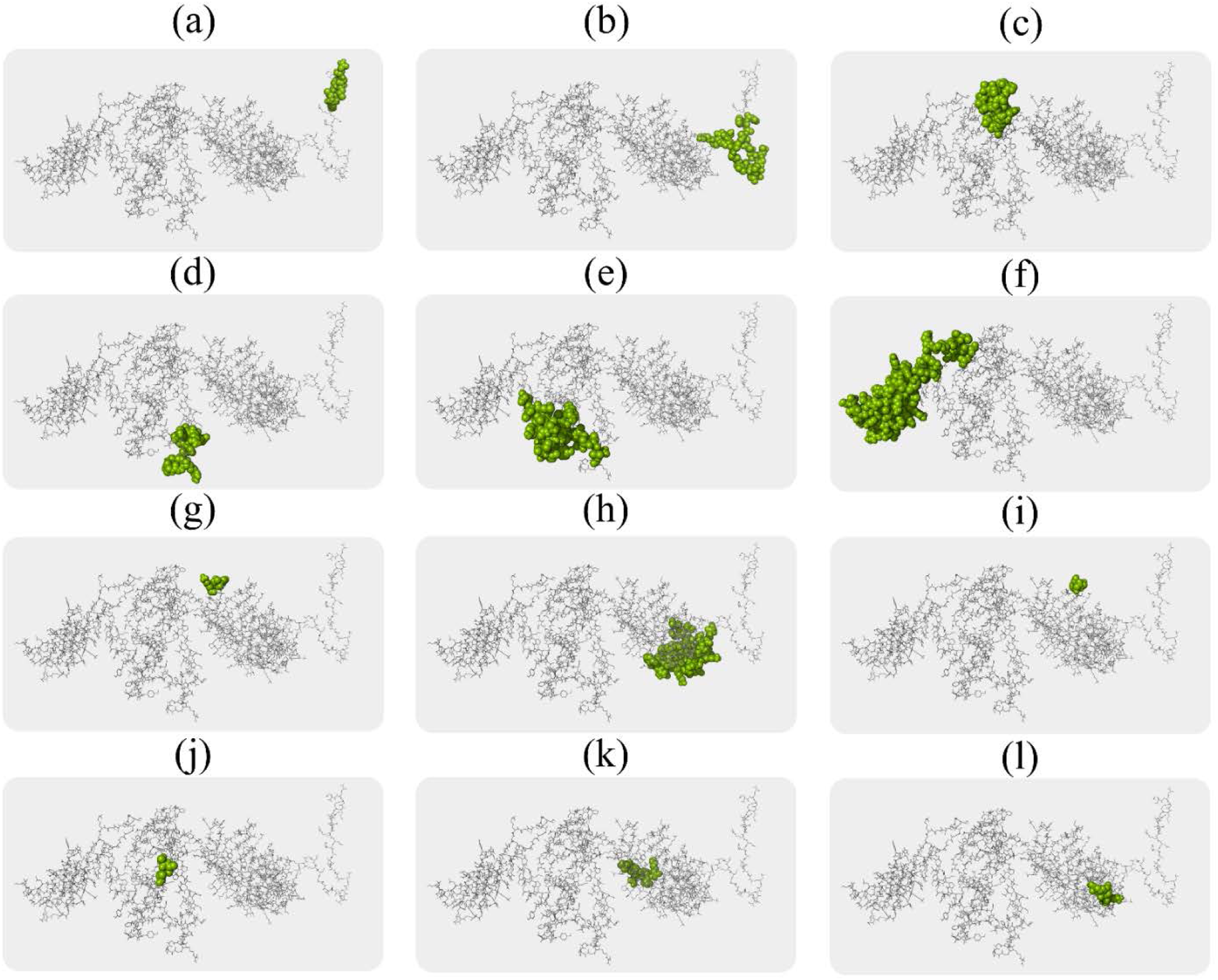
The conformational B-lymphocyte epitopes present in the vaccine. The light green spheres showing epitopes containing (a) 7 residues (AA_2-8_) with 0.99; (b) 23 residues (AA_10-32_) with 0.87; (c) 18 residues (AA_350-365_ and AA_367-368_) with 0.81; (d) 13 residues (AA_295-307_) with 0.79; (e) 38 residues (AA_283_, AA_286-294_, AA_379_, AA_382-407_, and AA_410_) with 0.78; (f) 98 residues (AA_339_, AA_343_, AA_347_, AA_417-419_, AA_423-434_, AA_438_, AA_441-477_, AA_479-520_) with 0.69; (g) 4 residues (AA_116-119_) with 0.67; (h) 43 residues (AA_33-47_, AA_49-51_, AA_53-54_, AA_138_, AA_150-157_, AA_159_, AA_161-164_, AA_166_, AA_169_, and AA_197-203_) with 0.66; (i) 4 residues (AA_96-99_) with 0.64; (j) 3 residues (AA_371-372_, and AA_375_) with 0.53; (k) 7 residues (AA_223_, AA_225-230_) with 0.52; and (l) 5 residues (AA_171_, AA_186-189_) with 0.51 scores.

#### 3.2.6. Vaccine stability improvement

A total of 58 pairs of residues were identified that are suitable for disulfide engineering (Table S10). After evaluating energy score and χ3 angle, only four pair of residues were finalized because their value satisfies the defined criteria *i.e.*, energy score should be less than 2.2 kcal/mol and χ3 value should range from −87 to +97 degree. Therefore, a total of eight mutations were generated on the residue pairs, Asn469-Gly487, Gly457-Asp475, Ala426-Ala432, and Ala174-Ala186, for which the χ3 angle were −77.33, 80.16, −82.98, and 94.45 degree and the energy scores were 0.52, 1.49, 1.99, and 2.04 kcal/mol, respectively as described in Fig. 9.

**Fig. 9:**
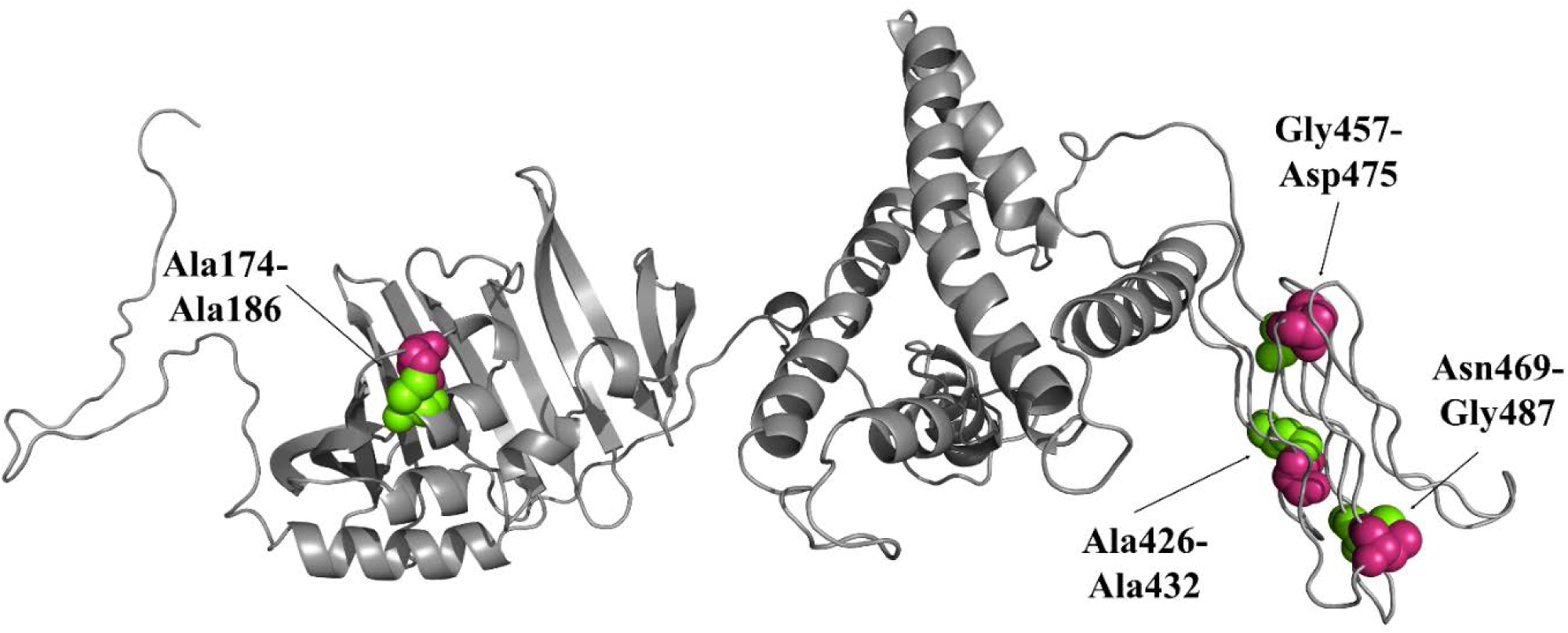
Disulphide engineering of the vaccine protein. Residue pairs showed in pink (Asn569, Gly457, Ala426, and Ala174) and light green spheres (Gly487, Asp475, Ala432, and Ala186) were mutated to Cysteine residues to form disulphide bridge between them.

#### 3.2.7. Interaction between vaccine construct and TLR2 receptor

A total of 30 poses of the vaccine-TLR2 complex was generated in the ClusPro docking and calculated energy score for each pose are tabulated in Table S11. The vaccine protein was found to be docked appropriately into the binding site of the TLR2 receptor in first pose (cluster_0). Moreover, the energy score for this model was found to be the lowest (−1372.6) among all predicted docked-poses, hence, considered as the best docked complex. The pose-view of the selected vaccine-TLR2 complex and their binding interactions was presented in Fig. 10. The binding interactions involved 9 hydrogen bonds, 4 electrostatic contacts, and 11 hydrophobic interactions. Among all hydrogen bonds, seven were classic hydrogen bonds which were provided by Gly3, Gln5, Arg7, Arg8, Arg9, Met1, Gln2, Thr6, Pro320, and Ile319 residues.

**Fig. 10:**
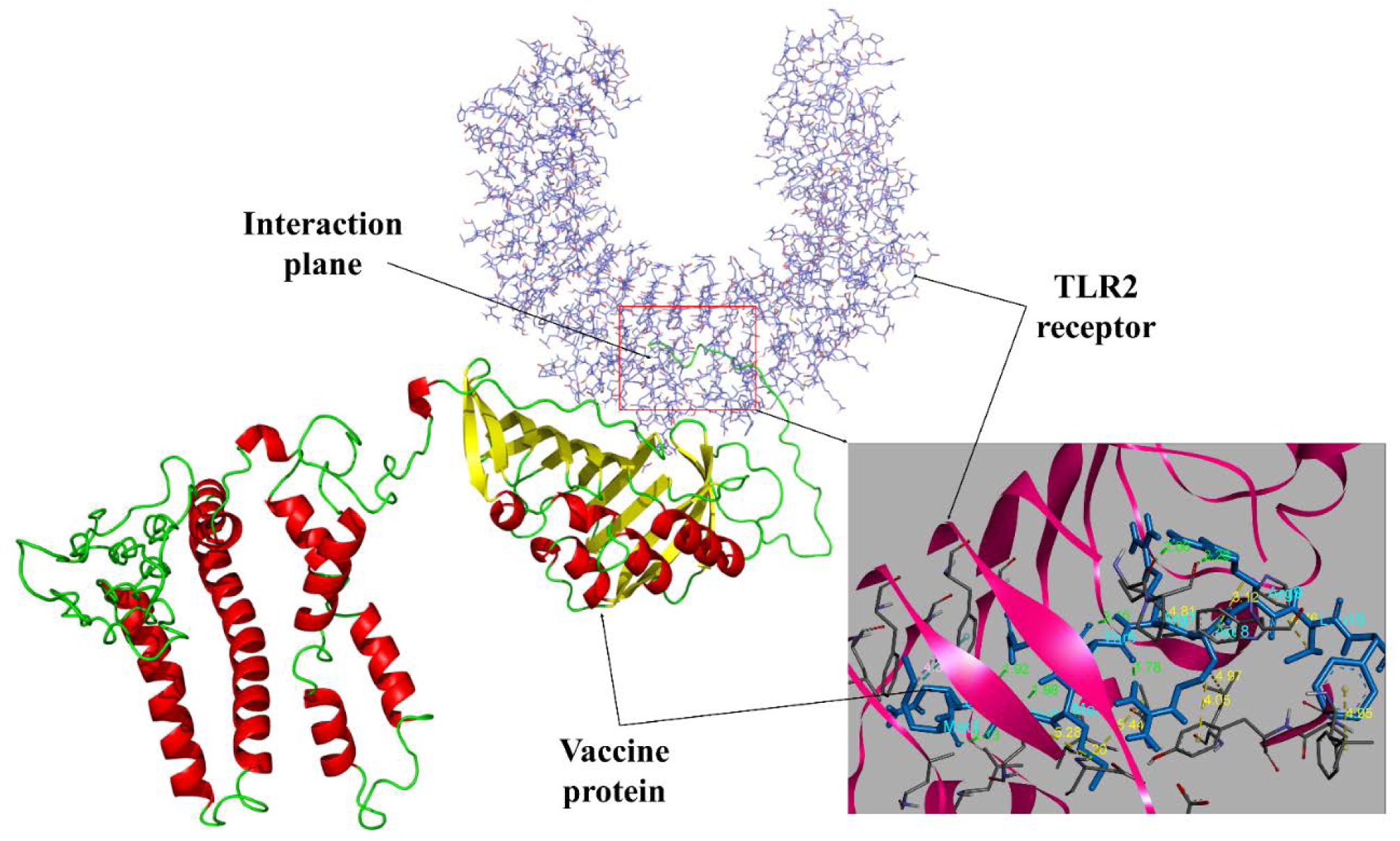
Molecular docking between the vaccine protein and TLR2 immune receptor. The vaccine molecule acts as the ligand while TLR2 as the receptor. The binding interactions are represented in different colors, *i.e.*, hydrogen bonds as green and hydrophobic bonds as yellow. The residues involved in the hydrogen bond interactions are Gly3, Gln5, Arg7, Arg8, Arg9, Met1, Gln2, Thr6, Pro320, and Ile319 as shown in cyan color.

#### 3.2.8. Binding free energy (ΔG) and dissociation constant (K_d_)

The binding interactions of the protein-protein complex depend on the binding free energy. The predicted ΔG of the vaccine-TLR2 complex was −15.3 kcal/mol while SSL3-TLR2 complex provided −9.0 kcal/mol. Therefore, the binding affinity of the vaccine-TLR2 complex was much higher than the control and well comparable to the PRODIGY’s internal dataset (https://nestor.science.uu.nl/prodigy/dataset). This prediction was parallel to the calculated dissociation constant (K_d_) 5.9e-12 M and 2.7e-07 M for vaccine-TLR2 complex and control, respectively. The K_d_ is the separation rate of a protein complex into smaller parts at equilibria. The lower the K_d_, the more tightly bound the ligand is, hence, our vaccine protein showed higher binding affinity towards the TLR2 immune receptor.

#### 3.2.9. Structural integrity of the vaccine-TLR2 complex

The binding stability of vaccine-TLR2 complex was investigated through MD simulation and the results are shown in Fig. 11 and Table S12. The simulations was carried out for 3 nanoseconds since the complex become stabilized around 2.4 nanoseconds with mild fluctuation afterwards (Fig. 9a). The average simulation energy was −15269666.89 kJ/mol, while the Coulombic charge and van der Waals interactions were −20235278.67 and 2861791.649 kJ/mol, respectively (Table S12). The average RMSD of the complex backbone was 7.673 Å (Fig. 9a) while the RMSD of side-chain residues ranged in between 0.522 Å to 10.962 Å (Fig. 11b). We also calculated the RMSF to evaluate the flexibility of side-chains. The RMSF and RMSD values ranged from 1.657 Å to 9.472 Å and 1.997 Å to 16.974 Å, respectively. The higher peaks at AA_550-600_, AA_800-850_ and AA_930-950_ in the plot indicate highly flexible regions in the complex (Fig. 9b).

**Fig. 11:**
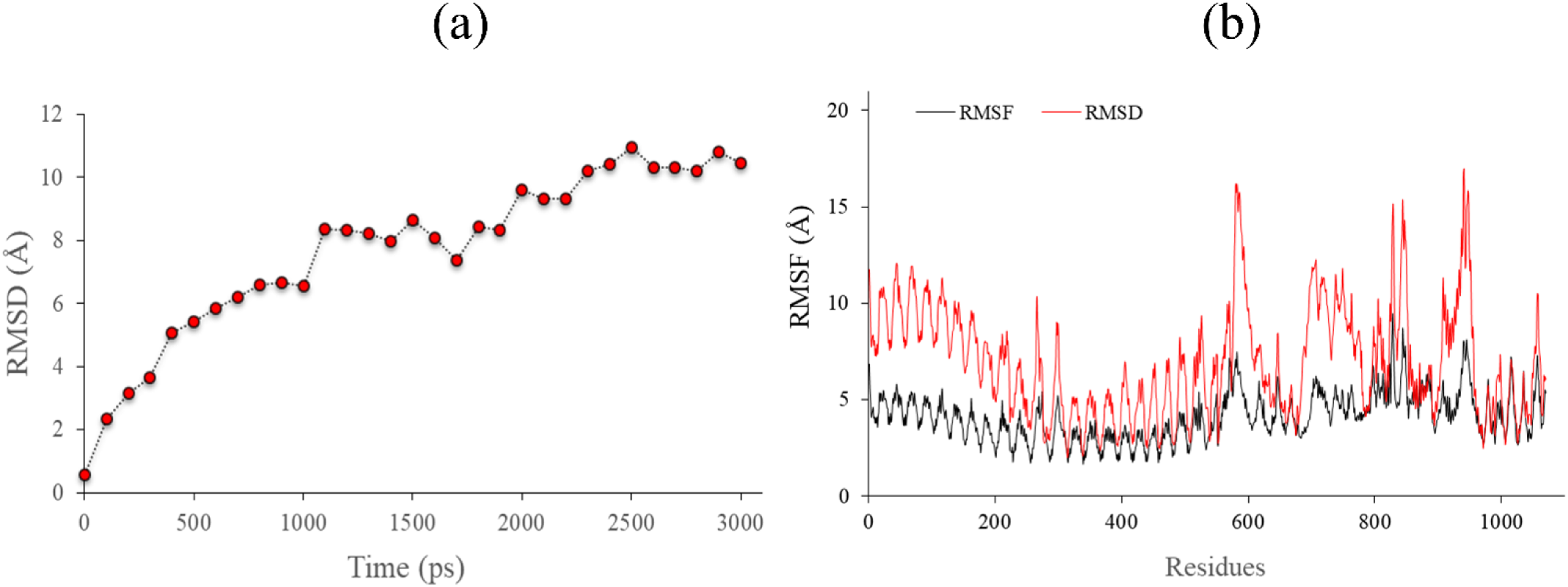
Molecular dynamic simulation of the vaccine-TLR2 complex. (a) RMSD value of the vaccine-TLR2 complex backbone; and (b) RMSF (black) and RMSD (red) values of side-chain residues.

#### 3.2.10. Immune simulation for vaccine efficacy

The computer-generated immune responses were consistent with real-life immune reactions in C-ImmSim server, where secondary and tertiary responses were higher than the primary response (Fig. 12). Both secondary and tertiary responses were characterized by the increased immunoglobulin activity (*i.e.*, IgG1 + IgG2, IgM, and IgG + IgM antibodies) with rapid antigen clearance (Fig. 12a). In addition, higher levels of B-cell activities, especially B isotype IgM and IgG1, was observed with prominent memory cell formation (Fig. 12b,c, Fig. S2). Similarly, the number of active T-cells (both CTL and HTL) was dramatically increased during the secondary and tertiary responses and slowly decreased afterwards (Fig. 12d,e). The development of helper T-cell memory (1800 cells/mm3) was the higher than that of the B-cells (500 cells/mm3, Fig. S2). In addition, macrophage activity was rapidly increased after each exposure and reduced with antigen clearance (Fig. 12f). Furthermore, high levels of IFN-γ, IL-10, and IL-2 were also evident (Fig. 12g). The amount and percentage of Th1 type immune response was higher than Th0 type response (Fig. 12h). Moreover, repeated exposure to 12 vaccine antigens elicited increasing IgG1 and consistent lower activity of IgM and IgG2 immunoglobulins. Also, the amount of IFN-γ and helper T-cell populations was higher throughout exposure (Fig. S2). All of these are relevant for immunity against *M. ulcerans* as we stated earlier. Therefore, the vaccine could be competent enough to control the Buruli ulcer pathogen.

**Fig. 12:**
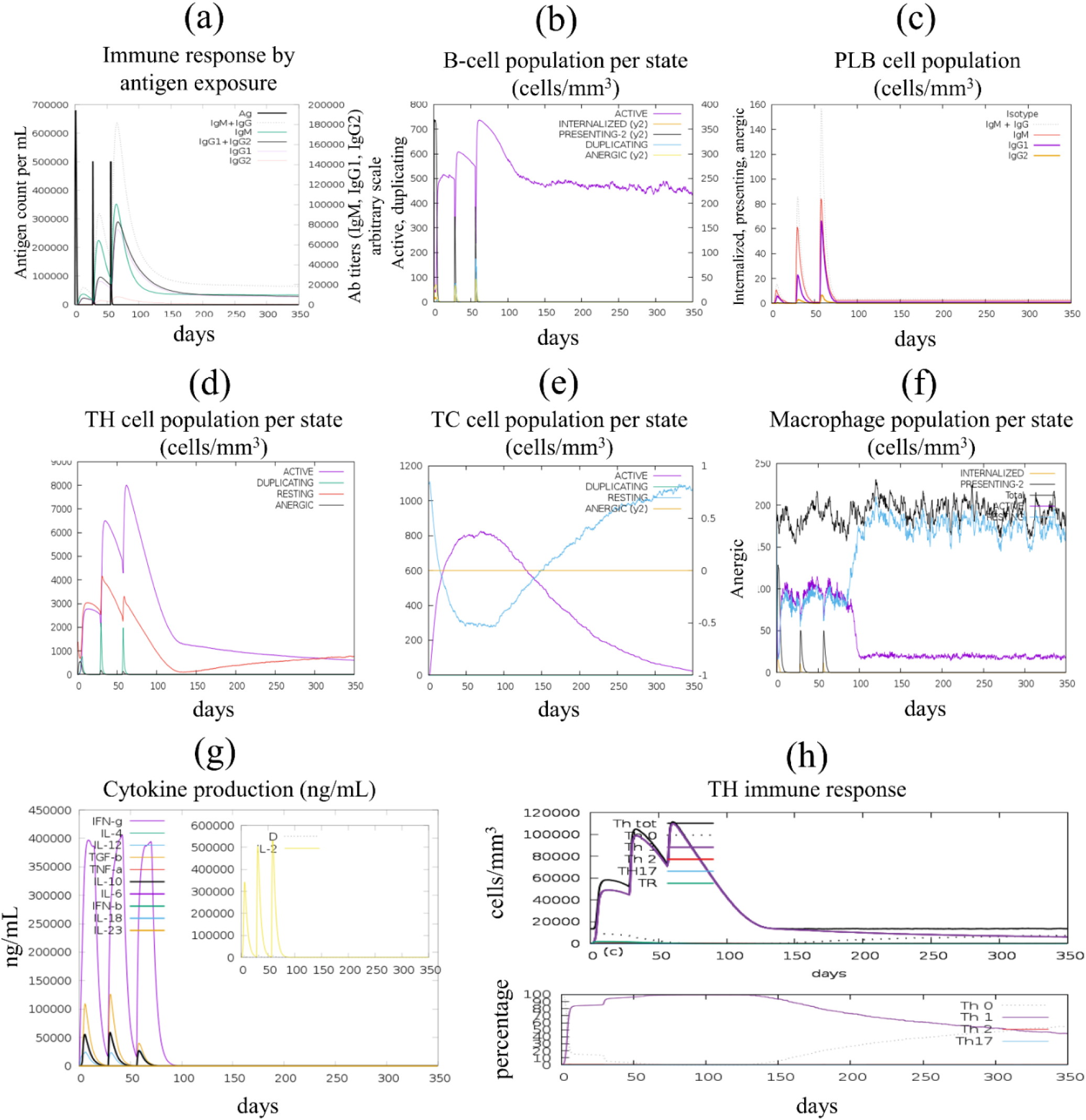
*In silico* generation of immune response using vaccine as antigen. (a) generation of immunoglobulins and B-cell isotypes upon exposure to antigen; (b) amount of active B-cell populations per state; (c) amount of plasma B-lymphocytes and their isotypes per state; (d) state of helper T-cell population during subsequent immune responses; (e) cytotoxic T-cell population per state of antigen exposure; (f) activity of macrophage population in three subsequent immune responses; (g) production of cytokine and interleukins in different states with Simpson index (D); and (h) amount (cells/mm3) and percentage (%) of Th1 mediated immune responses.

#### 3.2.11. In silico cloning within E. coli system

The main purpose of codon optimization and *in silico* cloning was to express the vaccine protein into the *E. coli* host system efficiently. Here, we adapted the *M. ulcerans* codons present in vaccine construct according to the codon usage of *E. coli* K12 strain. The optimized codon sequence was 1560 nucleotides long. The GC-content of the improved DNA sequence was 56.60% and CAI score was 0.986. The CAI score close to 1.0 suggested satisfactory adaptation. Later on, the optimized codon was inserted between the XhoI (158) and NcoI (1724) restriction sites at MCS of the *E. coli* vector pET30a (+) as shown in Fig. 13. The target sequence is also included S-tag and His-tag residues which may help in the affinity-dependent detection and purification process. Thus, the total length of the clone was 6.93 kbp.

**Fig. 13:**
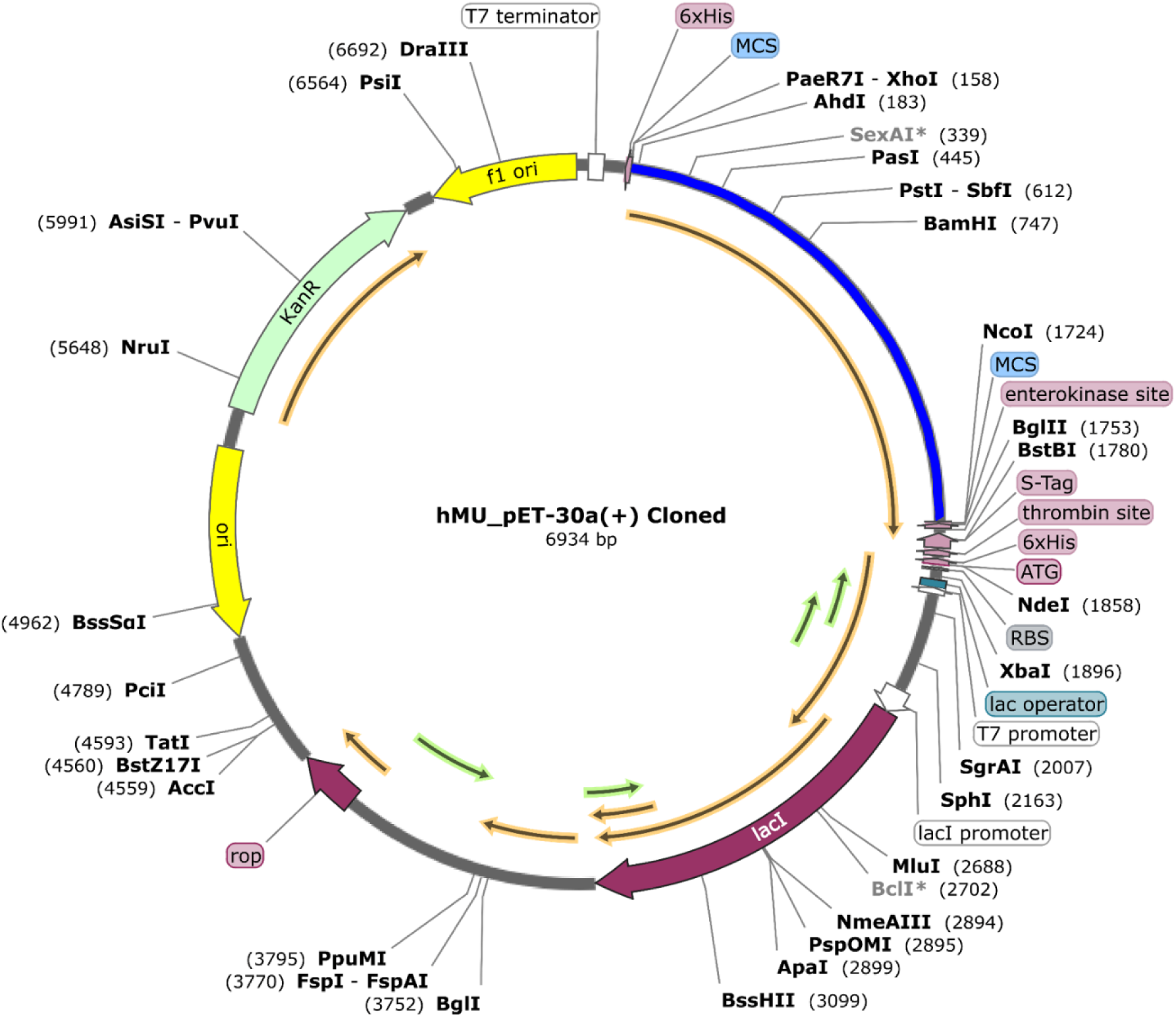
*In silico* cloning of optimized vaccine codon into *E. coli* K12 expression system. Vaccine protein sequence was optimized and inserted into *E. coli* vector pET30a (+) at the position between XhoI (158) and NcoI (1724). The inserted DNA sequence is shown in blue colour while keeping the plasmid grey colour.

## 4. Discussion

Despite being the strongest immune modalities (Nezafat et al. 2016), the development of a vaccine based on immunological experiments is relatively expensive, labor-intensive and time-consuming (Kazi et al. 2018; Plotkin et al. 2017). However, recent advances in immunological bioinformatics have resulted in viable tools and servers that can reduce the time and cost of conventional vaccine development (Kazi et al. 2018). However, the development of effective multi-epitope vaccines remains challenging due to the difficulties in the selection of appropriate antigen candidates, immunodominant epitopes and an efficient delivery system (L. Zhang 2018). Therefore, the prediction of suitable antigenic epitopes of a target protein by immunoinformatic methods is extremely important for designing a multi-epitope vaccine (Yin et al. 2016; Cherryholmes, Stanton, and Disis 2015).

In the current study, we designed a multi-epitope vaccine using the potent T and B-cell epitopes derived from the highest antigenic PE-PGRS family protein of *M. ulcerans*. The PE-PGRS proteins belong to the PE family of conserved mycobacterial protein featured by the presence of a polymorphic GC-rich repeating units (Sampson 2011; Brennan 2017). Although the function(s) of PE-PGRS protein is still remain elusive, emerging evidences support the association of PE/PPE proteins with several steps of mycobacterial pathogenesis (Sampson 2011; Bottai and Brosch 2009). In addition, PE-PGRSs interact with TLR2 receptor to activate the dendritic cells (DCs), macrophages and production of inflammatory cytokines like tumor necrosis factor-alpha (TNF-α) that promote apoptosis and necrosis of host cells (Brennan 2017; Dheenadhayalan, Delogu, and Brennan 2006). Further, they elicit both T-cells and cross-reactive antibodies; and may involve in antigenic variation (Brennan 2017). Moreover, surface or secreted proteins like PE-PGRS protein (Banu et al. 2002) are good targets of immune effector molecules due to their higher exposure to the immune system (Monterrubio-López, González-Y-Merchand, and Ribas-Aparicio 2015). Therefore, proteins belong to PE-PGRS family may be good candidates for vaccine development (Brennan 2017). Furthermore, the number of transmembrane (TM) helix should be considered in vaccine design (He 2014; Dalsass et al. 2019). Proteins with multiple TM helices are usually discouraged because of their inefficiency in cloning, expression, and purification (Monterrubio-López, González-Y-Merchand, and Ribas-Aparicio 2015). The selected PE-PGRS family protein had the highest antigenic score (2.3941) with no transmembrane helix, hence, considered as an appropriate candidate for vaccine design against *M. ulcerans*. Monterrubio-López et al. also found PE-PGRS protein to be highly antigenic and suggested as a potential vaccine candidate against *Mycobacterium tuberculosis* (Monterrubio-López, González-Y-Merchand, and Ribas-Aparicio 2015) However, we didn’t consider mycolactone, disease causing exotoxin, for vaccine design since polyketides are not immunogenic due to their chemical structure. In addition, antibodies against mycolactone were not found either in healthy contacts or in Buruli ulcer sufferers (Huygen et al. 2009).

An ideal multi-epitope vaccine should contain both T- and B-cell epitopes for stimulating a complete network of immune responses (L. Zhang 2018). Keeping it in mind, both types of epitopes were predicted from PE-PGRS protein and evaluated with rigorous analysis. The most potent epitopes were selected based on their immunogenic properties (*i.e.*, antigenicity, allergenicity, toxicity, and cytokine production). The number of HLA binding alleles (5 threshold) was also considered for the broad-spectrum functionality. However, overlapping epitopes were discarded from the final selection for allowing epitopic variability in vaccine protein. Conversely, an exception has been made in case of HTL epitopes since Th1 cells are important for immunity against *M. ulcerans* (Fraga et al. 2011; Nausch et al. 2017; Huygen et al. 2009). Besides, there are considerable differences in their docking affinity, binding residues, and antigenicity. For instance, overlapping HTL epitope (LSVAPDELAAAATEL) provided 11 hydrogen bonds and −6.6 kcal/mol of binding affinity while epitope (SVAPDELAAAATELR) showed 13 hydrogen bonds and −7.4 kcal/mol of binding energy. Moreover, epitope LSVAPDELAAAATEL and SVAPDELAAAATELR were involved with 15 and 18 active site residues where only 6 residues (*i.e.*, Asp55, Asn62 Arg70, Arg77, Asn82, and Ala10) were in common. The selected epitopes were then used for vaccine construction. On the other hand, a *M. ulcerans* derived lipoprotein LprG was added to the vaccine construct as an adjuvant to make vaccine more immunogenic. Therefore, it will mimic the presence of natural pathogen and improve the vaccine’s immunogenicity at the same time. The effectiveness of a vaccine (*e.g.*, BCG) is dependent on the population in which the vaccination is used (Phillips et al. 2015). The rate of Buruli ulcer incidence is the highest in many West African countries such as Benin, Côte d’Ivoire and Ghana (Huygen et al. 2009). Moreover, it has also been reported in tropical countries of Africa, America, Asia and Australia; and non-tropical areas such as Japan and China (Huygen et al. 2009). Importantly, T-cell epitopes included in vaccine construct showed 99.8% of world population coverage and more than 95% coverage in pathogen invaded areas including Buruli ulcer endemic regions, suggesting that the designed vaccine would be effective on majority of the world population.

The vaccine construct was found to be highly antigenic, immunogenic and non-allergenic indicating the potency of epitope vaccine in provoking robust immune responses without producing allergenic reactions. The vaccine construct has 520 amino acid residues including an adjuvant LprG which alone constitutes 241 residues. Chatterjee and co-workers have designed a multi-epitope vaccine which was 568 amino acid long (Chatterjee et al. 2018). On the other hand, Kalita et al. design a multi-epitope vaccine against *Fasciola gigantica* which even had 765 amino acid residues (Kalita et al. 2019). Further, the multi-epitope vaccine against *Schistosoma mansoni, Onchocerca volvulus, Anopheles stephensi*, Leishmania parasites have been designed where the length was 617, 599, 541, 528 amino acid long, respectively (Shey et al. 2019; Rahmani et al. 2019; Khatoon, Pandey, and Prajapati 2017). Moreover, the linkers were added to facilitate the functional preservation of each epitope (~9-15 residues) so that they can function separately after being imported into the human body (Pandey, Bhatt, and Prajapati 2018). Therefore, our results suggest that the size of our vaccine would not be a problem in term of efficacy, stability and expression. The molecular weight of the vaccine protein was 50.78 kDa. According to theoretical PI value, the vaccine is slightly basic which may provide stable interaction in physiological pH range. Furthermore, estimated aliphatic index and instability index scores revealed that the vaccine protein might be stable and thermostable, while the negative grand average hydropathicity score suggested its hydrophilicity; hence, has strong interactions with water molecules. However, peptides with short half-life are the biggest problem in the development of therapeutic proteins (Mathur et al. 2016). In this study, the vaccine protein has 30 hours of half-life in mammalian reticulocytes (*in vitro*) while more than 20 hours in yeast (*in vivo*) and more than 10 hours in *E. coli* cells (*in vivo*), which is satisfactory as stated previously (Khatoon, Pandey, and Prajapati 2017; Pandey, Bhatt, and Prajapati 2018). Moreover, the recombinant proteins should be soluble on overexpression for post-production studies (Magnan, Randall, and Baldi 2009). Importantly, both SolPro and Protein-Sol servers predicted our vaccine protein as highly soluble that ensured easy purification.

For effective transportation of vaccine into the body, strong binding affinity towards the immune receptor (*i.e.*, TLR2) is necessary (Black et al. 2010). In this study, minimal energy (−1372.6) was required by refined vaccine model for locating the binding site of TLR2 receptor appropriately. Moreover, binding free energy (−15.3 kcal/mol) and dissociation rate constant (5.9e-12 M) scores were lower than that of the control that further confirmed the higher binding affinity between the vaccine and TLR2 receptor. Furthermore, the vaccine-TLR2 complex was found to be stable during MD simulation where protein backbone undergone some microscale-changes (~7.67 Å) and mild fluctuations in their side-chain residues that can occur due to the flexible regions (Khatoon, Pandey, and Prajapati 2017; Pandey, Bhatt, and Prajapati 2018). These results are in line with the previous study where stabilization of vaccine-receptor complex was achieved within similar time-scale (M. Ali et al. 2017; Mirza et al. 2016). The hydrogen bond helps in molecular recognition by providing specificity and directionality to the interaction (Hubbard and Kamran Haider 2010). Thus, the presence of multiple hydrogen bonds on the vaccine and TLR2 interaction plane suggested their precise binding. The stability of the vaccine protein was also improved by creating 4 disulfide bridges. Therefore, our data suggest that the vaccine protein will be able to provide stable binding interactions.

A multi-epitope vaccine should be able to elicit both T-cells (*i.e.*, CTL and HTL) and B-cells for effective immune responses (L. Zhang 2018). The control of Buruli ulcer is largely depends on the cellular immunity involving activated macrophages and Th1 type immune response through cytokine production (Gooding et al. 2001; Huygen et al. 2009; Prévot et al. 2004; Yeboah-Manu et al. 2006). Computer simulation of immune response triggered by our vaccine protein showed increased macrophage activity and longer-lasting Th1-mediated immune reactions that are essential for *M. ulcerans* clearance (Hirayama, Iida, and Nakase 2018). The active helper T-cell population was increasingly higher in secondary and tertiary responses compared to the primary response. In addition to T-cells, antibodies also provide protection against extracellular *M. ulcerans* (Huygen et al. 2009). Interestingly, the multi-epitope vaccine has 12 conformational B-cell epitopes throughout its three protein domains. Furthermore, we observed a plethora of active immunoglobulins *i.e.*, IgM and IgG and their isotypes which could contribute to isotype switching. Therefore, the simulated immune response was characterized by higher levels of helper T-cells and B-cells activities. These results coincide with the previous findings where PE-PGRS proteins from *M. tuberculosis* were used to trigger immune response *in silico* (Monterrubio-López, González-Y-Merchand, and Ribas-Aparicio 2015). As in tuberculosis and leprosy, the macrophage activating cytokine IFN-γ seems to play a pivotal role in the control of *M. ulcerans* infection. However, whole *M. ulcerans* had shown reduced amount of IFN-γ production upon *in vitro* stimulation (Gooding et al. 2001; Huygen et al. 2009; Prévot et al. 2004). In a study on mouse model, mycolactone has been observed to attack mononuclear cells (*i.e.*, lymphocytes, monocytes, DCs and macrophages) and thus interfere with cellular immunity in the immunosuppressed host (Hong et al. 2008). Thus, mycolactone affects human lymphocytes and macrophages to inhibit the production of IL-2 and TNF-α, respectively (Coutanceau et al. 2005; Torrado et al. 2007). Besides, it also interrupts DCs to prime cell-mediated immunity and produces chemotactic inflammatory signals (Coutanceau et al. 2007). Importantly, IFN-γ production by newly designed vaccine was the highest among other cytokines. Furthermore, prominent activity of IL-2 and IL-10 was also observed. However, the peptide vaccine was not able to induce TNF-α production through macrophage activation. In endemic or pathogen invaded regions, the simulated immune response showed rapid antigen clearance and strong protection against *M. ulcerans* (Fig. S2). Moreover, the negligible Simpson index (D) suggests a diverse immune response (Rapin et al. 2010) which is plausible since the vaccine chimera contained multiple B and T-cell epitopes.

The translational efficiency of foreign genes may differ within the host system due to the incompatibility of mRNA codons which necessitates the optimization of codon for higher expression (Pandey, Bhatt, and Prajapati 2018). Satisfactorily, the obtained CAI value (0.986) was very close to 1.0 and GC contents (56.60%) was also within the optimal limit (30-70%) indicating possible higher expression within *E. coli* K12 system (Shey et al. 2019; Morla, Makhija, and Kumar 2016). As a suggestion for prospective vaccine synthesis, *in silico* cloning was also performed using pET30a(+) vector. This plasmid has both His-tag and S-tag as fusion partners that facilitate the ease of purification (Hengen 1995; Raines et al. 2000). Besides, S-tag sequence increases protein stability by providing its abundance of charged and polar residues (Raines et al. 2000). Moreover, *in silico* cloning with pET30a(+) vector was described earlier (Shey et al. 2019).

## 5. Conclusions

This study was performed to design an effective vaccine against *M. ulcerans* as a Buruli ulcer prophylaxis using integrated vaccinomics approach. The designed vaccine has antigenic, immunogenic, non-allergenic and non-toxic properties with higher binding affinity towards TLR2 immune receptor. Moreover, the vaccine was found stable at physiological pH range. In addition, disulfide bridges were created for structural improvement and *in silico* cloning was performed to ensure good expression in widely used *E. coli* K12 system. Furthermore, the simulated immune response was diverse and characterized by cellular and humoral immune responses with efficient memory cell formation. However, the current study is the sole outcome of an integrated vaccinomics approach; therefore, it needs experimental validation to prove the efficacy and safeness of the designed vaccine which may include the synthesis of vaccine protein followed by the rigorous assessment *in vitro* and *in vivo*.

## Author Contributions

Study design: ZN and UKA; Immunoinformatic analysis and draft manuscript: ZN; Critical revision: MMK and MKS; Study supervision: UKA. All authors approved the final manuscript.

## Abbreviations

ΔG: binding free energy
3D: three-dimensional
AI: aliphatic index
BCG: Bacillus Calmette-Guérin
CAI: codon adaptation index
CTL: cytotoxic T-lymphocyte
DC: dendritic cells
GC: guanine-cytosine
GRAVY: grand average of hydropathicity
HLA: human leukocytes antigen
HTL: helper T-lymphocyte
IFN-γ: interferon- gamma
IgG: immunoglobulin G
IgM: immunoglobulin M
II: instability index
IL: interleukin
IMG/M: Integrated Microbial Genomes & Microbiomes
Kd: dissociation constant
LBL: linear B-lymphocyte
MCS: multiple cloning site
MD: molecular dynamics
MHC: major histocompatibility complex
MW: molecular weight
NCBI: national center for biotechnology information
PE-PGRS: proline-glutamate polymorphic GC-rich sequence
RMSD: root mean square deviation
RMSF: root mean square fluctuation
SSL3: *Staphylococca*l superantigen-like protein 3
Theoretical PI: theoretical isoelectric point
TLR: toll-like receptor
TM: transmembrane
TNF-α: tumor necrosis factor-alpha

