## Supplementary Tables and Figures for "Structural Basis and Designing of Peptide Vaccine using PE-PGRS Family Protein of *Mycobacterium ulcerans* – An Integrated Vaccinomics Approach"

**List of Supplementary Tables**

**Table S1:** List of predicted 14 antigenic & immunogenic CTL epitopes having ≥5 MHC-I binding alleles

| **Supertype** | **CTL Epitope** | **C-Score** | **Antigenicity** | **No. of Alleles** | **Immunogenicity** | **Allergenicity** | **Toxicity** |
| --- | --- | --- | --- | --- | --- | --- | --- |
| B7 | AAAVPTTAL | 1.5339 | 0.5564 | 12 | 0.1356 | No | Non-toxin |
| B7 | AADEVSAAI | 0.7703 | 0.6396 | 10 | 0.0483 | Yes | Non-toxin |
| A3 | AANGAVGSR | 0.5103 | 2.1085 | 5 | 0.0409 | No | Non-toxin |
| B62 | AGGTGGLLF | 0.6031 | 1.616 | 2 | 0.09912 | Yes | Non-toxin |
| B62 | ALAAGGASY | 1.5616 | 0.8068 | 15 | 0.05333 | Yes | Non-toxin |
| B7 | APNGGAGGL | 1.4081 | 1.8634 | 7 | 0.15023 | Yes | Non-toxin |
| B44 | DEVSAAIAA | 0.519 | 0.5864 | 6 | 0.05704 | Yes | Non-toxin |
| A26 | EVSAAIAAL | 1.3825 | 0.575 | 15 | 0.20493 | Yes | Non-toxin |
| B7 | GARGADGLL | 0.661 | 1.2613 | 4 | 0.132 | No | Non-toxin |
| A1 | GGDGGHGAY | 0.7632 | 2.6448 | 7 | 0.15621 | No | Non-toxin |
| B7 | GIRGADGVL | 0.6497 | 1.0974 | 3 | 0.1626 | No | Non-toxin |
| B62 | GQGGGAGTL | 1.0216 | 3.2446 | 5 | 0.16621 | Yes | Non-toxin |
| B27 | GRGGGTGGA | 0.6368 | 4.2972 | 1 | 0.16304 | No | Non-toxin |
| B62 | GSGGRGGLF | 0.8373 | 2.0711 | 4 | 0.14952 | Yes | Non-toxin |
| B39 | HGARGADGL | 0.7019 | 1.4358 | 2 | 0.17313 | Yes | Non-toxin |
| A3 | LLDGEAGSK | 1.0037 | 1.9529 | 4 | 0.10757 | Yes | Non-toxin |
| A24 | LWPAAADEV | 0.5785 | 0.9216 | 5 | 0.18792 | Yes | Non-toxin |
| B62 | LYGNGGNGY | 0.7471 | 2.0088 | 7 | 0.08373 | Yes | Non-toxin |
| B7 | SPAALDGGA | 0.505 | 0.7087 | 4 | 0.11055 | No | Non-toxin |
| A1 | SSDAANGAV | 2.1709 | 1.3153 | 7 | 0.13004 | Yes | Non-toxin |
| B58 | VSAAIAALF | 1.7464 | 0.6021 | 16 | 0.24504 | No | Non-toxin |
| B39 | YLSVAPDEL | 1.2856 | 1.0078 | 17 | 0.09272 | No | Non-toxin |
| B7 | AAAVPTTAL | 1.5339 | 0.5564 | 12 | 0.1356 | No | Non-toxin |
| B7 | AADEVSAAI | 0.7703 | 0.6396 | 10 | 0.0483 | Yes | Non-toxin |
| A3 | AANGAVGSR | 0.5103 | 2.1085 | 5 | 0.0409 | No | Non-toxin |
| B62 | AGGTGGLLF | 0.6031 | 1.616 | 2 | 0.09912 | Yes | Non-toxin |
| B62 | ALAAGGASY | 1.5616 | 0.8068 | 15 | 0.05333 | Yes | Non-toxin |
| B7 | APNGGAGGL | 1.4081 | 1.8634 | 7 | 0.15023 | Yes | Non-toxin |

**Table S2:** List of predicted 53 HTL epitopes with ≥5 MHC-II binding alleles

| **HTL Epitope** | **No. of MHC-II Alleles** | **IFN-γ Prediction** | **IL-4 Prediction** | **IL-10 Prediction** |
| --- | --- | --- | --- | --- |
| AAAATELRGVGSIVS | 23 | POSITIVE | Non IL4 inducer | IL10 non-inducer |
| AAATELRGVGSIVST | 23 | POSITIVE | Non IL4 inducer | IL10 inducer |
| AATELRGVGSIVSTV | 24 | POSITIVE | Non IL4 inducer | IL10 non-inducer |
| AAYHDQFVRALAAGG | 14 | NEGATIVE | IL4 inducer | IL10 non-inducer |
| AQQYQSLSVQVAAYH | 10 | NEGATIVE | IL4 inducer | IL10 non-inducer |
| ASPLEQLLDLINLPS | 8 | NEGATIVE | Non IL4 inducer | IL10 inducer |
| ATELRGVGSIVSTVN | 25 | POSITIVE | Non IL4 inducer | IL10 inducer |
| AYHDQFVRALAAGGA | 19 | POSITIVE | IL4 inducer | IL10 non-inducer |
| DEVSAAIAALFSSHA | 5 | NEGATIVE | Non IL4 inducer | IL10 non-inducer |
| DQFVRALAAGGASYS | 20 | POSITIVE | Non IL4 inducer | IL10 non-inducer |
| ELRGVGSIVSTVNAA | 25 | POSITIVE | Non IL4 inducer | IL10 non-inducer |
| EQLLDLINLPSQTLL | 12 | NEGATIVE | Non IL4 inducer | IL10 inducer |
| EVSAAIAALFSSHAQ | 6 | POSITIVE | Non IL4 inducer | IL10 non-inducer |
| FVRALAAGGASYSLA | 16 | POSITIVE | Non IL4 inducer | IL10 non-inducer |
| GGGGSQVGSDGVGGA | 6 | POSITIVE | IL4 inducer | IL10 non-inducer |
| GGGSQVGSDGVGGAG | 6 | POSITIVE | Non IL4 inducer | IL10 non-inducer |
| GGSQVGSDGVGGAGG | 6 | POSITIVE | Non IL4 inducer | IL10 non-inducer |
| GSIVSTVNAAAAVPT | 6 | POSITIVE | IL4 inducer | IL10 non-inducer |
| GSQVGSDGVGGAGGA | 6 | POSITIVE | Non IL4 inducer | IL10 non-inducer |
| GVGSIVSTVNAAAAV | 10 | POSITIVE | IL4 inducer | IL10 non-inducer |
| HAQQYQSLSVQVAAY | 10 | POSITIVE | IL4 inducer | IL10 inducer |
| HDQFVRALAAGGASY | 20 | NEGATIVE | IL4 inducer | IL10 non-inducer |
| IVSTVNAAAAVPTTA | 10 | POSITIVE | Non IL4 inducer | IL10 non-inducer |
| LDLINLPSQTLLGRP | 6 | NEGATIVE | Non IL4 inducer | IL10 inducer |
| LEQLLDLINLPSQTL | 11 | NEGATIVE | Non IL4 inducer | IL10 inducer |
| LLDLINLPSQTLLGR | 12 | POSITIVE | Non IL4 inducer | IL10 inducer |
| LRGVGSIVSTVNAAA | 27 | POSITIVE | Non IL4 inducer | IL10 non-inducer |
| LSVAPDELAAAATEL | 6 | POSITIVE | IL4 inducer | IL10 non-inducer |
| MWYLSVAPDELAAAA | 13 | POSITIVE | Non IL4 inducer | IL10 non-inducer |
| PLEQLLDLINLPSQT | 8 | NEGATIVE | Non IL4 inducer | IL10 inducer |
| QFVRALAAGGASYSL | 17 | POSITIVE | Non IL4 inducer | IL10 non-inducer |
| QLLDLINLPSQTLLG | 12 | NEGATIVE | Non IL4 inducer | IL10 inducer |
| QQYQSLSVQVAAYHD | 9 | NEGATIVE | IL4 inducer | IL10 non-inducer |
| QVGSDGVGGAGGAGG | 6 | POSITIVE | Non IL4 inducer | IL10 non-inducer |
| QYQSLSVQVAAYHDQ | 8 | NEGATIVE | IL4 inducer | IL10 non-inducer |
| RGVGSIVSTVNAAAA | 10 | POSITIVE | IL4 inducer | IL10 non-inducer |
| SAAIAALFSSHAQQY | 6 | POSITIVE | Non IL4 inducer | IL10 inducer |
| SHAQQYQSLSVQVAA | 9 | POSITIVE | IL4 inducer | IL10 inducer |
| SIVSTVNAAAAVPTT | 11 | POSITIVE | IL4 inducer | IL10 non-inducer |
| SPLEQLLDLINLPSQ | 8 | NEGATIVE | Non IL4 inducer | IL10 inducer |
| SQVGSDGVGGAGGAG | 6 | POSITIVE | Non IL4 inducer | IL10 non-inducer |
| SSHAQQYQSLSVQVA | 9 | NEGATIVE | IL4 inducer | IL10 inducer |
| SVAPDELAAAATELR | 6 | POSITIVE | IL4 inducer | IL10 non-inducer |
| TELRGVGSIVSTVNA | 25 | POSITIVE | Non IL4 inducer | IL10 inducer |
| VAPDELAAAATELRG | 6 | POSITIVE | IL4 inducer | IL10 non-inducer |
| VGSDGVGGAGGAGGT | 6 | POSITIVE | Non IL4 inducer | IL10 non-inducer |
| VGSIVSTVNAAAAVP | 10 | POSITIVE | IL4 inducer | IL10 non-inducer |
| VRALAAGGASYSLAE | 6 | POSITIVE | Non IL4 inducer | IL10 non-inducer |
| VSAAIAALFSSHAQQ | 5 | POSITIVE | Non IL4 inducer | IL10 non-inducer |
| WYLSVAPDELAAAAT | 6 | POSITIVE | Non IL4 inducer | IL10 non-inducer |
| YHDQFVRALAAGGAS | 20 | POSITIVE | IL4 inducer | IL10 non-inducer |
| YLSVAPDELAAAATE | 7 | POSITIVE | Non IL4 inducer | IL10 non-inducer |
| YQSLSVQVAAYHDQF | 8 | NEGATIVE | Non IL4 inducer | IL10 non-inducer |
| AAAATELRGVGSIVS | 23 | POSITIVE | Non IL4 inducer | IL10 non-inducer |
| AAATELRGVGSIVST | 23 | POSITIVE | Non IL4 inducer | IL10 inducer |

**Table S3:** Potential LBL epitopes from *M. ulcerans* derived PE-PGRS family protein (PS>0.75)

| **LBL Epitope** | **Probability score** | **Antigenicity** | **Allergenicity** | **Toxicity** |
| --- | --- | --- | --- | --- |
| GNGGRGGDGIRG | 0.8431 | 3.7111 | No | No |
| GGNGGRGGDGIR | 0.8392 | 4.0397 | No | No |
| LLYGNGGNGYDN | 0.8293 | 1.0946 | No | No |
| GAGGAGGTGGAG | 0.8285 | 4.2749 | No | No |
| GGAGGAGGTGGA | 0.8285 | 4.1151 | No | No |
| GNGGNGYDNSAS | 0.8095 | 2.087 | No | No |
| TPGGSGGTGGGG | 0.7969 | 4.5182 | No | No |
| GTPGGSGGTGGG | 0.7969 | 3.8817 | No | No |
| AGGAGGAGGGGS | 0.7945 | 4.3426 | No | No |
| LYGNGGNGYDNS | 0.7897 | 1.4508 | No | No |
| DGESGTAGGNGG | 0.784 | 3.4194 | No | No |
| HGGIGGNGARGA | 0.779 | 2.5634 | No | No |
| SGGAGGAGGAGG | 0.7749 | 3.9851 | No | No |
| GSGGTGGGGSAG | 0.7746 | 4.7979 | No | No |
| DGADGTASAPNG | 0.7702 | 1.4195 | No | No |
| GHGGIGGNGARG | 0.7681 | 2.8838 | No | No |
| YGNGGNGYDNSA | 0.7658 | 1.7089 | No | No |
| GNGARGADGVGD | 0.7633 | 2.5455 | No | No |
| GADGTASAPNGG | 0.7589 | 1.5176 | No | No |
| GGGNGGRGGDGI | 0.7535 | 4.2413 | No | No |
| GDGADGTASAPN | 0.752 | 2.0047 | No | No |
| GLLYGNGGNGYD | 0.751 | 0.8936 | No | No |

**Table S4:** Predicted population coverage by T-cell epitopes selected for vaccine construction

| **MHC class I** | | **MHC class II** | | **MHC class I and II (combined)** | |
| --- | --- | --- | --- | --- | --- |
| **Regions** | **Coverage** | **Regions** | **Coverage** | **Regions** | **Coverage** |
| American Samoa | 95.22% | Algeria | 45.71% | Algeria | 45.71% |
| American Samoa Polynesian | 95.22% | Algeria Arab | 45.71% | Algeria Arab | 45.71% |
| Argentina | 98.70% | Argentina | 49.23% | American Samoa | 95.22% |
| Argentina Amerindian | 98.70% | Argentina Amerindian | 47.18% | American Samoa Polynesian | 95.22% |
| Australia | 96.19% | Argentina Caucasoid | 51.43% | Argentina | 99.34% |
| Australia Australian Aborigines | 93.76% | Australia | 12.52% | Argentina Amerindian | 99.31% |
| Australia Caucasoid | 99.45% | Australia Australian Aborigines | 12.52% | Argentina Caucasoid | 51.43% |
| Austria | 94.52% | Austria | 61.44% | Australia | 96.67% |
| Austria Caucasoid | 94.52% | Austria Caucasoid | 61.44% | Australia Australian Aborigines | 94.54% |
| Belgium | 93.83% | Belarus | 40.76% | Australia Caucasoid | 99.45% |
| Belgium Caucasoid | 93.83% | Belarus Caucasoid | 40.76% | Austria | 97.89% |
| Brazil | 97.53% | Belgium | 56.18% | Austria Caucasoid | 97.89% |
| Brazil Amerindian | 98.40% | Belgium Caucasoid | 56.18% | Belarus | 40.76% |
| Brazil Caucasoid | 95.54% | Bolivia | 84.16% | Belarus Caucasoid | 40.76% |
| Brazil Mixed | 98.42% | Bolivia Amerindian | 84.16% | Belgium | 97.30% |
| Bulgaria | 99.19% | Borneo | 4.74% | Belgium Caucasoid | 97.30% |
| Bulgaria Caucasoid | 98.53% | Borneo Austronesian | 4.74% | Bolivia | 84.16% |
| Bulgaria Other | 100.00% | Brazil | 61.36% | Bolivia Amerindian | 84.16% |
| Burkina Faso | 74.84% | Brazil Amerindian | 63.53% | Borneo | 4.74% |
| Burkina Faso Black | 74.84% | Brazil Caucasoid | 66.89% | Borneo Austronesian | 4.74% |
| Cameroon | 97.46% | Brazil Mixed | 59.41% | Brazil | 99.05% |
| Cameroon Black | 97.46% | Brazil Mulatto | 52.67% | Brazil Amerindian | 99.42% |
| Cape Verde | 97.42% | Bulgaria | 70.30% | Brazil Caucasoid | 98.52% |
| Cape Verde Black | 97.42% | Bulgaria Caucasoid | 70.30% | Brazil Mixed | 99.36% |
| Central Africa | 95.19% | Cameroon | 32.60% | Brazil Mulatto | 52.67% |
| Central African Republic | 36.80% | Cameroon Black | 32.60% | Bulgaria | 99.76% |
| Central African Republic Black | 36.80% | Canada | 28.19% | Bulgaria Caucasoid | 99.56% |
| Central America | 7.20% | Canada Amerindian | 28.19% | Bulgaria Other | 100.00% |
| Chile | 99.80% | Cape Verde | 59.93% | Burkina Faso | 74.84% |
| Chile Amerindian | 99.98% | Cape Verde Black | 59.93% | Burkina Faso Black | 74.84% |
| Chile Mixed | 94.68% | Central Africa | 40.38% | Cameroon | 98.29% |
| China | 96.44% | Central African Republic | 5.91% | Cameroon Black | 98.29% |
| China Oriental | 96.44% | Central African Republic Black | 5.91% | Canada | 28.19% |
| Colombia | 70.77% | Central America | 44.28% | Canada Amerindian | 28.19% |
| Colombia Amerindian | 53.00% | Chile | 55.14% | Cape Verde | 98.96% |
| Colombia Black | 51.66% | Chile Amerindian | 72.65% | Cape Verde Black | 98.96% |
| Colombia Mestizo | 80.12% | Chile Mixed | 37.24% | Central Africa | 97.13% |
| Croatia | 97.57% | China | 41.21% | Central African Republic | 40.53% |
| Croatia Caucasoid | 97.57% | China Oriental | 41.21% | Central African Republic Black | 40.53% |
| Cuba | 96.15% | Colombia | 49.88% | Central America | 48.29% |
| Cuba Caucasoid | 96.45% | Colombia Amerindian | 57.52% | Chile | 99.91% |
| Cuba Mulatto | 95.82% | Colombia Black | 48.08% | Chile Amerindian | 99.99% |
| Czech Republic | 98.05% | Colombia Mestizo | 38.53% | Chile Mixed | 96.66% |
| Czech Republic Caucasoid | 98.05% | Congo | 45.61% | China | 97.90% |
| East Africa | 94.35% | Congo Black | 45.61% | China Oriental | 97.90% |
| East Asia | 99.48% | Cook Islands | 58.98% | Colombia | 85.35% |
| Ecuador | 99.66% | Cook Islands Polynesian | 58.98% | Colombia Amerindian | 80.03% |
| Ecuador Amerindian | 99.66% | Costa Rica | 17.19% | Colombia Black | 74.90% |
| England | 99.63% | Costa Rica Mestizo | 17.19% | Colombia Mestizo | 87.78% |
| England Caucasoid | 99.75% | Croatia | 69.74% | Congo | 45.61% |
| England Jew | 47.29% | Croatia Caucasoid | 69.74% | Congo Black | 45.61% |
| Equatorial Guinea | 1.00% | Cuba | 43.30% | Cook Islands | 58.98% |
| Equatorial Guinea Black | 1.00% | Cuba Mixed | 43.30% | Cook Islands Polynesian | 58.98% |
| Europe | 99.62% | Czech Republic | 54.41% | Costa Rica | 17.19% |
| Finland | 99.33% | Czech Republic Caucasoid | 55.04% | Costa Rica Mestizo | 17.19% |
| Finland Caucasoid | 99.33% | Czech Republic Other | 49.26% | Croatia | 99.26% |
| France | 99.23% | Denmark | 61.44% | Croatia Caucasoid | 99.26% |
| France Caucasoid | 99.23% | Denmark Caucasoid | 61.44% | Cuba | 97.81% |
| Georgia | 98.16% | East Africa | 54.57% | Cuba Caucasoid | 96.45% |
| Georgia Caucasoid | 98.70% | East Asia | 63.84% | Cuba Mixed | 43.30% |
| Georgia Kurd | 97.22% | Ecuador | 68.09% | Cuba Mulatto | 95.82% |
| Germany | 99.43% | Ecuador Amerindian | 68.09% | Czech Republic | 99.11% |
| Germany Caucasoid | 99.43% | England | 60.86% | Czech Republic Caucasoid | 99.12% |
| Guatemala | 7.20% | England Caucasoid | 60.86% | Czech Republic Other | 49.26% |
| Guatemala Amerindian | 7.20% | Equatorial Guinea | 36.96% | Denmark | 61.44% |
| Guinea-Bissau | 92.41% | Equatorial Guinea Black | 36.96% | Denmark Caucasoid | 61.44% |
| Guinea-Bissau Black | 92.41% | Ethiopia | 46.99% | East Africa | 97.43% |
| Hong Kong | 95.29% | Ethiopia Black | 46.99% | East Asia | 99.81% |
| Hong Kong Oriental | 95.29% | Europe | 57.44% | Ecuador | 99.89% |
| India | 99.42% | Fiji | 76.62% | Ecuador Amerindian | 99.89% |
| India Asian | 99.42% | Fiji Melanesian | 76.62% | England | 99.85% |
| Indonesia | 81.47% | Finland | 43.00% | England Caucasoid | 99.90% |
| Indonesia Austronesian | 81.47% | Finland Caucasoid | 43.00% | England Jew | 47.29% |
| Iran | 97.62% | France | 63.85% | Equatorial Guinea | 37.59% |
| Iran Persian | 97.62% | France Caucasoid | 63.85% | Equatorial Guinea Black | 37.59% |
| Ireland Northern | 99.68% | Gabon | 33.74% | Ethiopia | 46.99% |
| Ireland Northern Caucasoid | 99.68% | Gabon Black | 33.74% | Ethiopia Black | 46.99% |
| Ireland South | 99.68% | Georgia | 59.41% | Europe | 99.84% |
| Ireland South Caucasoid | 99.68% | Georgia Caucasoid | 59.41% | Fiji | 76.62% |
| Israel | 94.20% | Germany | 61.12% | Fiji Melanesian | 76.62% |
| Israel Arab | 97.23% | Germany Caucasoid | 61.12% | Finland | 99.62% |
| Israel Jew | 95.99% | Greece | 65.57% | Finland Caucasoid | 99.62% |
| Italy | 98.55% | Greece Caucasoid | 65.57% | France | 99.72% |
| Italy Caucasoid | 98.55% | Guatemala | 50.44% | France Caucasoid | 99.72% |
| Ivory Coast | 60.97% | Guatemala Amerindian | 50.44% | Gabon | 33.74% |
| Ivory Coast Black | 60.97% | Guinea-Bissau | 52.25% | Gabon Black | 33.74% |
| Japan | 99.30% | Guinea-Bissau Black | 52.25% | Georgia | 99.25% |
| Japan Oriental | 99.30% | India | 35.75% | Georgia Caucasoid | 99.47% |
| Jordan | 86.50% | India Asian | 35.75% | Georgia Kurd | 97.22% |
| Jordan Arab | 86.50% | Indonesia | 16.63% | Germany | 99.78% |
| Kenya | 94.91% | Indonesia Austronesian | 16.63% | Germany Caucasoid | 99.78% |
| Kenya Black | 94.91% | Iran | 45.54% | Greece | 65.57% |
| Korea; South | 99.40% | Iran Kurd | 31.94% | Greece Caucasoid | 65.57% |
| Korea; South Oriental | 99.40% | Iran Persian | 49.49% | Guatemala | 54.01% |
| Lebanon | 74.80% | Ireland Northern | 53.05% | Guatemala Amerindian | 54.01% |
| Lebanon Mixed | 74.80% | Ireland Northern Caucasoid | 53.05% | Guinea-Bissau | 96.38% |
| Macedonia | 35.67% | Ireland South | 49.31% | Guinea-Bissau Black | 96.38% |
| Macedonia Caucasoid | 35.67% | Ireland South Caucasoid | 49.31% | Hong Kong | 95.29% |
| Malaysia | 81.70% | Israel | 60.24% | Hong Kong Oriental | 95.29% |
| Malaysia Austronesian | 70.71% | Israel Arab | 58.30% | India | 99.63% |
| Malaysia Oriental | 87.50% | Israel Jew | 60.70% | India Asian | 99.63% |
| Mali | 96.41% | Italy | 42.77% | Indonesia | 84.55% |
| Mali Black | 96.41% | Italy Caucasoid | 42.77% | Indonesia Austronesian | 84.55% |
| Martinique | 24.31% | Jamaica | 8.22% | Iran | 98.71% |
| Martinique Black | 24.31% | Jamaica Black | 8.22% | Iran Kurd | 31.94% |
| Mexico | 99.82% | Japan | 63.40% | Iran Persian | 98.80% |
| Mexico Amerindian | 99.93% | Japan Oriental | 63.40% | Ireland Northern | 99.85% |
| Mexico Mestizo | 98.80% | Jordan | 43.75% | Ireland Northern Caucasoid | 99.85% |
| Mongolia | 88.93% | Jordan Arab | 43.75% | Ireland South | 99.84% |
| Mongolia Oriental | 88.93% | Kiribati | 3.17% | Ireland South Caucasoid | 99.84% |
| Morocco | 98.77% | Kiribati Micronesian | 3.17% | Israel | 97.69% |
| Morocco Arab | 98.81% | Korea; South | 56.58% | Israel Arab | 98.85% |
| Morocco Caucasoid | 98.79% | Korea; South Oriental | 56.58% | Israel Jew | 98.42% |
| New Caledonia | 98.65% | Lebanon | 78.24% | Italy | 99.17% |
| New Caledonia Melanesian | 98.65% | Lebanon Arab | 78.24% | Italy Caucasoid | 99.17% |
| North Africa | 96.60% | Macedonia | 63.08% | Ivory Coast | 60.97% |
| North America | 99.76% | Macedonia Caucasoid | 63.08% | Ivory Coast Black | 60.97% |
| Northeast Asia | 96.55% | Malaysia | 35.27% | Jamaica | 8.22% |
| Oceania | 96.65% | Malaysia Austronesian | 32.46% | Jamaica Black | 8.22% |
| Oman | 98.07% | Malaysia Oriental | 43.84% | Japan | 99.74% |
| Oman Arab | 98.07% | Martinique | 62.59% | Japan Oriental | 99.74% |
| Pakistan | 94.40% | Martinique Black | 62.59% | Jordan | 92.40% |
| Pakistan Asian | 93.16% | Mexico | 55.54% | Jordan Arab | 92.40% |
| Pakistan Mixed | 96.57% | Mexico Amerindian | 52.10% | Kenya | 94.91% |
| Papua New Guinea | 99.07% | Mexico Mestizo | 60.98% | Kenya Black | 94.91% |
| Papua New Guinea Melanesian | 99.07% | Mongolia | 58.82% | Kiribati | 3.17% |
| Peru | 100.00% | Mongolia Oriental | 58.82% | Kiribati Micronesian | 3.17% |
| Peru Amerindian | 100.00% | Morocco | 41.29% | Korea; South | 99.74% |
| Peru Mestizo | 1.99% | Morocco Arab | 44.15% | Korea; South Oriental | 99.74% |
| Philippines | 98.12% | Morocco Caucasoid | 37.27% | Lebanon | 94.52% |
| Philippines Austronesian | 98.12% | Nauru | 23.76% | Lebanon Arab | 78.24% |
| Poland | 99.09% | Nauru Micronesian | 23.76% | Lebanon Mixed | 74.80% |
| Poland Caucasoid | 99.09% | Netherlands | 60.75% | Macedonia | 76.25% |
| Portugal | 98.58% | Netherlands Caucasoid | 60.75% | Macedonia Caucasoid | 76.25% |
| Portugal Caucasoid | 98.58% | New Caledonia | 69.32% | Malaysia | 88.16% |
| Romania | 96.93% | New Caledonia Melanesian | 69.32% | Malaysia Austronesian | 80.22% |
| Romania Caucasoid | 96.93% | New Zealand | 47.99% | Malaysia Oriental | 92.98% |
| Russia | 98.94% | New Zealand Polynesian | 47.99% | Mali | 96.41% |
| Russia Caucasoid | 70.84% | Niue | 38.38% | Mali Black | 96.41% |
| Russia Mixed | 66.31% | Niue Polynesian | 38.38% | Martinique | 71.68% |
| Russia Other | 99.47% | North Africa | 49.96% | Martinique Black | 71.68% |
| Russia Siberian | 98.95% | North America | 60.62% | Mexico | 99.92% |
| Rwanda | 43.89% | Northeast Asia | 41.21% | Mexico Amerindian | 99.97% |
| Rwanda Black | 43.89% | Norway | 57.75% | Mexico Mestizo | 99.53% |
| Sao Tome and Principe | 91.45% | Norway Caucasoid | 57.75% | Mongolia | 95.44% |
| Sao Tome and Principe Black | 91.45% | Oceania | 43.73% | Mongolia Oriental | 95.44% |
| Saudi Arabia | 98.47% | Papua New Guinea | 54.37% | Morocco | 99.28% |
| Saudi Arabia Arab | 98.47% | Papua New Guinea Melanesian | 54.37% | Morocco Arab | 99.34% |
| Scotland | 45.85% | Paraguay | 4.90% | Morocco Caucasoid | 99.24% |
| Scotland Caucasoid | 45.85% | Paraguay Amerindian | 4.90% | Nauru | 23.76% |
| Senegal | 98.04% | Peru | 64.73% | Nauru Micronesian | 23.76% |
| Senegal Black | 98.04% | Peru Amerindian | 64.73% | Netherlands | 60.75% |
| Serbia | 45.09% | Philippines | 23.42% | Netherlands Caucasoid | 60.75% |
| Serbia Caucasoid | 45.09% | Philippines Austronesian | 23.42% | New Caledonia | 99.58% |
| Singapore | 95.47% | Poland | 58.52% | New Caledonia Melanesian | 99.58% |
| Singapore Austronesian | 93.95% | Poland Caucasoid | 58.52% | New Zealand | 47.99% |
| Singapore Oriental | 96.45% | Portugal | 52.81% | New Zealand Polynesian | 47.99% |
| South Africa | 96.29% | Portugal Caucasoid | 52.81% | Niue | 38.38% |
| South Africa Black | 84.70% | Russia | 62.19% | Niue Polynesian | 38.38% |
| South Africa Other | 99.33% | Russia Caucasoid | 64.05% | North Africa | 98.30% |
| South America | 99.20% | Russia Other | 62.57% | North America | 99.91% |
| South Asia | 99.60% | Russia Siberian | 62.98% | Northeast Asia | 97.97% |
| Southeast Asia | 98.97% | Rwanda | 44.05% | Norway | 57.75% |
| Southwest Asia | 94.18% | Rwanda Black | 44.05% | Norway Caucasoid | 57.75% |
| Spain | 98.63% | Samoa | 62.78% | Oceania | 98.12% |
| Spain Caucasoid | 98.63% | Samoa Polynesian | 62.78% | Oman | 98.07% |
| Sri Lanka | 60.18% | Sao Tome and Principe | 46.30% | Oman Arab | 98.07% |
| Sri Lanka Asian | 60.18% | Sao Tome and Principe Black | 46.30% | Pakistan | 94.40% |
| Sudan | 96.57% | Saudi Arabia | 38.09% | Pakistan Asian | 93.16% |
| Sudan Arab | 85.12% | Saudi Arabia Arab | 38.09% | Pakistan Mixed | 96.57% |
| Sudan Black | 2.19% | Scotland | 48.88% | Papua New Guinea | 99.58% |
| Sudan Mixed | 92.33% | Scotland Caucasoid | 48.88% | Papua New Guinea Melanesian | 99.58% |
| Sweden | 96.33% | Senegal | 26.56% | Paraguay | 4.90% |
| Sweden Caucasoid | 96.33% | Senegal Black | 26.56% | Paraguay Amerindian | 4.90% |
| Switzerland | 72.65% | Singapore | 23.96% | Peru | 100.00% |
| Switzerland Caucasoid | 72.65% | Singapore Austronesian | 23.96% | Peru Amerindian | 100.00% |
| Taiwan | 98.53% | Slovakia | 0.00% | Peru Mestizo | 1.99% |
| Taiwan Oriental | 98.53% | Slovakia Caucasoid | 0.00% | Philippines | 98.56% |
| Thailand | 98.27% | Slovenia | 61.85% | Philippines Austronesian | 98.56% |
| Thailand Oriental | 98.27% | Slovenia Caucasoid | 61.85% | Poland | 99.62% |
| Tunisia | 97.63% | South Africa | 1.79% | Poland Caucasoid | 99.62% |
| Tunisia Arab | 97.63% | South Africa Black | 1.79% | Portugal | 99.33% |
| Turkey | 60.53% | South America | 54.19% | Portugal Caucasoid | 99.33% |
| Turkey Caucasoid | 60.53% | South Asia | 36.20% | Romania | 96.93% |
| Uganda | 96.90% | Southeast Asia | 39.53% | Romania Caucasoid | 96.93% |
| Uganda Black | 96.90% | Southwest Asia | 41.52% | Russia | 99.60% |
| United Arab Emirates | 5.72% | Spain | 51.81% | Russia Caucasoid | 89.52% |
| United Arab Emirates Arab | 5.72% | Spain Caucasoid | 51.86% | Russia Mixed | 66.31% |
| United Kingdom | 80.29% | Sudan | 49.02% | Russia Other | 99.80% |
| United Kingdom Caucasoid | 80.29% | Sudan Mixed | 49.02% | Russia Siberian | 99.61% |
| United States | 99.76% | Sweden | 83.95% | Rwanda | 68.61% |
| United States Amerindian | 99.42% | Sweden Caucasoid | 83.95% | Rwanda Black | 68.61% |
| United States Asian | 97.85% | Taiwan | 51.72% | Samoa | 62.78% |
| United States Black | 99.46% | Taiwan Oriental | 51.72% | Samoa Polynesian | 62.78% |
| United States Caucasoid | 99.44% | Thailand | 38.32% | Sao Tome and Principe | 95.41% |
| United States Hispanic | 98.64% | Thailand Oriental | 38.32% | Sao Tome and Principe Black | 95.41% |
| United States Mestizo | 98.89% | Tokelau | 20.79% | Saudi Arabia | 99.05% |
| United States Polynesian | 99.57% | Tokelau Polynesian | 20.79% | Saudi Arabia Arab | 99.05% |
| Venezuela | 91.27% | Tonga | 61.56% | Scotland | 72.32% |
| Venezuela Amerindian | 90.80% | Tonga Polynesian | 61.56% | Scotland Caucasoid | 72.32% |
| Venezuela Caucasoid | 12.51% | Tunisia | 53.49% | Senegal | 98.56% |
| Venezuela Mestizo | 11.55% | Tunisia Arab | 53.11% | Senegal Black | 98.56% |
| Vietnam | 96.39% | Tunisia Berber | 57.48% | Serbia | 45.09% |
| Vietnam Oriental | 96.39% | Turkey | 56.18% | Serbia Caucasoid | 45.09% |
| Wales | 1.00% | Turkey Caucasoid | 56.18% | Singapore | 96.55% |
| Wales Caucasoid | 1.00% | Ukraine | 46.48% | Singapore Austronesian | 95.40% |
| West Africa | 97.16% | Ukraine Caucasoid | 46.48% | Singapore Oriental | 96.45% |
| West Indies | 96.43% | United States | 60.50% | Slovakia | 0.00% |
| World | 99.55% | United States Amerindian | 53.24% | Slovakia Caucasoid | 0.00% |
| Zambia | 96.38% | United States Asian | 49.73% | Slovenia | 61.85% |
| Zambia Black | 96.38% | United States Austronesian | 41.94% | Slovenia Caucasoid | 61.85% |
| Zimbabwe | 95.94% | United States Black | 49.76% | South Africa | 96.35% |
| Zimbabwe Black | 95.94% | United States Caucasoid | 59.36% | South Africa Black | 84.98% |
|  |  | United States Hispanic | 60.78% | South Africa Other | 99.33% |
|  |  | United States Mestizo | 63.42% | South America | 99.63% |
|  |  | United States Polynesian | 49.13% | South Asia | 99.75% |
|  |  | Vietnam | 29.71% | Southeast Asia | 99.38% |
|  |  | Vietnam Oriental | 29.71% | Southwest Asia | 96.60% |
|  |  | West Africa | 50.22% | Spain | 99.34% |
|  |  | West Indies | 50.58% | Spain Caucasoid | 99.34% |
|  |  | World | 56.36% | Sri Lanka | 60.18% |
|  |  | Zimbabwe | 54.57% | Sri Lanka Asian | 60.18% |
|  |  | Zimbabwe Black | 54.57% | Sudan | 98.25% |
|  |  |  |  | Sudan Arab | 85.12% |
|  |  |  |  | Sudan Black | 2.19% |
|  |  |  |  | Sudan Mixed | 96.09% |
|  |  |  |  | Sweden | 99.41% |
|  |  |  |  | Sweden Caucasoid | 99.41% |
|  |  |  |  | Switzerland | 72.65% |
|  |  |  |  | Switzerland Caucasoid | 72.65% |
|  |  |  |  | Taiwan | 99.29% |
|  |  |  |  | Taiwan Oriental | 99.29% |
|  |  |  |  | Thailand | 98.93% |
|  |  |  |  | Thailand Oriental | 98.93% |
|  |  |  |  | Tokelau | 20.79% |
|  |  |  |  | Tokelau Polynesian | 20.79% |
|  |  |  |  | Tonga | 61.56% |
|  |  |  |  | Tonga Polynesian | 61.56% |
|  |  |  |  | Tunisia | 98.90% |
|  |  |  |  | Tunisia Arab | 98.89% |
|  |  |  |  | Tunisia Berber | 57.48% |
|  |  |  |  | Turkey | 82.70% |
|  |  |  |  | Turkey Caucasoid | 82.70% |
|  |  |  |  | Uganda | 96.90% |
|  |  |  |  | Uganda Black | 96.90% |
|  |  |  |  | Ukraine | 46.48% |
|  |  |  |  | Ukraine Caucasoid | 46.48% |
|  |  |  |  | United Arab Emirates | 5.72% |
|  |  |  |  | United Arab Emirates Arab | 5.72% |
|  |  |  |  | United Kingdom | 80.29% |
|  |  |  |  | United Kingdom Caucasoid | 80.29% |
|  |  |  |  | United States | 99.91% |
|  |  |  |  | United States Amerindian | 99.73% |
|  |  |  |  | United States Asian | 98.92% |
|  |  |  |  | United States Austronesian | 41.94% |
|  |  |  |  | United States Black | 99.73% |
|  |  |  |  | United States Caucasoid | 99.77% |
|  |  |  |  | United States Hispanic | 99.47% |
|  |  |  |  | United States Mestizo | 99.59% |
|  |  |  |  | United States Polynesian | 99.78% |
|  |  |  |  | Venezuela | 91.27% |
|  |  |  |  | Venezuela Amerindian | 90.80% |
|  |  |  |  | Venezuela Caucasoid | 12.51% |
|  |  |  |  | Venezuela Mestizo | 11.55% |
|  |  |  |  | Vietnam | 97.46% |
|  |  |  |  | Vietnam Oriental | 97.46% |
|  |  |  |  | Wales | 1.00% |
|  |  |  |  | Wales Caucasoid | 1.00% |
|  |  |  |  | West Africa | 98.59% |
|  |  |  |  | West Indies | 98.24% |
|  |  |  |  | World | 99.80% |
|  |  |  |  | Zambia | 96.38% |
|  |  |  |  | Zambia Black | 96.38% |
|  |  |  |  | Zimbabwe | 98.15% |
|  |  |  |  | Zimbabwe Black | 98.15% |

**Table S5:** Potential CTL epitopes with their respective MHC-I binding alleles (Consensus: PR≤2)

| **CTL Epitopes** | **MHC-I binding Alleles** | **Total** |
| --- | --- | --- |
| AAAVPTTAL | HLA-C*03:03,HLA-B*14:02,HLA-B*07:02,HLA-B*46:01,HLA-B*15:02,HLA-B*39:01,HLA-B*15:17,HLA-B*83:01,HLA-B*48:01,HLA-A*32:01,HLA-B*35:01,HLA-A*69:01 | 12 |
| AADEVSAAI | HLA-C*05:01,HLA-C*12:03,HLA-B*08:03,HLA-C*08:02,HLA-B*39:01,HLA-A*69:01,HLA-A*02:06,HLA-B*83:01,HLA-B*51:01,HLA-B*46:01 | 10 |
| AANGAVGSR | HLA-A*31:01,HLA-B*83:01,HLA-A*26:03,HLA-A*68:01,HLA-A*11:01 | 5 |
| ALAAGGASY | HLA-B*15:01,HLA-A*30:02,HLA-A*29:02,HLA-A*80:01,HLA-A*26:01,HLA-B*15:02,HLA-A*03:01,HLA-A*26:02,HLA-B*15:03,HLA-A*01:01,HLA-B*44:02,HLA-B*15:17,HLA-C*03:03,HLA-A*26:03,HLA-B*46:01 | 15 |
| APNGGAGGL | HLA-B*83:01,HLA-B*07:02,HLA-B*42:01,HLA-B*39:01,HLA-B*35:03,HLA-A*26:03,HLA-B*15:09 | 7 |
| DEVSAAIAA | HLA-B*18:01,HLA-B*45:01,HLA-B*44:03,HLA-B*44:02,HLA-B*73:01,HLA-B*40:01 | 6 |
| EVSAAIAAL | HLA-A*25:01,HLA-A*26:03,HLA-A*26:01,HLA-A*68:02,HLA-A*26:02,HLA-A*69:01,HLA-C*03:03,HLA-A*66:01,HLA-B*39:01,HLA-A*68:23,HLA-B*48:01,HLA-B*35:01,HLA-A*02:19,HLA-B*35:03,HLA-B*07:02 | 15 |
| GGDGGHGAY | HLA-A*30:02,HLA-A*01:01,HLA-A*29:02,HLA-C*05:01,HLA-A*80:01,HLA-C*08:02,HLA-B*15:02 | 7 |
| GQGGGAGTL | HLA-B*48:01,HLA-B*40:01,HLA-B*38:01,HLA-B*39:01,HLA-A*02:12 | 5 |
| LWPAAADEV | HLA-A*24:03,HLA-C*04:01,HLA-A*02:17,HLA-A*02:50,HLA-B*51:01 | 5 |
| LYGNGGNGY | HLA-A*29:02,HLA-C*14:02,HLA-A*30:02,HLA-A*24:03,HLA-A*32:07,HLA-A*80:01,HLA-B*15:03 | 7 |
| SSDAANGAV | HLA-A*01:01,HLA-C*08:02,HLA-A*69:01,HLA-C*05:01,HLA-C*15:02,HLA-C*12:03,HLA-B*39:01 | 7 |
| VSAAIAALF | HLA-B*58:01,HLA-B*15:17,HLA-B*57:01,HLA-A*24:02,HLA-A*23:01,HLA-B*58:02,HLA-A*29:02,HLA-A*01:01,HLA-B*15:01,HLA-E*01:01,HLA-A*30:02,HLA-A*26:02,HLA-A*32:01,HLA-A*26:01,HLA-B*53:01,HLA-C*04:01 | 16 |
| YLSVAPDEL | HLA-A*02:02,HLA-A*02:19,HLA-A*02:50,HLA-A*02:12,HLA-A*02:16,HLA-A*02:01,HLA-A*02:03,HLA-C*03:03,HLA-B*08:02,HLA-A*02:11,HLA-B*15:09,HLA-B*08:03,HLA-B*15:02,HLA-C*06:02,HLA-B*39:01,HLA-B*46:01,HLA-C*07:02 | 17 |

**Table S6:** Potential HTL epitopes with their respective MHC-II binding alleles (Consensus: PR≤2)

| **HTL Epitopes** | **MHC-II binding Alleles** | **Total** |
| --- | --- | --- |
| AAAATELRGVGSIVS | HLA-DRB1*04:23,HLA-DRB1*04:08,HLA-DRB1*13:11,HLA-DRB1*11:04,HLA-DRB1*11:06,HLA-DRB1*04:10,HLA-DRB1*04:02,HLA-DRB5*01:05,HLA-DRB1*11:28,HLA-DRB1*13:05,HLA-DRB1*11:07,HLA-DRB1*01:02,HLA-DRB1*15:06,HLA-DRB1*03:08,HLA-DRB1*03:06,HLA-DRB1*03:07,HLA-DRB1*13:07,HLA-DRB1*13:21,HLA-DRB1*08:04,HLA-DRB1*15:02,HLA-DRB1*03:05,HLA-DRB1*07:03,HLA-DRB1*04:26 | 23 |
| AAATELRGVGSIVST | HLA-DRB1*04:23,HLA-DRB1*04:08,HLA-DRB1*13:11,HLA-DRB1*11:04,HLA-DRB1*11:06,HLA-DRB1*04:10,HLA-DRB1*04:02,HLA-DRB5*01:05,HLA-DRB1*11:28,HLA-DRB1*13:05,HLA-DRB1*11:07,HLA-DRB1*01:02,HLA-DRB1*15:06,HLA-DRB1*03:08,HLA-DRB1*03:06,HLA-DRB1*03:07,HLA-DRB1*13:07,HLA-DRB1*13:21,HLA-DRB1*08:04,HLA-DRB1*15:02,HLA-DRB1*03:05,HLA-DRB1*07:03,HLA-DRB1*04:26 | 23 |
| AATELRGVGSIVSTV | HLA-DRB1*04:23,HLA-DRB1*04:08,HLA-DRB1*13:11,HLA-DRB1*11:04,HLA-DRB1*11:06,HLA-DRB1*04:10,HLA-DRB1*04:02,HLA-DRB5*01:05,HLA-DRB1*11:28,HLA-DRB1*13:05,HLA-DRB1*11:07,HLA-DRB1*01:02,HLA-DRB1*15:06,HLA-DRB1*03:08,HLA-DRB1*03:06,HLA-DRB1*03:07,HLA-DRB1*13:07,HLA-DRB1*13:21,HLA-DRB1*08:04,HLA-DRB1*15:02,HLA-DRB1*03:05,HLA-DRB1*07:03,HLA-DRB1*04:26,HLA-DQA1*05:01/DQB1*03:01 | 24 |
| AAYHDQFVRALAAGG | HLA-DRB1*09:01,HLA-DRB1*01:02,HLA-DRB1*13:07,HLA-DRB1*11:20,HLA-DRB1*11:28,HLA-DRB1*08:01,HLA-DRB1*13:05,HLA-DRB1*04:21,HLA-DRB1*13:21,HLA-DRB5*01:05,HLA-DRB5*01:01,HLA-DRB1*03:09,HLA-DRB1*04:05,HLA-DRB1*04:08 | 14 |
| AQQYQSLSVQVAAYH | HLA-DRB1*13:07,HLA-DRB1*04:08,HLA-DRB1*04:26,HLA-DRB1*13:23,HLA-DRB1*11:14,HLA-DRB1*08:13,HLA-DRB1*11:01,HLA-DRB1*03:05,HLA-DRB1*04:21,HLA-DRB1*09:01 | 10 |
| ASPLEQLLDLINLPS | HLA-DRB1*04:04,HLA-DRB1*04:23,HLA-DRB1*04:08,HLA-DRB1*13:11,HLA-DRB1*11:04,HLA-DRB1*11:06,HLA-DRB1*04:01,HLA-DRB1*04:26 | 8 |
| ATELRGVGSIVSTVN | HLA-DRB1*04:23,HLA-DRB1*04:08,HLA-DRB1*13:11,HLA-DRB1*11:04,HLA-DRB1*11:06,HLA-DRB1*04:10,HLA-DRB1*04:02,HLA-DRB5*01:05,HLA-DRB1*11:28,HLA-DRB1*13:05,HLA-DRB1*11:07,HLA-DRB1*01:02,HLA-DRB1*15:06,HLA-DRB1*03:08,HLA-DRB1*03:06,HLA-DRB1*03:07,HLA-DRB1*04:04,HLA-DRB1*13:07,HLA-DRB1*13:21,HLA-DRB1*08:04,HLA-DRB1*15:02,HLA-DRB1*03:05,HLA-DRB1*07:03,HLA-DQA1*05:01/DQB1*03:01,  HLA-DRB1*04:26 | 25 |
| AYHDQFVRALAAGGA | HLA-DRB1*09:01,HLA-DRB1*13:07,HLA-DRB1*01:02,HLA-DRB1*11:20,HLA-DRB1*13:11,HLA-DRB1*11:04,HLA-DRB1*11:06,HLA-DRB1*11:28,HLA-DRB1*08:01,HLA-DRB1*13:05,HLA-DRB1*08:04,HLA-DRB1*04:21,HLA-DRB1*13:21,HLA-DRB5*01:05,HLA-DRB5*01:01,HLA-DRB1*03:09,HLA-DRB1*04:05,HLA-DRB1*11:01,HLA-DRB1*04:08 | 19 |
| DEVSAAIAALFSSHA | HLA-DQA1*04:01/DQB1*04:02,HLA-DRB1*09:01,HLA-DRB1*04:04,HLA-DRB1*04:23,HLA-DRB1*15:01 | 5 |
| DQFVRALAAGGASYS | HLA-DRB1*09:01,HLA-DRB1*13:07,HLA-DRB1*01:02,HLA-DRB1*11:20,HLA-DRB1*13:11,HLA-DRB1*11:04,HLA-DRB1*11:06,HLA-DRB5*01:01,HLA-DRB1*11:28,HLA-DRB1*08:01,HLA-DRB1*13:05,HLA-DRB1*01:01,HLA-DRB1*08:04,HLA-DRB1*04:21,HLA-DRB1*13:21,HLA-DRB5*01:05,HLA-DRB1*03:09,HLA-DRB1*11:01,HLA-DRB1*04:05,HLA-DRB1*04:08 | 20 |
| ELRGVGSIVSTVNAA | HLA-DRB1*04:23,HLA-DRB1*04:08,HLA-DRB1*13:11,HLA-DRB1*11:04,HLA-DRB1*11:06,HLA-DRB1*04:10,HLA-DRB1*04:02,HLA-DRB5*01:05,HLA-DRB1*11:28,HLA-DRB1*13:05,HLA-DRB1*11:07,HLA-DRB1*01:02,HLA-DRB1*15:06,HLA-DRB1*03:08,HLA-DRB1*03:06,HLA-DRB1*03:07,HLA-DRB1*13:07,HLA-DRB1*13:21,HLA-DRB1*08:04,HLA-DRB1*15:02,HLA-DRB1*03:05,HLA-DRB1*07:03,HLA-DRB1*04:26,HLA-DQA1*05:01/DQB1*03:01,  HLA-DRB1*04:04 | 25 |
| EQLLDLINLPSQTLL | HLA-DRB1*01:02,HLA-DRB1*04:04,HLA-DRB1*04:23,HLA-DRB1*01:01,HLA-DRB1*04:10,HLA-DRB1*04:08,HLA-DRB1*07:03,HLA-DRB1*13:11,HLA-DRB1*11:04,HLA-DRB1*11:06,HLA-DRB1*04:01,HLA-DRB1*04:26 | 12 |
| EVSAAIAALFSSHAQ | HLA-DQA1*04:01/DQB1*04:02,HLA-DRB1*09:01,HLA-DRB1*08:02,HLA-DRB1*04:23,HLA-DRB1*15:01,HLA-DRB1*04:04 | 6 |
| FVRALAAGGASYSLA | HLA-DRB1*13:07,HLA-DRB1*01:02,HLA-DRB1*11:20,HLA-DRB1*09:01,HLA-DRB1*13:11,HLA-DRB1*11:04,HLA-DRB1*11:06,HLA-DRB1*11:28,HLA-DRB1*08:01,HLA-DRB1*13:05,HLA-DRB1*08:04,HLA-DRB1*04:21,HLA-DRB1*13:21,HLA-DRB5*01:05,HLA-DRB5*01:01,HLA-DRB1*03:09,HLA-DRB1*04:08 | 16 |
| GGGGSQVGSDGVGGA | HLA-DRB1*11:01,HLA-DRB1*03:05,HLA-DRB1*03:08,HLA-DRB1*03:06,HLA-DRB1*03:07,HLA-DRB1*03:09 | 6 |
| GGGSQVGSDGVGGAG | HLA-DRB1*11:07,HLA-DRB1*03:05,HLA-DRB1*03:08,HLA-DRB1*03:06,HLA-DRB1*03:07,HLA-DRB1*03:09 | 6 |
| GGSQVGSDGVGGAGG | HLA-DRB1*11:07,HLA-DRB1*03:05,HLA-DRB1*03:08,HLA-DRB1*03:06,HLA-DRB1*03:07,HLA-DRB1*03:09 | 6 |
| GSIVSTVNAAAAVPT | HLA-DRB1*09:01,HLA-DRB1*04:04,HLA-DRB1*04:23,HLA-DRB1*04:21,HLA-DRB1*01:02,HLA-DRB1*04:08 | 6 |
| GSQVGSDGVGGAGGA | HLA-DRB1*11:07,HLA-DRB1*03:05,HLA-DRB1*03:08,HLA-DRB1*03:06,HLA-DRB1*03:07,HLA-DRB1*03:09 | 6 |
| GVGSIVSTVNAAAAV | HLA-DRB1*08:02,HLA-DRB1*04:01,HLA-DRB1*04:26,HLA-DRB1*04:02,HLA-DRB1*04:10,HLA-DRB1*04:04,HLA-DRB1*04:23,HLA-DRB1*04:21,HLA-DRB1*09:01,HLA-DRB1*04:08 | 10 |
| HAQQYQSLSVQVAAY | HLA-DRB1*13:07,HLA-DRB1*04:08,HLA-DRB1*04:26,HLA-DRB1*13:23,HLA-DRB1*11:14,HLA-DRB1*08:13,HLA-DRB1*11:01,HLA-DRB1*03:05,HLA-DRB1*04:21,HLA-DRB1*09:01 | 10 |
| HDQFVRALAAGGASY | HLA-DRB1*09:01,HLA-DRB5*01:01,HLA-DRB1*13:07,HLA-DRB1*01:02,HLA-DRB1*11:01,HLA-DRB1*11:20,HLA-DRB1*13:11,HLA-DRB1*11:04,HLA-DRB1*11:06,HLA-DRB1*11:28,HLA-DRB1*08:01,HLA-DRB1*13:05,HLA-DRB1*01:01,HLA-DRB1*08:04,HLA-DRB1*04:21,HLA-DRB1*13:21,HLA-DRB5*01:05,HLA-DRB1*03:09,HLA-DRB1*04:05,HLA-DRB1*04:08 | 20 |
| IVSTVNAAAAVPTTA | HLA-DRB1*04:26, HLA-DRB1*08:02,HLA-DRB1*04:02,HLA-DRB1*04:23,HLA-DRB1*04:21,HLA-DRB1*09:01,HLA-DRB1*04:04,HLA-DQA1*01:02/DQB1*06:02,  HLA-DRB1*01:02,HLA-DRB1*04:08 | 10 |
| LDLINLPSQTLLGRP | HLA-DRB1*01:02,HLA-DRB1*01:01,HLA-DRB1*04:10,HLA-DRB1*04:23,HLA-DRB1*07:03,HLA-DRB1*04:04 | 6 |
| LEQLLDLINLPSQTL | HLA-DRB1*01:02,HLA-DRB1*04:04,HLA-DRB1*04:23,HLA-DRB1*04:10,HLA-DRB1*04:08,HLA-DRB1*07:03,HLA-DRB1*13:11,HLA-DRB1*11:04,HLA-DRB1*11:06,HLA-DRB1*04:01,HLA-DRB1*04:26 | 11 |
| LLDLINLPSQTLLGR | HLA-DRB1*01:02,HLA-DRB1*04:04,HLA-DRB1*04:23,HLA-DRB1*01:01,HLA-DRB1*04:10,HLA-DRB1*04:08,HLA-DRB1*07:03,HLA-DRB1*13:11,HLA-DRB1*11:04,HLA-DRB1*11:06,HLA-DRB1*04:01,HLA-DRB1*04:26 | 12 |
| LRGVGSIVSTVNAAA | HLA-DRB1*04:23,HLA-DRB1*04:08,HLA-DRB1*13:11,HLA-DRB1*11:04,HLA-DRB1*11:06,HLA-DRB1*04:10,HLA-DRB1*04:02,HLA-DRB1*08:02,HLA-DRB1*04:01,HLA-DRB1*04:26,HLA-DRB5*01:05,HLA-DRB1*11:28,HLA-DRB1*13:05,HLA-DRB1*11:07,HLA-DRB1*01:02,HLA-DRB1*15:06,HLA-DRB1*04:04,HLA-DRB1*03:08,HLA-DRB1*03:06,HLA-DRB1*03:07,HLA-DRB1*13:07,HLA-DRB1*04:21,HLA-DRB1*13:21,HLA-DRB1*08:04,HLA-DRB1*15:02,HLA-DRB1*03:05,HLA-DRB1*07:03 | 27 |
| LSVAPDELAAAATEL | HLA-DQA1*04:01/DQB1*04:02,HLA-DQA1*03:01/DQB1*03:02,HLA-DRB1*03:08  HLA-DRB1*03:06,HLA-DRB1*03:07,HLA-DRB1*11:07 | 6 |
| MWYLSVAPDELAAAA | HLA-DQA1*05:01/DQB1*02:01,HLA-DQA1*04:01/DQB1*04:02,HLA-DRB1*08:06,  HLA-DRB1*04:10,HLA-DRB1*03:08,HLA-DRB1*03:06,HLA-DRB1*03:07,HLA-DRB1*13:21,HLA-DRB1*13:04,HLA-DRB1*11:07,HLA-DRB1*03:09,HLA-DRB1*08:01,HLA-DRB1*04:05 | 13 |
| PLEQLLDLINLPSQT | HLA-DRB1*04:04,HLA-DRB1*04:23,HLA-DRB1*04:08,HLA-DRB1*13:11,HLA-DRB1*11:04,HLA-DRB1*11:06,HLA-DRB1*04:01,HLA-DRB1*04:26 | 8 |
| QFVRALAAGGASYSL | HLA-DRB1*09:01,HLA-DRB1*13:07,HLA-DRB1*01:02,HLA-DRB1*11:20,HLA-DRB1*13:11,HLA-DRB1*11:04,HLA-DRB1*11:06,HLA-DRB1*11:28,HLA-DRB1*08:01,HLA-DRB1*13:05,HLA-DRB5*01:01,HLA-DRB1*08:04,HLA-DRB1*04:21,HLA-DRB1*13:21,HLA-DRB5*01:05,HLA-DRB1*03:09,HLA-DRB1*04:08 | 17 |
| QLLDLINLPSQTLLG | HLA-DRB1*01:02,HLA-DRB1*04:04,HLA-DRB1*04:23,HLA-DRB1*01:01,HLA-DRB1*04:10,HLA-DRB1*04:08,HLA-DRB1*07:03,HLA-DRB1*13:11,HLA-DRB1*11:04,HLA-DRB1*11:06,HLA-DRB1*04:01,HLA-DRB1*04:26 | 12 |
| QQYQSLSVQVAAYHD | HLA-DRB1*13:07,HLA-DRB1*04:08,HLA-DRB1*04:26,HLA-DRB1*13:23,HLA-DRB1*11:14,HLA-DRB1*08:13,HLA-DRB1*11:01,HLA-DRB1*03:05,HLA-DRB1*04:21 | 9 |
| QVGSDGVGGAGGAGG | HLA-DRB1*11:07,HLA-DRB1*03:05,HLA-DRB1*03:08,HLA-DRB1*03:06,HLA-DRB1*03:07,HLA-DRB1*03:09 | 6 |
| QYQSLSVQVAAYHDQ | HLA-DRB1*13:07,HLA-DRB1*04:08,HLA-DRB1*04:26,HLA-DRB1*13:23,HLA-DRB1*11:14,HLA-DRB1*08:13,HLA-DRB1*03:05,HLA-DRB1*04:21 | 8 |
| RGVGSIVSTVNAAAA | HLA-DRB1*08:02,HLA-DRB1*04:01,HLA-DRB1*04:26,HLA-DRB1*04:02,HLA-DRB1*04:10,HLA-DRB1*04:04,HLA-DRB1*04:23,HLA-DRB1*04:21,HLA-DRB1*09:01,HLA-DRB1*04:08 | 10 |
| SAAIAALFSSHAQQY | HLA-DRB1*09:01,HLA-DRB1*08:02,HLA-DRB1*04:23,HLA-DRB1*04:01,HLA-DRB1*15:01,HLA-DRB1*04:04 | 6 |
| SHAQQYQSLSVQVAA | HLA-DRB1*13:07,HLA-DRB1*04:08,HLA-DRB1*04:26,HLA-DRB1*13:23,HLA-DRB1*11:14,HLA-DRB1*08:13,HLA-DRB1*11:01,HLA-DRB1*03:05,HLA-DRB1*04:21 | 9 |
| SIVSTVNAAAAVPTT | HLA-DRB1*08:02,HLA-DRB1*04:26,HLA-DRB1*04:02,HLA-DRB1*09:01,HLA-DRB1*04:04,HLA-DRB1*04:23,HLA-DRB1*04:21,HLA-DQA1*01:02/DQB1*06:02, HLA-DRB1*04:01,HLA-DRB1*01:02,HLA-DRB1*04:08 | 11 |
| SPLEQLLDLINLPSQ | HLA-DRB1*04:04,HLA-DRB1*04:23,HLA-DRB1*04:08,HLA-DRB1*13:11,HLA-DRB1*11:04,HLA-DRB1*11:06,HLA-DRB1*04:01,HLA-DRB1*04:26 | 8 |
| SQVGSDGVGGAGGAG | HLA-DRB1*11:07,HLA-DRB1*03:05,HLA-DRB1*03:08,HLA-DRB1*03:06,HLA-DRB1*03:07,HLA-DRB1*03:09 | 6 |
| SSHAQQYQSLSVQVA | HLA-DRB1*13:07,HLA-DRB1*04:08,HLA-DRB1*04:26,HLA-DRB1*13:23,HLA-DRB1*11:14,HLA-DRB1*08:13,HLA-DRB1*11:01,HLA-DRB1*03:05,HLA-DRB1*04:21 | 9 |
| SVAPDELAAAATELR | HLA-DQA1*04:01/DQB1*04:02,HLA-DQA1*03:01/DQB1*03:02,HLA-DRB1*03:08, HLA-DRB1*03:06,HLA-DRB1*03:07,HLA-DRB1*11:07 | 6 |
| TELRGVGSIVSTVNA | HLA-DRB1*04:23,HLA-DRB1*04:08,HLA-DRB1*13:11,HLA-DRB1*11:04,HLA-DRB1*11:06,HLA-DRB1*04:10,HLA-DRB1*04:02,HLA-DRB5*01:05,HLA-DRB1*11:28,HLA-DRB1*13:05,HLA-DRB1*11:07,HLA-DRB1*01:02,HLA-DRB1*15:06,HLA-DRB1*03:08,HLA-DRB1*03:06,HLA-DRB1*03:07,HLA-DRB1*04:04,HLA-DRB1*13:07,HLA-DRB1*13:21,HLA-DRB1*08:04,HLA-DRB1*15:02,HLA-DRB1*03:05,HLA-DRB1*07:03,HLA-DRB1*04:26,HLA-DQA1*05:01/DQB1*03:01 | 25 |
| VAPDELAAAATELRG | HLA-DQA1*04:01/DQB1*04:02,HLA-DQA1*03:01/DQB1*03:02,HLA-DRB1*03:08  HLA-DRB1*03:06,HLA-DRB1*03:07,HLA-DRB1*11:07 | 6 |
| VGSDGVGGAGGAGGT | HLA-DRB1*11:07,HLA-DRB1*03:05,HLA-DRB1*03:08,HLA-DRB1*03:06,HLA-DRB1*03:07,HLA-DRB1*03:09 | 6 |
| VGSIVSTVNAAAAVP | HLA-DRB1*08:02,HLA-DRB1*04:01,HLA-DRB1*04:26,HLA-DRB1*04:02,HLA-DRB1*04:10,HLA-DRB1*04:04,HLA-DRB1*04:23,HLA-DRB1*04:21,HLA-DRB1*09:01,HLA-DRB1*04:08 | 10 |
| VRALAAGGASYSLAE | HLA-DRB1*13:07,HLA-DRB1*01:02,HLA-DRB1*13:11,HLA-DRB1*11:04,HLA-DRB1*11:06,HLA-DRB1*08:04 | 6 |
| VSAAIAALFSSHAQQ | HLA-DRB1*09:01,HLA-DRB1*08:02,HLA-DRB1*04:23,HLA-DRB1*15:01,HLA-DRB1*04:04 | 5 |
| WYLSVAPDELAAAAT | HLA-DQA1*05:01/DQB1*02:01,HLA-DRB1*03:08,HLA-DRB1*03:06,HLA-DRB1*03:07,HLA-DRB1*11:07,HLA-DRB1*03:09 | 6 |
| YHDQFVRALAAGGAS | HLA-DRB1*09:01,HLA-DRB1*13:07,HLA-DRB1*01:02,HLA-DRB1*11:20,HLA-DRB1*13:11,HLA-DRB1*11:04,HLA-DRB1*11:06,HLA-DRB1*11:28,HLA-DRB1*08:01,HLA-DRB1*13:05,HLA-DRB1*11:01,HLA-DRB5*01:01,HLA-DRB1*08:04,HLA-DRB1*04:21,HLA-DRB1*13:21,HLA-DRB5*01:05,HLA-DRB1*03:09,HLA-DRB1*01:01,HLA-DRB1*04:05,HLA-DRB1*04:08 | 20 |
| YLSVAPDELAAAATE | HLA-DQA1*04:01/DQB1*04:02,HLA-DQA1*03:01/DQB1*03:02,HLA-DQA1*05:01/DQB1*02:01,HLA-DRB1*03:08,HLA-DRB1*03:06,HLA-DRB1*03:07,  HLA-DRB1*11:07 | 7 |
| YQSLSVQVAAYHDQF | HLA-DRB1*13:07,HLA-DRB1*04:08,HLA-DRB1*04:26,HLA-DRB1*13:23,HLA-DRB1*11:14,HLA-DRB1*08:13,HLA-DRB1*03:05,HLA-DRB1*04:21 | 8 |

**Table S7:** Prediction of three-dimensional structure of the chimeric vaccine protein

| **Parameters** | **Results** |
| --- | --- |
| Number of predicted domains | 3 |
| Best template | 3mh9A, p-value 1.43e-09 |
| Overall uGDT (GDT) | 254 (48) |
| Residues modeled | 520(100%) |
| Positions predicted as disordered | 63(12%) |
| Secondary structure | 30%H, 20%E, 48%C |
| Solvent access | 40%E, 26%M, 32%B |

**Table S8:** Refinement of the tertiary structure of the vaccine protein

| **Model** | **GDT-HA** | **RMSD** | **MolProbity** | **Clash Score** | **Poor Rotamers** | **Rama Favored** |
| --- | --- | --- | --- | --- | --- | --- |
| Initial | 1.0000 | 0.000 | 3.225 | 79.8 | 3.8 | 93.8 |
| MODEL 1 | 0.9505 | 0.411 | 1.999 | 14.4 | 0.9 | 95.2 |
| MODEL 2 | 0.9558 | 0.413 | 1.974 | 14.0 | 0.6 | 95.4 |
| MODEL 3 | 0.9471 | 0.429 | 2.047 | 15.6 | 1.2 | 95.8 |
| MODEL 4 | 0.9519 | 0.411 | 2.095 | 14.7 | 1.5 | 95.8 |
| MODEL 5 | 0.9433 | 0.419 | 1.991 | 14.1 | 0.6 | 95.2 |

**Table S9:** List of conformational B-cell epitopes predicted in the vaccine protein

| **SN** | **Residues** | **No.** | **Scores** |
| --- | --- | --- | --- |
| 1 | Q2, G3, M4, Q5, T6, R7, R8 | 7 | 0.99 |
| 2 | L10, F11, A12, V13, L14, A15, A16, L17, G18, T19, A20, T21, A22, L23, V24, A25, G26, C27, S28, S29, G30, T31, K32 | 23 | 0.868 |
| 3 | A350, A351, A352, A353, T354, E355, L356, G357, P358, G359, P360, G361, Y362, H363, D364, Q365, V367, R368 | 18 | 0.809 |
| 4 | R295, G296, G297, P298, G299, P300, G301, S302, H303, A304, Q305, Q306, Y307 | 13 | 0.785 |
| 5 | A283, E286, L287, A288, A289, A290, A291, T292, E293, L294, Y379, A382, A383, I384, A385, A386, L387, F388, A389, A390, Y391, A392, A393, A394, V395, P396, T397, T398, A399, L400, A401, A402, Y403, Y404, L405, S406, V407, D410 | 38 | 0.776 |
| 6 | G339, S343, D347, G417, D418, G419, A423, Y424, A425, A426, Y427, A428, A429, N430, G431, A432, V433, G434, K438, G441, G442, A443, G444, G445, T446, G447, G448, A449, G450, K451, K452, G453, G454, A455, G456, G457, A458, G459, G460, T461, G462, G463, A464, K465, K466, G467, G468, N469, G470, G471, R472, G473, G474, D475, G476, I477, K479, K480, G481, N482, G483, G484, R485, G486, G487, D488, G489, I490, R491, G492, K493, K494, G495, N496, G497, G498, N499, G500, Y501, D502, N503, S504, A505, S506, K507, K508, L509, L510, Y511, G512, N513, G514, G515, N516, G517, Y518, D519, N520 | 98 | 0.687 |
| 7 | T116, P117, N118, K119 | 4 | 0.665 |
| 8 | D33, S34, G35, G36, P37, L38, P39, D40, A41, T42, P43, L44, V45, K46, Q47, T49, E50, L51, R53, N54, P138, Q150, G151, A152, K153, A154, D155, G156, R157, T159, D161, G162, Q163, T164, I166, T169, Q197, E198, N199, G200, D201, H202, Q203 | 43 | 0.661 |
| 9 | G96, G97, S98, A99 | 4 | 0.637 |
| 10 | A371, A372, A375 | 3 | 0.533 |
| 11 | K223, G225, E226, Q227, V228, S229, V230 | 7 | 0.525 |
| 12 | K171, A186, T187, E188, P189 | 5 | 0.509 |

**Table S10:** List of mutable residues in the multiepitope vaccine for disulfide engineering

| **Res1 Seq#** | **Res1 AA** | **Res2 Seq#** | **Res2 AA** | **Chi3** | **Energy** | **B-Factors** |
| --- | --- | --- | --- | --- | --- | --- |
| 38 | LEU | 154 | ALA | -103.55 | 6.99 | 0 |
| 41 | ALA | 152 | ALA | 102.97 | 4.45 | 0 |
| 49 | THR | 146 | LEU | 113.25 | 3.88 | 0 |
| 53 | ARG | 138 | PRO | 124.91 | 4.36 | 0 |
| 57 | SER | 81 | THR | 120.49 | 3.7 | 0 |
| 59 | HIS | 223 | LYS | 101.09 | 1.63 | 0 |
| 66 | GLY | 215 | ASN | -70.17 | 4.17 | 0 |
| 70 | GLY | 183 | PRO | 78.88 | 2.51 | 0 |
| 71 | LEU | 183 | PRO | 125.87 | 5.3 | 0 |
| 86 | ALA | 230 | VAL | 86.73 | 4.24 | 0 |
| 87 | ALA | 106 | VAL | 89.46 | 4.02 | 0 |
| 103 | ASP | 114 | THR | 96.74 | 5.05 | 0 |
| 111 | LEU | 126 | ALA | 99.68 | 4.33 | 0 |
| 120 | TRP | 233 | PRO | 109.56 | 3.67 | 0 |
| 137 | LYS | 141 | GLY | 103.99 | 2.2 | 0 |
| 140 | VAL | 179 | GLN | 113.16 | 5.37 | 0 |
| 148 | ASN | 173 | THR | 118.42 | 4.06 | 0 |
| 148 | ASN | 176 | ALA | -106.78 | 3.37 | 0 |
| 155 | ASP | 167 | ARG | -96.56 | 3.55 | 0 |
| 165 | THR | 195 | TRP | 119.48 | 5.98 | 0 |
| 171 | LYS | 189 | PRO | 115.12 | 4.88 | 0 |
| 174 | ALA | 186 | ALA | 94.45 | 2.04 | 0 |
| 181 | ALA | 184 | PHE | 120.82 | 3.8 | 0 |
| 195 | TRP | 206 | GLN | 81.57 | 5.32 | 0 |
| 211 | ARG | 215 | ASN | -89.11 | 4.17 | 0 |
| 223 | LYS | 226 | GLU | -94.19 | 3.01 | 0 |
| 235 | VAL | 239 | ALA | 91.62 | 4.19 | 0 |
| 245 | GLY | 258 | PRO | 107.53 | 5.82 | 0 |
| 250 | SER | 267 | GLU | -97.67 | 1.78 | 0 |
| 252 | GLY | 263 | VAL | -111.32 | 4.2 | 0 |
| 283 | ALA | 286 | GLU | -82.59 | 4.68 | 0 |
| 296 | GLY | 300 | PRO | -108.74 | 3.65 | 0 |
| 330 | SER | 336 | TYR | 118.2 | 2.57 | 0 |
| 341 | GLY | 343 | SER | 110.55 | 7.63 | 0 |
| 343 | SER | 443 | ALA | 111.35 | 4.04 | 0 |
| 347 | ASP | 446 | THR | -113.82 | 4.32 | 0 |
| 353 | ALA | 359 | GLY | 87.91 | 3.63 | 0 |
| 389 | ALA | 400 | LEU | -113.32 | 5.84 | 0 |
| 396 | PRO | 399 | ALA | 113.17 | 0.69 | 0 |
| 422 | GLY | 476 | GLY | 107.25 | 5.11 | 0 |
| 425 | ALA | 471 | GLY | -83.22 | 7.44 | 0 |
| 426 | ALA | 430 | ASN | 101.09 | 1.45 | 0 |
| 426 | ALA | 432 | ALA | -82.98 | 1.99 | 0 |
| 428 | ALA | 464 | ALA | -96.07 | 3.24 | 0 |
| 428 | ALA | 468 | GLY | -57.56 | 4.98 | 0 |
| 435 | SER | 456 | GLY | 108.53 | 3.54 | 0 |
| 443 | ALA | 447 | GLY | -64.61 | 6.7 | 0 |
| 454 | GLY | 516 | ASN | 113.55 | 3 | 0 |
| 456 | GLY | 476 | GLY | 84.56 | 3.01 | 0 |
| 457 | GLY | 475 | ASP | 80.16 | 1.49 | 0 |
| 458 | ALA | 514 | GLY | -116.39 | 7.25 | 0 |
| 460 | GLY | 472 | ARG | -112.9 | 3.26 | 0 |
| 461 | THR | 511 | TYR | 79.26 | 7.61 | 0 |
| 469 | ASN | 487 | GLY | -77.33 | 0.52 | 0 |
| 472 | ARG | 498 | GLY | -80.6 | 5.29 | 0 |
| 472 | ARG | 500 | GLY | -89.3 | 2.06 | 0 |
| 503 | ASN | 508 | LYS | 89.83 | 3.09 | 0 |
| 514 | GLY | 518 | TYR | -95.17 | 5.83 | 0 |

*The value of energy should be less than 2.2 and Chi3 should be in between −87 and +97 degree

**Table S11**: List of center and energy scores of total 26 vaccine-receptor docked complexes

| **Cluster** | **Members** | **Representative** | **Weighted Score** |
| --- | --- | --- | --- |
| 0 | 159 | Center | -1100.2 |
|  | 159 | Lowest Energy | -1372.6 |
| 1 | 84 | Center | -1129.6 |
|  | 84 | Lowest Energy | -1327.5 |
| 2 | 79 | Center | -1119.2 |
|  | 79 | Lowest Energy | -1163.5 |
| 3 | 74 | Center | -1266.6 |
|  | 74 | Lowest Energy | -1304.7 |
| 4 | 47 | Center | -1113.4 |
|  | 47 | Lowest Energy | -1267.3 |
| 5 | 43 | Center | -1198.3 |
|  | 43 | Lowest Energy | -1294.8 |
| 6 | 42 | Center | -1089.8 |
|  | 42 | Lowest Energy | -1141.4 |
| 7 | 37 | Center | -1161.1 |
|  | 37 | Lowest Energy | -1208.6 |
| 8 | 36 | Center | -1176.9 |
|  | 36 | Lowest Energy | -1307.1 |
| 9 | 35 | Center | -1082 |
|  | 35 | Lowest Energy | -1184.4 |
| 10 | 35 | Center | -1246.2 |
|  | 35 | Lowest Energy | -1246.2 |
| 11 | 34 | Center | -1066.4 |
|  | 34 | Lowest Energy | -1281.1 |
| 12 | 33 | Center | -1283.5 |
|  | 33 | Lowest Energy | -1283.5 |
| 13 | 31 | Center | -1071.1 |
|  | 31 | Lowest Energy | -1237.2 |
| 14 | 26 | Center | -1065 |
|  | 26 | Lowest Energy | -1200 |
| 15 | 22 | Center | -1083 |
|  | 22 | Lowest Energy | -1253.5 |
| 16 | 22 | Center | -1116.5 |
|  | 22 | Lowest Energy | -1260.3 |
| 17 | 21 | Center | -1151.7 |
|  | 21 | Lowest Energy | -1151.7 |
| 18 | 17 | Center | -1061.9 |
|  | 17 | Lowest Energy | -1102.5 |
| 19 | 15 | Center | -1056.4 |
|  | 15 | Lowest Energy | -1351.8 |
| 20 | 15 | Center | -1226.4 |
|  | 15 | Lowest Energy | -1226.4 |
| 21 | 11 | Center | -1099.8 |
|  | 11 | Lowest Energy | -1158.4 |
| 22 | 10 | Center | -1091.7 |
|  | 10 | Lowest Energy | -1212.7 |
| 23 | 3 | Center | -1100.6 |
|  | 3 | Lowest Energy | -1100.6 |
| 24 | 3 | Center | -1054.8 |
|  | 3 | Lowest Energy | -1072.9 |
| 25 | 1 | Center | -1063.6 |
|  | 1 | Lowest Energy | -1063.6 |

**Table S12**: MD simulation studies of the best vaccine-TLR2 complex

| **Time** | **Energy (kJ/mol)** | **Bond** | **Coulomb** | **VdW** | **RMSD** | | |
| --- | --- | --- | --- | --- | --- | --- | --- |
|  |  |  |  |  | **CA** | **Backbone** | **Heavy Atoms** |
| 0 | -19715108.45 | 412163.918 | -24453769.45 | 3873245.966 | 0.522 | 0.586 | 0.641 |
| 100 | -15351252.29 | 1361201.424 | -20624154.85 | 2963525.402 | 2.3 | 2.338 | 2.42 |
| 200 | -15239018.2 | 1299256.565 | -20362752.15 | 2896027.264 | 3.118 | 3.151 | 3.219 |
| 300 | -15171417.1 | 1264284.026 | -20206734.27 | 2858210.173 | 3.636 | 3.664 | 3.728 |
| 400 | -15132389.84 | 1238529.694 | -20117465.46 | 2839283.785 | 5.041 | 5.059 | 5.102 |
| 500 | -15118834.2 | 1231280.011 | -20065941.8 | 2815681.501 | 5.415 | 5.434 | 5.475 |
| 600 | -15100847.03 | 1231970.392 | -20045963.28 | 2813150.84 | 5.833 | 5.851 | 5.892 |
| 700 | -15088106.95 | 1218160.471 | -20021020.81 | 2815808.792 | 6.176 | 6.191 | 6.236 |
| 800 | -15081473.13 | 1220129.941 | -20002137.64 | 2802090.26 | 6.56 | 6.584 | 6.585 |
| 900 | -15068805.21 | 1219682.023 | -19994562.29 | 2807375.099 | 6.65 | 6.675 | 6.696 |
| 1000 | -15067713.43 | 1222400.669 | -19990195.09 | 2801798.693 | 6.549 | 6.573 | 6.579 |
| 1100 | -15061928.5 | 1224502.345 | -19986710.69 | 2800721.849 | 8.362 | 8.375 | 8.419 |
| 1200 | -15058029.3 | 1220879.091 | -19986167.35 | 2806331.88 | 8.308 | 8.325 | 8.327 |
| 1300 | -15064965.8 | 1216095.19 | -19988888.98 | 2810765.114 | 8.22 | 8.233 | 8.257 |
| 1400 | -15062486.83 | 1221474.272 | -19989049.49 | 2806155.747 | 7.951 | 7.967 | 7.985 |
| 1500 | -15103070.5 | 1221974.726 | -20042519.49 | 2815333.605 | 8.645 | 8.662 | 8.691 |
| 1600 | -15113099.77 | 1238062.988 | -20062099.29 | 2811016.548 | 8.063 | 8.077 | 8.116 |
| 1700 | -15121455.31 | 1234255.486 | -20088619.24 | 2828883.308 | 7.344 | 7.362 | 7.39 |
| 1800 | -15120084.03 | 1236472.419 | -20086459.54 | 2827562.426 | 8.439 | 8.449 | 8.481 |
| 1900 | -15124111.82 | 1236598.59 | -20099753.04 | 2836525.808 | 8.33 | 8.34 | 8.412 |
| 2000 | -15121408.34 | 1235476.68 | -20090079.39 | 2826686.656 | 9.623 | 9.618 | 9.715 |
| 2100 | -15122649.04 | 1242156.224 | -20091039.43 | 2821467.026 | 9.331 | 9.328 | 9.43 |
| 2200 | -15131896.12 | 1240294.617 | -20105598.7 | 2831307.768 | 9.316 | 9.315 | 9.388 |
| 2300 | -15134854.74 | 1244834.797 | -20109293.5 | 2827712.631 | 10.229 | 10.223 | 10.281 |
| 2400 | -15126396.93 | 1242332.391 | -20098257.85 | 2823550.136 | 10.432 | 10.424 | 10.491 |
| 2500 | -15133848.84 | 1238777.692 | -20108979.1 | 2831345.047 | 10.962 | 10.944 | 11.031 |
| 2600 | -15126972.86 | 1244417.302 | -20097573.15 | 2821696.09 | 10.347 | 10.336 | 10.424 |
| 2700 | -15122097.51 | 1240022.668 | -20087483.14 | 2819127.699 | 10.311 | 10.303 | 10.362 |
| 2800 | -15125592.11 | 1242106.906 | -20095915.92 | 2821717.754 | 10.207 | 10.2 | 10.278 |
| 2900 | -15125935 | 1237953.834 | -20099852.05 | 2830986.144 | 10.81 | 10.802 | 10.891 |
| 3000 | -15123824.27 | 1233475.526 | -20094602.27 | 2830450.101 | 10.77 | 10.474 | 10.796 |
| Mean | -15269666.89 | 1213265.254 | -20235278.67 | 2861791.649 | 7.671 | 7.673 | 7.734 |

**List of Supplementary Figures**

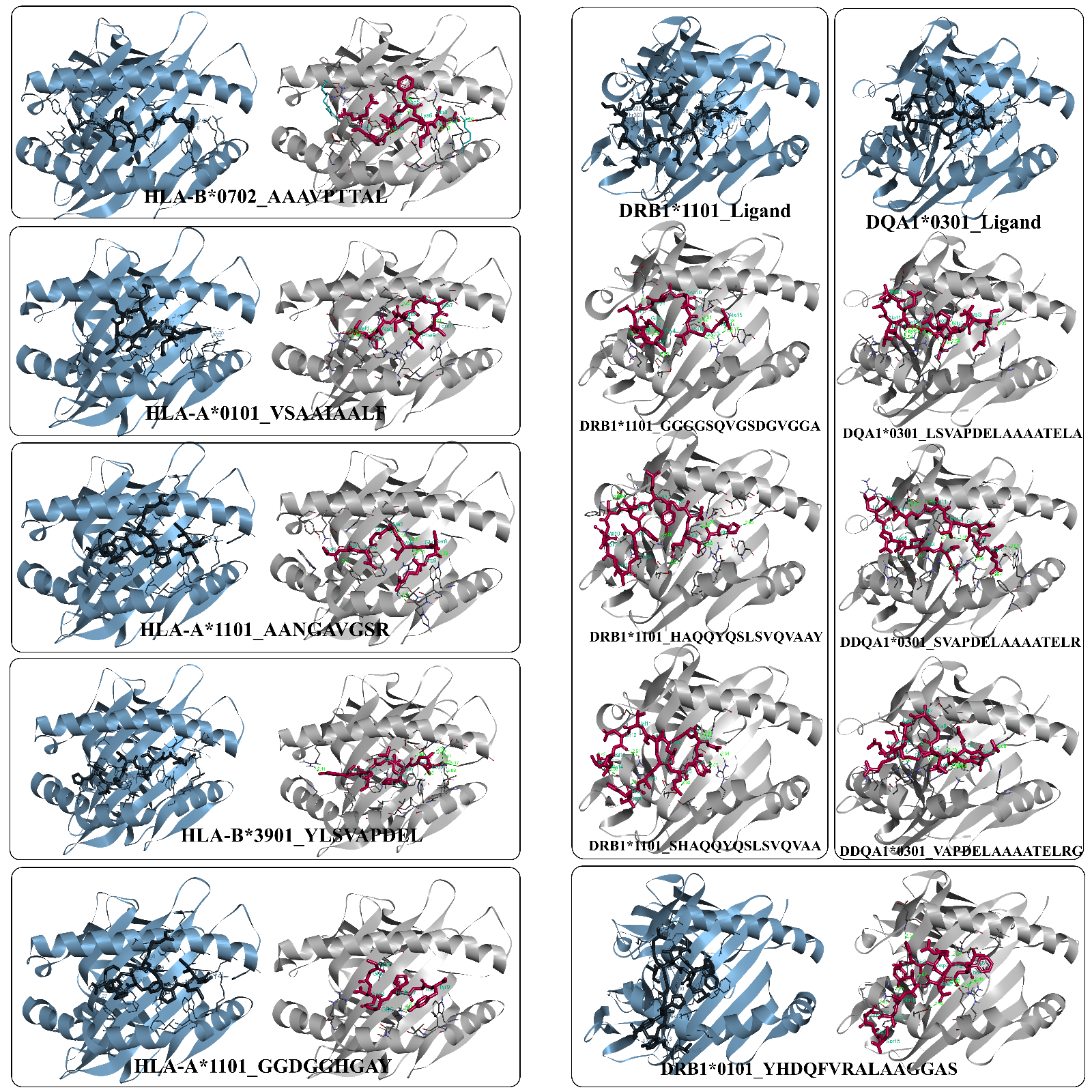

**Fig. S1:** Molecular docking and binding interactions between T-cell epitopes and their respective alleles. The docked complexes of alleles and cocrystallized ligands were considered as control and are shown in blue color.

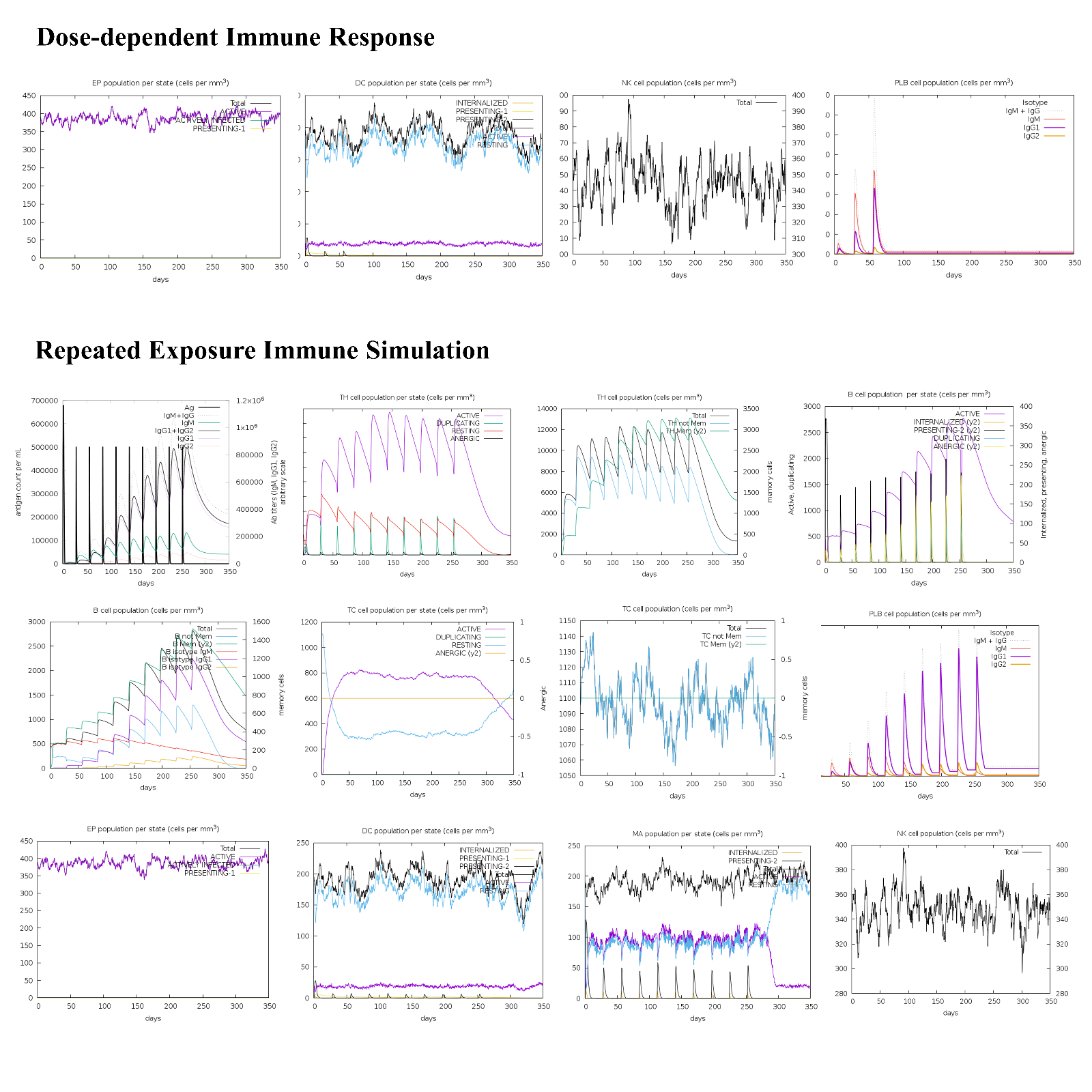

**Fig. S2:** Dose and repeated exposure induced immune responses by C-IMMSIM immune simulator
